# Unbiased profiling of multipotency landscapes reveals spatial modulators of clonal fate biases

**DOI:** 10.1101/2024.11.15.623687

**Authors:** Alek G. Erickson, Sergey Isaev, Artem Artemov, Jingyan He, Bettina Semsch, Aliia Murtazina, Jia Sun, Katrin Mangold, Anthi Chalou, Jonas Frisen, Michael Ratz, Emma R. Andersson, Peter V. Kharchenko, Igor Adameyko

## Abstract

The proportion of cell types varies systematically across the body, but it remains unclear how individual progenitor cells integrate positional information to establish patterns of cellular composition. In this study we profile the clonal landscape of the embryo, using single-cell lineage tracing of mouse embryos from neurulation until mid-gestation. To analyze the complex clonal patterns derived from highly multipotent progenitors, we developed *clone2vec*, which uses unsupervised learning to categorize individual clones into lineages based on shared transcriptional context. This revealed a body-wide gradient of clonal fate biases, in which anatomical position and clonal composition are mutually predictive. Comparison of clonal lineages revealed spatial transcription factor programs associated with dynamic cell biasing towards skeletal versus non-skeletal fates. Mosaic combinatorial perturbations targeting the Hedgehog pathway generated clones in which positional identity was mismatched with clonal composition, suggesting a potential signaling influence on somite patterning. We explore the effects of position and heterochrony on fate biases in cranial, trunk, and caudal neural crest clones. Altogether, our work demonstrates an effective practical approach for dissecting mechanisms of lineage specification.

## Main

Stochasticity during embryonic development poses a challenge for understanding how mammals robustly emerge from a single cell, through an orderly sequence of cell fate decisions and mitotic divisions (Figure 1a)^1^. Each tissue, organ, and body part, is formed from a defined set of progenitor cells that generate progressively elaborate clonal structures, varying in composition according to unique lineage histories, each contributing just a part of the overall repertoire of the body’s cell types (Figure 1b). Consequently, the classic model of cell diversification as a “tree-of-fates” represents a kind of aggregation of many individual clones at a given location. However, cell fate decisions and their underlying tree structure are thought to be highly influenced by anatomical position within the developing organism (Figure 1c). Such tree models can benefit by a massively parallel clonal analysis aimed at quantitatively establishing limits on the combinations of cell fates that emerge over time from a single progenitor cell, for each given embryonic location.

**Figure 1.**
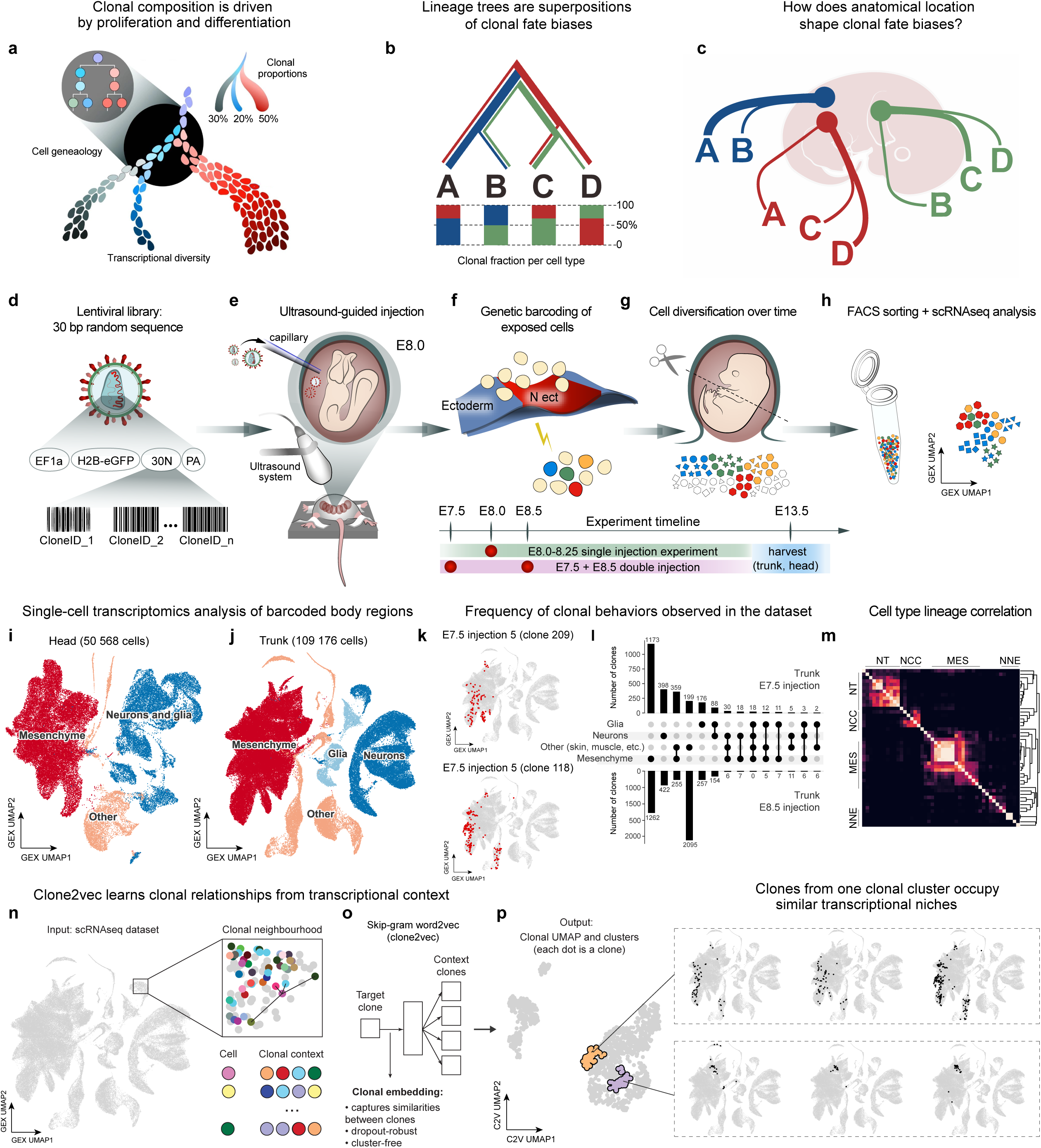
Machine learning-based profiling of clonal behavior in mouse embryos. **(a)** Scheme of a single clone developing over time via mitotic divisions and cell maturation, determining the final clonal composition. **(b)** Left, collective multipotency model, where partially overlapping fate sub-spectrums, from a population of three clones, collectively generate a fate spectrum (full repertoire of cell types). **(c)** Anatomical position of the theoretical clones shown in (b). **(d)** Clonal barcoding system. **(e)** Experimental setup. Ultrasound is used to visualize the nanoinjector and embryo while injecting lentivirus into the amniotic cavity, once or twice, between the stages of E7.5-E8.5. **(f)** Infection of the exposed virus-accessible cells in neural and non-neural ectoderm (uninfected cells beige, infected cells colorful) as well as some mesoderm. **(g)** Clones are allowed to develop until E13.5 when tissue is harvested from head and trunk body regions (each clone color-coded; each cell type shaped differently). **(h)** Barcoded cells are analyzed by single cell transcriptomics for clone reconstruction. **(i-j)** UMAP embedding of the barcoded mouse embryonic E13.5 cells from head (i) and trunk (j), color coded by the major cell types retrieved. **(k)** Examples of individual clones with partially overlapping compositions. **(l)** Upset plot of major types of trunk clones resulting from injection at E7.5 (top) versus E8.5 (bottom). **(m)** Pearson’s correlation matrix based on clone sharing pairwise across all retrieved cell types. **(n)** Schematic of clonal context neighborhood. **(o)** Skip-gram production from clonal neighborhoods using word2vec architecture. **(p)** Example of a clonal cluster within the resulting clone2vec UMAP embedding. (right) Example of similar clones constituting a clonal cluster.

Clonal lineage tracing techniques have long been used to assign progenitor populations to their derivatives^2^. Earlier approaches, such as those that rely on dye injections or multiplexed imaging of fluorescent proteins, have generally been limited in scale and throughput. Advances in multiplexed genetic tracing methods, such as Brainbow^3^, scGESTALT^4,5^, PolyLoxExpress^6^, CARLIN/DARLIN^7,8^, and LINNAEUS^9^, enabled construction of high-resolution lineage trees. These technologies promise to refine our understanding of cell differentiation mechanisms by profiling progenitor fates at scale. However, the regulatory logic and signalling interactions that establish patterns of cell type proportions remain incompletely described ^4,10^. Quantifying the spatiotemporal variation of clonal fate biases, and identifying potential biasing factors, are important steps towards elucidating the genetic mechanisms that coordinate cell destinies in vivo.

Despite recent advancements in combining highly multiplexed tracing with analysis of cellular state, a few practical challenges remain. One such difficulty is sparse coverage, where only a small fraction of cells within a tissue or organism are effectively labeled and recovered, making it difficult to reconstruct comprehensive lineage trees. More fundamentally, the measurements of the cellular state that accompany tracing information are performed at the final timepoint. Consequently, molecular differences associated with fate decisions cannot be observed directly and must be inferred under additional assumptions, such as ergodicity of the differentiation process or persistence of past progenitor signatures at the final time point. The asynchronous nature of embryonic development, for instance in the case of the neural crest^11,12^ and neuro-mesodermal progenitor cells (NMP cells)^13^, represents a partial solution to these challenges by enabling faithful differentiation trajectories to be constructed from lineage-aware gene expression measurements.

While traditionally, lineage tracing has been used to establish the range of cell types that arise from progenitor populations, here we focus on quantitatively mapping the relative frequencies of transcriptional states occurring among the progeny of a cell (clonal fate biases). Even for progenitor cells with identical lineage potency, such frequencies will vary due to the influence of external signals and stochastic differences in the progenitor state. Our aim is to characterize such fate variation and identify factors biasing fate realization in embryonic development. We used a cellular barcoding approach to perform clonal analysis, combining highly multiplexed sparse tracing with unsupervised learning techniques and analysis of expression states, to study the diversity of fate biases occurring in ectodermal and mesodermal clones during mammalian embryonic development. This approach captured gradients of continuous clonal fate variation along anatomical axes as well as clonal fate restrictions over time, enabling computational inference of genes that are likely to influence fate modulation in neural crest and mesodermal progenitor cells. This system of axial clonal patterning was also susceptible to disruption by signaling receptor perturbations, suggesting that the clonal proportionality of somite cell types established by the Hox code during organogenesis can be influenced by modulating Hedgehog signaling.

## Results

### Profiling of clonal behavior in mouse embryos

To reconstruct joint patterns of transcriptional states and cell lineages in post-neurulation murine development, we performed ultrasound-guided nano-injections^14^ into amniotic fluid to transduce mesoderm and ectoderm with heritable, transcribed genetic TREX barcodes^15,16^ (Figure 1d). The injections were carried out once or twice serially between days E7.5 and E8.5 (Figure 1e-F, Extended Data Fig. 1) to capture rapid developmental shifts during embryonic development. Trunk and cranial segments were harvested separately at E13.5 when most definitive cell types manifest, and labeled cells were analyzed by scRNA-seq for linking clonal information to each transcriptome (Figure 1g-h, Extended Data Fig. 2). We optimized the study design to maximize the fraction of cells captured from each individual clone. To accomplish this, we injected embryos with low amounts of virus (in almost all cases 46-206 nL) such that only 10-40k cells would be labeled in each harvested body segment – an amount that could be taken through downstream processing steps without significant additional losses.

Although our microinjection approach predominantly targets ectoderm^14–16^, we found that a substantial portion of traced cells were derivatives of mesoderm. This is because the ultrasound guided injection method showers the exterior of the early neurula with barcoded lentiviral particles, which preferentially labels exposed epiblast cells^14^. As expected, in our experiments, we did not recover traced endodermal derivatives. Following transduction of the embryos, retroviral integration in dividing cells takes approximately eight hours^17^. Two days after the injection we saw the formation of large clones by whole mount analysis of embryos injected with low volumes of viral particles (Extended Data Fig. 3), and broad labeling thereafter (Extended Data Fig. 4), indicating that the tracer system activates swiftly. Thus, this method facilitates rapid, accurate, and broad lineage tracing of early ectodermal cells, as well as mesodermal and NMP cells. Single-cell RNA-seq analysis of eight control embryos (4 cranial, 7 trunk sections), recovered a total of 159,744 clonally barcoded cells. Specifically, we recovered 4411 E7.5-traced clones and 6661 E8.5-traced clones. Of those, 3442 E7.5-traced clones and 4722 E8.5-traced clones contained at least two cells. These informative clones encompassed 49% of all sequenced barcoded cells.

Multicellular clones were distributed across the major embryonic ectodermal and mesodermal derivatives, including keratinocytes of the skin, various mesenchymal cells, and neuroglial cells of the central and peripheral nervous system (Figure 1i-j, Extended Data Fig. 5 and Extended Data Fig. 6). Most clones contained similar cells belonging to only one germ layer derivative (Figure 1k-l; for cranial, see Extended Data Fig. 7). Multipotent clones spanning more than one major lineage were more abundant in the earlier developmental timepoint (1.7% in E8.5 vs. 5.2% in E7.5) even though the same cell types were captured by both injections, indicating that early fate restrictions are already imposed in ectodermal and mesodermal cell lineages prior to neurulation (Figure 1l). The patterns of clonal co-occurrence grouped similar cell types, recovering the expected consensus cell type lineage phylogeny (Figure 1m and Extended Data Fig. 8). Reconstruction of such lineage structure from fixed barcode data, however, is known to be challenging^18^, particularly when tracing cells from various body regions. The resulting phylogeny would represent a population ‘average’ of cell fate decisions, even though clones contributing to any given cell type can vary greatly in their overall lineage distribution (Figure 1b, k).

Lineage phylogeny indicated that at the time of labelling at E7.5, trunk ectodermal cells had committed to either non-neural or neural fates, while the paraxial mesodermal cells remained multipotent across numerous mesenchymal fates (Extended Data Fig. 8). This result suggested that during the midgestational period, mesoderm clones generate a continuous spectrum of clone types, while ectoderm clones diversify into a few discrete patterns of fate combinations. However, to test this hypothesis, a dropout-robust computational approach is needed to analyze the processes underlying the substantial diversity in clonal fate distributions.

### Clone2vec recapitulates spatiotemporal patterns of clonal multipotency in the embryo

To investigate variability among numerous sparsely sampled clonal populations, we developed a computational method, called clone2vec, which adapts word2vec^19^ word embeddings to lineage tracing information^20^. Clone2vec represents individual clones in a low-dimensional space that captures the variation of clonal fates^20^. In such an embedding, neighboring clones exhibit similar cell fate distributions across the continuous expression space (Figure 1p, Extended Data Fig. 9). By establishing this metric space, clone2vec robustly enables exploration of recurrent discrete and continuous clonal patterns to facilitate the subsequent analysis of features associated with them, showing advantages in clonal pattern reconstruction compared to traditional analytical methodologies even with very sparse sampling of cells^20^. These features make clone2vec a useful tool for learning patterns of clonal behavior in the embryo.

Clone2vec analysis of our dataset revealed three major compartments corresponding to derivatives of the neural ectoderm, non-neural ectoderm, and mesoderm (Figure 2a). Clone2vec also facilitated the assignment of clones into clonal clusters within each compartment. The general structure of the clonal embedding reflected the anticipated continuous spectrum of mesoderm clone diversity as well as the clonally separated non-neural and neural ectoderm compartments. The only apparent exceptions to the aforementioned germ layer restrictions were two clonal clusters exhibiting neural-mesenchymal multipotency (Figure 2b, dotted box) representing posterior axial NMPs.

**Figure 2.**
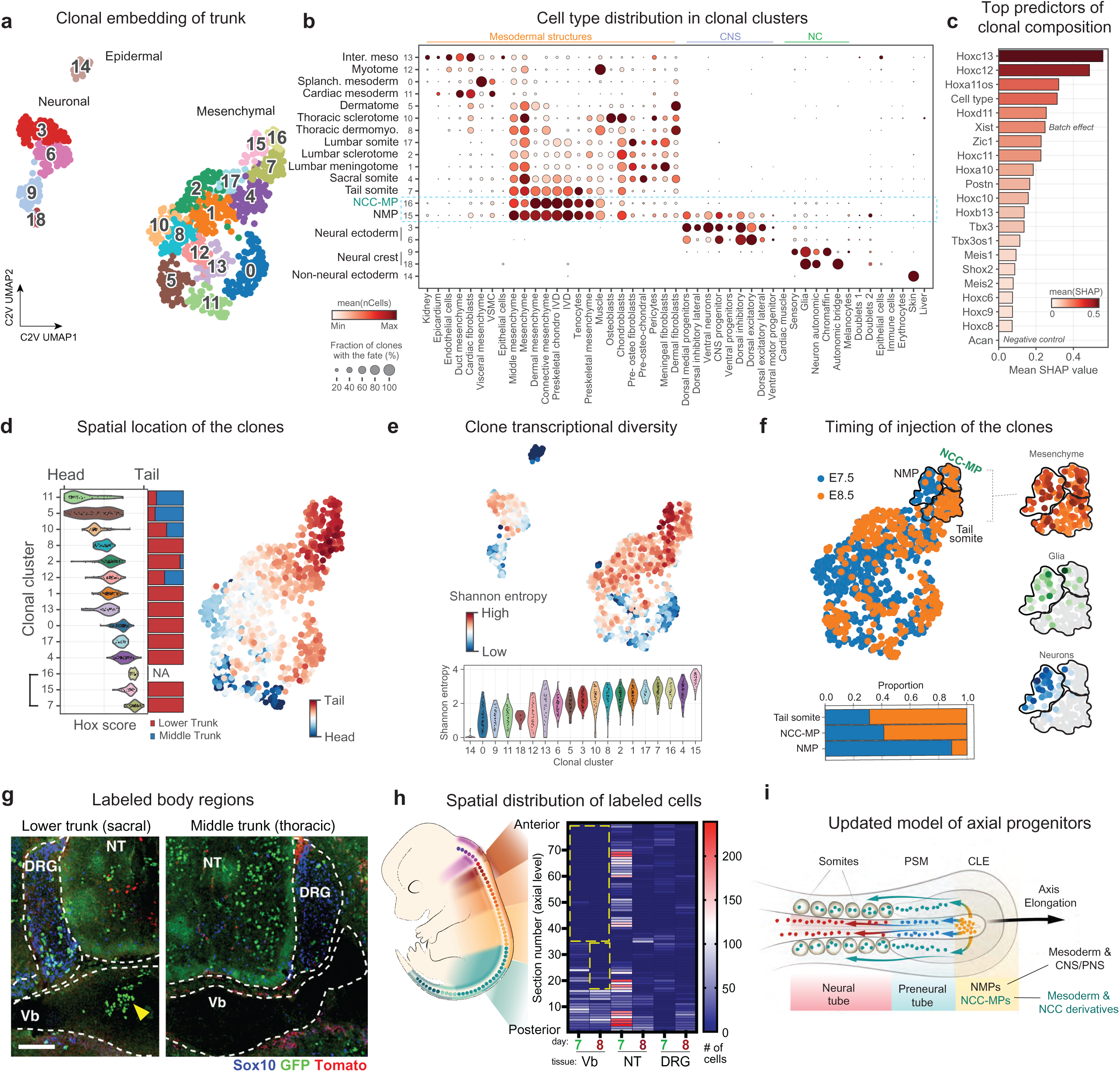
Clone2vec captures spatiotemporal patterns of multipotency in the embryo. **(a-g)** Analysis of the Clone2vec UMAP embedding of E13.5 barcoded trunk cells. **(a)** Embedding color-coded by clonal clusters (clustering resolution 2), labeled according to the fate biases predominantly observed in each of the three main super-clusters (corresponding to neural ectoderm, non-neural ectoderm, and mesodermal derivatives). **(b)** Dot plot showing cell type composition among clonal clusters. Color represents average cells within each fate, per clone, within that cluster; dot size represents the fraction of the clones in the cluster containing that fate. **(c)** Elbow plot showing the top twenty most important genes for predicting cell type composition of clones using the CatBoost model, color coded by the Shapley value (feature importance) for each gene for that model. **(d)** Left, distribution of each clonal cluster Hox gene expression profile, plotted per clone against the known anteroposterior position of embryonic Hox domains (x-axis) and compared to ground-truth manually dissected body regions (horizontal bar plot). Right, feature plot of each mesenchymal clone’s Hox score. **(e)** Top, feature plot showing the number of fates per clone. Bottom, violin plot showing the distribution of Shannon entropy (here used as a metric for transcriptional diversity) per clone within each clonal cluster. **(f)** Top left, mesenchyme embedding color-coded by injection time. Insets highlight the multipotent neuronal-mesenchymal clones in tail mesenchyme (see supplementary figures for validation of NCC-MP clones). Bottom: the composition of each clonal cluster by each injection time. **(g)** Confocal micrographs of a dual injected (GFP, at E7.5; Tomato, at E8.5) E13.5 embryo at sacral (left) and thoracic (right) axial levels, stained for Sox10 (blue), GFP (green) and Tomato (red). Scale bar = 100 μm. Vb; vertebrae. DRG; dorsal root ganglion. NT; neural tube. **(h)** Heat map displaying the labeled GFP^+^ and Tomato^+^ cells within the vertebra, neural tube, and sensory ganglia per section, ordered by axial position. Dotted box highlights the absence of labeled vertebral chondrocytes from anterior sections. **(i)** Schematic of axial extension model featuring canonical as well as NCC-biased NMP clones (termed as NCC-MP, or “neural crest cell and mesodermal progenitor”) in the caudal lateral epiblast. NMP; neuromesodermal progenitor. NCC; neural crest cell. UMAP; uniform manifold approximation projection.

Surveying the mesenchymal fate combinations found among mesodermal derivatives revealed that adjacent clonal clusters often share many cell types, but differ chiefly in their proportionality in which they are found (Figure 2b). Hypothesizing that adjacent mesoderm clonal clusters represent points along a continuum of clonal behavior, we performed principal components analysis (PCA) from discrete versus contiguous clonal clusters (Extended Data Fig. 10a-b) and examined gene expression and cell fate distributions across a constructed trajectory of clones following the top principal components (Extended Data Fig. 10c-d). The clones in contiguous clusters exhibited a continuous gradient of gene expression (Extended Data Fig. 10e) reflecting gradual transitions across mesenchymal cell types (Extended Data Fig. 10f-j). However, because each part of the somite contributes to different structures that form repeatedly along the entire trunk (musculature, skeleton, etc), the clonal clusters could reasonably represent either the micro-anatomy of the somite, or instead reflect general body position along the anterior-posterior axis.

To study the molecular drivers underlying continuous variation in clonal composition, we devised a machine learning approach to identify genes that could serve as indicators of a specific clonal fate distribution (irrespective of cell type). We built a gradient boosting model to predict the position of clones in the embedding based on cell type-stratified gene expression in cells belonging to each clone (see Methods). We then scored gene importance by quantifying each gene’s predictive power using Shapley’s values (measure of feature importance, see Methods). Next, to exclude the most obvious predictors (cell type markers) we added an additional categorical feature (cell type identity) to account for cell type-dependent effects. Applying this approach to the trunk mesodermal clones revealed that Hox transcription factors ranking as the most important features for predicting clonal composition (outperforming cell type info). This result suggests that the observed gradient of fate biases was driven by clones arising from somites at different positions along the body axis (Figure 2c). To test this idea further, we constructed a ‘Hox score’ to locate clones by aggregating their expression levels of Hox genes (Figure 2d, Extended Data Fig. 11a-d). Indeed, clone position along the observed clone2vec fate gradient, and clone Hox score, were linearly correlated (Extended Data Fig. 11c). We exploited this metric to uncover several genes (beyond the Hox code) that are correlated with the body axis (Extended Data Fig. 11e-h). This linear relationship was driven by somite clones, not other mesenchymal cells (Extended Data Fig. 11i-l) nor the cell cycle (Extended Data Fig. 11m). To experimentally validate that the clonal embedding captured the anterior-posterior axis of the embryo, we separately dissected lower and middle trunk prior to sorting, confirming a striking difference in their representation along the mesenchymal domain embedding (Figure 2d). These findings suggest that somite clonal composition and body position are mutually predictive, and that knowledge of a few genes can reconstruct both, even though the cell types in question occur along the entire body axis.

To identify molecular correlates of early biases during progenitor cell differentiation, we examined predictors of clone2vec embedding position of clones, based exclusively on the expression profile of progenitor cells (Extended Data Fig. 12a-b and e, see embeddings of both neuronal and mesodermal lineages). We built a model that predicts cell type composition of the clone based on the expression in progenitor cells (see Methods and Extended Data Fig. 12c-d, f-g). Applying this approach revealed, for example, that during mesenchymal progenitor cell differentiation, the most powerful predictor to be the *Tbx3* gene rather than one of the Hox genes (Extended Data Fig. 12d), and predicted that gene expression of matricellular proteins such as *Col1a1, Col14a1* and *Col12a1* could be useful markers of mesenchymal progenitor fate outcomes. This shows that gene expression indicators of clonal behavior are distinct when restricting analysis to progenitor cells, as compared to when considering all cells (including the differentiated progeny).

### Analysis of neural-mesodermal progenitors reveals unexpected contributions to neural crest

Because the embryo develops from head to tail, caudal cells should exhibit greater developmental potential than their anterior counterparts at a given moment. Clones derived from neuromesodermal progenitors (NMPs, clonal cluster 15 and 16), a cell population that contributes to secondary neurulation and somitogenesis during axial extension, were observed near the tail^13,21^. We noted that, while the posterior-most clones corresponding to NMPs exhibited the greatest multipotency (Figure 2e), levels of clone transcriptional diversity (measured via Shannon entropy) did not always correspond with body position (compare feature plots of mesenchymal clonal embeddings in Figure 2d versus 2e). For instance, tail clusters 15, 16, and 7 had rather similar Hox scores (bracket, Figure 2d), but the most multipotent NMP cluster 16 was enriched with E7.5-labeled clones (Figure 2f), while other tail clusters (15, 7) contained a mix of E7.5- and E8.5-labeled clones, which may reflect fate restrictions in the axial progenitor pool over time.

To visualize the spatiotemporal labeling dynamics of NMP derivatives, we serially sectioned littermate embryos double-injected with EGFP (at E7.5) and tdTomato (at E8.5) barcoded lentivirus and counted clusters of transduced cells from vertebrae and neuronal cells in every section along the anterior-posterior axis (Figure 2g-h). Vertebrae were unlabeled in the mid-trunk but labeled in the posterior trunk, while cells in neural structures such as the neural tube and dorsal root ganglia showed both labels throughout the entire body. Labeling of vertebrae spread longer along the body axis in E7.5 injected embryos as compared to E8.5 injected samples (Figure 2h), consistent with an expected posteriorizing of NMP contributions over time.

We were intrigued that clonal cluster 15 behaved somewhat like NMPs by generating neural crest cells and somite derivatives, but not CNS cells (Figure 2b, f). Consistent with this, we observed some sections with labeling of both crest-derived neurons and vertebrae but few CNS neurons (Figure 2h). Together, this evidence suggests some NMPs can directly generate neural crest during secondary neurulation without contributing to neural tube epithelialization, demonstrating existence of a neural crest - mesodermal bipotent progenitor (here abbreviated as NCC-MP, Figure 2i). To enrich our dataset for this population, we analyzed the tail mesenchyme of four additional embryo samples (Extended Data Fig. 13a-f), again recovering a clonal cluster corresponding to NCC-MPs (Extended Data Fig. 13g-j). Compositional analysis of tail clones revealed that NCC-MPs differ significantly from canonical NMP clones in the number of CNS cells, but both produce similar amounts of neural crest cells; as a result, these clonal clusters are marked by a clear bias towards either CNS or neural crest cells (Extended Data Fig. 13k-m). Given that CNS neurons are more abundant than neural crest cells in our dataset, these clusters are unlikely to be an artifact of dropout (modeling these clones as two clusters describes the underlying data significantly better than one cluster, even when considering higher degrees of freedom in a two-cluster model, likelihood ratio test *p* = 0.00000). Overall, we detected a total of 102 clones containing mesenchyme + NCC fates and zero CNS neurons (at least one clone detected in each of 9/10 embryos), out of 217 NMP clones total (detected in 10/10 embryos). This rare NCC-MP population has been predicted or implied by earlier work ^22,23^ but not yet reported. Further studies of NCC-MP biology might prove relevant to the etiology of some *spina bifida* subtypes.

### Positional gene programs are superimposed on mesenchymal cell maturation trajectories

During the development of the axial skeleton, cells from the somitic mesoderm build the ribs and vertebrae under the control of Hox-dependent patterning, as gain- or loss- of Hox gene function can result in homeotic transformations between ribs and vertebral identity^24–26^, ultimately affecting the cell types produced by a somite. However, how activity of spatial transcriptional programs correlates with differential fate choices among otherwise similar progenitor cell populations remains mysterious. We approached this problem by using clonal embeddings to delineate and compare sets of clones that demonstrate distinct fate outcomes.

Cell fate determination involves coordinated induction of transcriptional programs and, often, suppression of alternate fates, which can in some cases be detected by scRNAseq cell profiling^11^. However, applying this technology to study mesenchymal fate decisions has been limited by the transcriptional heterogeneity associated with mixed populations of overlapping clone types. Indeed, in our transcriptional embedding, trunk mesenchymal populations form a large diffuse cluster (Figure 1j), containing progenitor as well as early differentiated populations (Figure 3a, top left). While standard trajectory inference analyses on such a diverse population would fail due to uncertainty, clone2vec neatly separated the mesenchymal cells into a set of partially overlapping lineages (Figure 3a). Ordering these lineages according to their Hox score confirmed their graded contribution to skeletal cells, dermis, meningeal fibroblasts, and perivascular cells as a function of body position (Figure 3b and boxed insets in Figure 3a, Extended Data Fig. 10-11). We noted that cartilage cells were present in many of these lineages and therefore may be a good model to study position-specific modulation of regulatory programs in the body.

**Figure 3.**
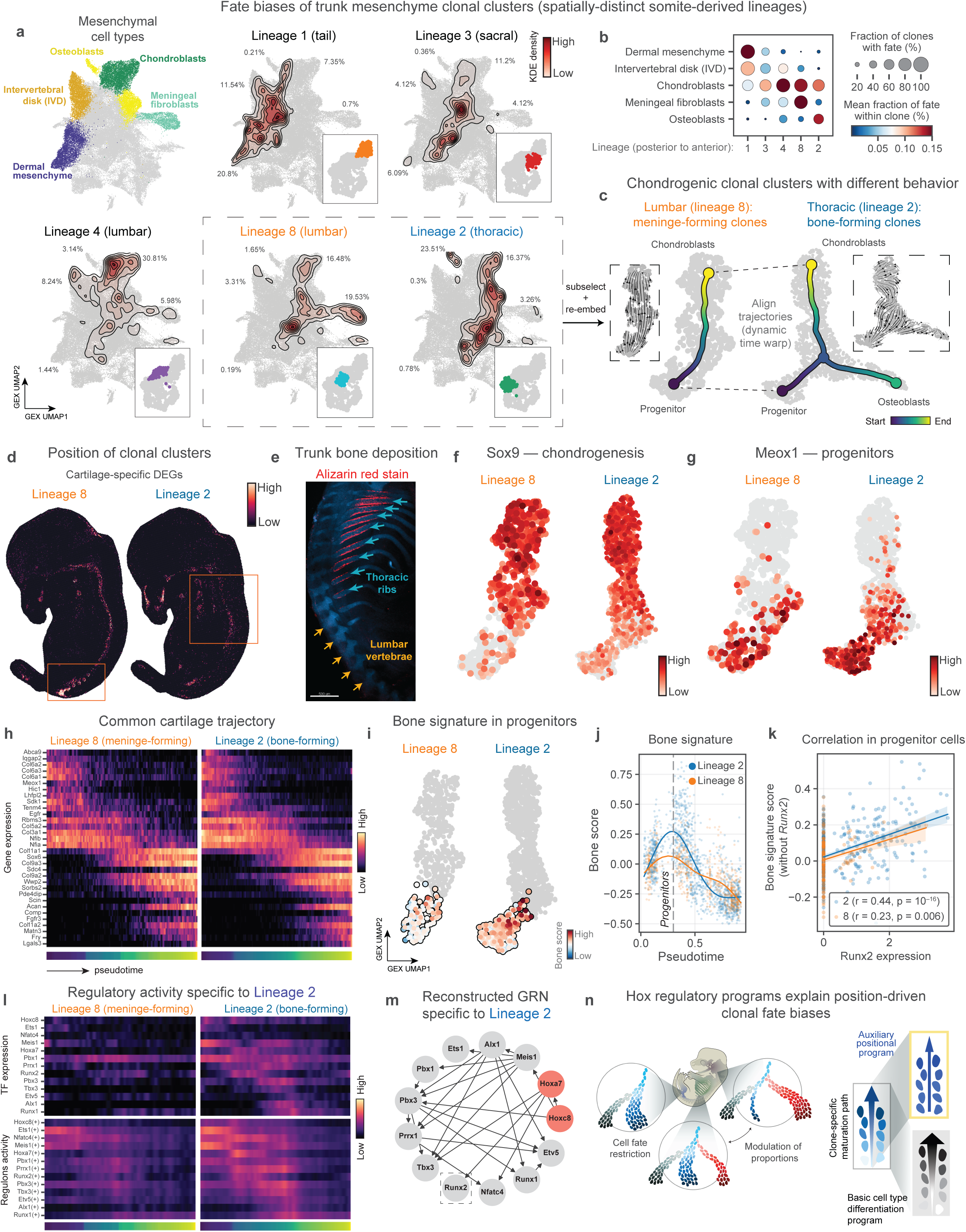
Position-specific programs are superimposed on common paths of cell maturation. **(a)** Kernel density estimate (KDE) plots of cells from each clonal cluster (clustering resolution 1) showing the proportions dedicated to each selected mesenchymal cell type (color-coded in the top left cell embedding). Insets mark the position of the clonal cluster on the mesenchymal portion of the clone2vec embedding (Figure 2a). **(b)** A dot plot summarizing the fate distribution of each lineage shown in (a), ordered by their position along the body axis. **(c)** Trajectory inference and alignment of the chondrocyte-meningeal clone type (lineage 8) and the osteochondral clone type (lineage 2) cells by dynamic time warp (DTW). Dark blue represents the origin of each trajectory (mesenchymal progenitors). Dotted insets show RNA velocity arrows denoting the directionality of cell maturation. **(d)** E13.5 mouse Stereoseq feature plot of gene scores derived from transcriptional profiles of chondrocytes within fibro-chondral (left) versus osteochondral (right) clone types. **(e)** Confocal fluorescence micrograph of a skeletal preparation of the trunk of an E14.5 embryo. Blue indicates cartilage; red indicates mineralized bone tissue. **(f-g)** Feature plots displaying *Sox9* (f) and Meox1 (g) expression per cell within the selected chondrocyte-restricted (left) and bipotent osteochondral (right) clone types. **(h)** Heatmap displaying common gene expression pattern along the chondrogenic trajectories of the restricted (left) and bipotent (right) clone types. **(i)** Expression of bone-related genes in progenitors (pseudotime < 0.3) cells. **(j)** Scatterplot of gene expression of the bone-related gene signature, as a function of pseudotime, in cells belonging to the chondral versus osteochondral clone types. **(k)** Correlation between *Runx2* and bone-related genes’ expression in progenitor cells from these two clone types. **(l)** Heatmap displaying patterns of regulon activity (top) and transcription factor expression (bottom) unique to osteochondrogenic clones. **(m)** Gene regulatory networks predicted within this lineage. **(n)** Model of positional regulation of clone development, in which cell differentiation dynamics (driven by a core set of cell-type-specific genes) is under the influence of position-specific auxiliary programs (such as Hox genes).

To understand the factors driving variation in multipotency profiles across body regions, we focused on comparing two chondrogenic cell lineages – Cluster 8, containing lumbar cartilage/meningeal clones (that did not engage in osteogenic differentiation, here referred to as ‘chondrogenic’), and Cluster 2, containing thoracic ‘osteochondral’ cartilage/bone clones (Figure 3a). To perform the analysis we extracted bone, cartilage and progenitor cells from these lineages, filtered out meningeal cells, and re-embedded them to compare their chondrogenic trajectories (Figure 3c). Mapping lineage-specific cartilage gene expression signatures to publicly available Stereo-seq data confirmed that chondrogenic lineage 8 indeed represent the forming vertebrae in the lower trunk (Figure 3d, Extended Data Fig. 14), while the osteochondral lineage 2 represent anterior structures including early bone collar of ribs, which starts forming from cells in the perichondrium at E13.5 (Figure 3e) ^27,28^. Of note, the presence of meningeal cells in Cluster 8 but not Cluster 2 indicates that the absence of osteoblasts in Cluster 8 could not solely be explained by the asynchronous nature of embryonic development, but rather imply differences in the fate decisions made by thoracic versus lumbar paraxial mesoderm. Indeed, we observed a full spectrum of cell states at E13.5 due to continuous differentiation from mesenchymal progenitors to mature chondrocytes from both clonal clusters (Figure 3f-g), suggesting a *bona fide* developmental trajectory in both cases, although the thoracic osteochondral clones appear to maintain expression of progenitor markers such as *Meox1* for longer (Figure 3g).

We were hopeful that identifying shared and unique features of these anatomically-isolated clonal chondrogenesis trajectories could help distinguish general versus position-dependent aspects of skeletal cell development. We found that both lineages executed a common transcriptional program, upregulating the same suite of standard chondrocyte gene markers (Figure 3h and Extended Data Fig. 15a-d) including *Col11a1*, *Sox6*, *Col9a3*, *Acan*, *Comp*, *Fgfr3*, *Col11a2*, and *Matn3*. In addition, expression of bone-related genes increased at the beginning of both differentiation trajectories, though to a lesser extent in the non-osteogenic lumbar clonal cluster (Figure 3i and Extended Data Fig. 15e-f). Analysis of the bone-related genes with clear early activation in the osteochondral lineage revealed gene expression of the well-known TF *Runx2* in mesenchymal progenitors as the strongest indicator of overall clonal osteogenic potential (Figure 3j), as expected^29^. Despite being downregulated in the non-osteogenic lumbar lineage (Extended Data Fig. 15 g-k), *Runx2* expression significantly correlated with the bone-related signature even in lumbar cells (Figure 3k). Overall, these analyses suggest that dosage of *Runx2* in a mesenchymal progenitor cell can tune its clonal potential for osteogenic versus meningeal fibroblast fates in vivo, implying that fate selection in axial skeletal mesenchyme may occur using multilineage priming, in which some competing transcriptional programs are transiently activated but then decommissioned^11^.

Differential expression analysis contrasting Clusters 2 and 8 implicated numerous TFs related to the Hox code and skeletal development (Extended Data Fig. 16a-h). Using SCENIC^30^, we inferred transcriptional factor activity and predicted gene regulatory networks acting during cartilage development in both Clusters (Figure 3l-m and Extended Data Fig. 16i-j). The analysis suggested an upstream regulatory role of Hox TFs, with *Hoxa10/Hoxd10* coordinating the regulatory network in the lumbar clones, and *Hoxa7/Hoxc8* in thoracic osteochondral clones. Unexpectedly, *Hoxa7* was positioned directly upstream of *Runx2* expression in the reconstructed gene regulatory network (Figure 3m). Additionally, *Hoxc8* (Figure 3m) was situated upstream of mesenchymal TF *Prrx1* (via intermediates *Meis1* and *Pbx3*), which like *Meox1* exhibited sustained expression specifically in the bone-forming lineage 2, suggesting that positional factors might regulate the tempo of early chondrogenesis by maintaining a mesenchymal progenitor status (*Meox1*/*Prrx1* expression). These analyses explain the positional differences in clone skeletogenic potential by predicting a direct regulatory function of Hox genes in mesenchymal progenitor fate decisions, in agreement with observations at later developmental stages^31^.

Overall, combining high-throughput lineage tracing with trajectory inference on cell manifolds^10^ uncovered how the body pattern influences clonal fate biases in the embryo, by decoupling cell type-specific gene modules from spatial gene modules (Figure 3n). Comparing lumbar and thoracic mesoderm revealed 1) the core transcriptional program of axial pro-chondrogenic differentiation, 2) early molecular indicators of osteochondrogenic biopotency, and 3) evidence that spatial programs drive bone collar formation in thoracic cells, potentially explaining why ribs mineralize faster than the posterior vertebral cartilages. Untangling how spatial regulatory mechanisms are influenced by heterochrony, and how this directs specific clonal dynamics such as differentiation tempo and proliferative potential, remains a future challenge.

### Positional genes in facial mesenchyme revealed through clonal analysis

In contrast to the trunk, the anterior cranial region is largely devoid of Hox expression. While previous studies revealed many TFs that are critical for establishing the facial prominences^32,33^, the mechanisms of how spatial programs influence multipotency of facial cells at a molecular level, and the timeline of persistence of spatial programs in differentiated cells, remain insufficiently understood. Our results from the trunk suggested that clonal profiling can proxy for anatomical position of some lineages, which could be useful when only few gene markers are available to locate cells. Thus, we applied our clonal variation approach to this problem, reasoning that a focus on transcriptional variation within a single cell type such as chondrocytes would accentuate position-specific programs by minimizing artefacts related to cell-type composition.

We first tested whether or not craniofacial clonal outcomes would group by anterior-posterior organization. The traced cranial dataset captured a rich spectrum of clonal fate biases (Figure 4a-b), covering populations contributing to central and peripheral nervous system, dermis, meninges, and mesenchymal derivatives of the musculoskeletal system which was segregated into the major lineages corresponding the major facial prominences (Figure 4b-d). Examining the spatial distributions of lineage-specific genes in Stereo-seq data, we found that they consistently mapped to different facial subregions (Figure 4c).

**Figure 4.**
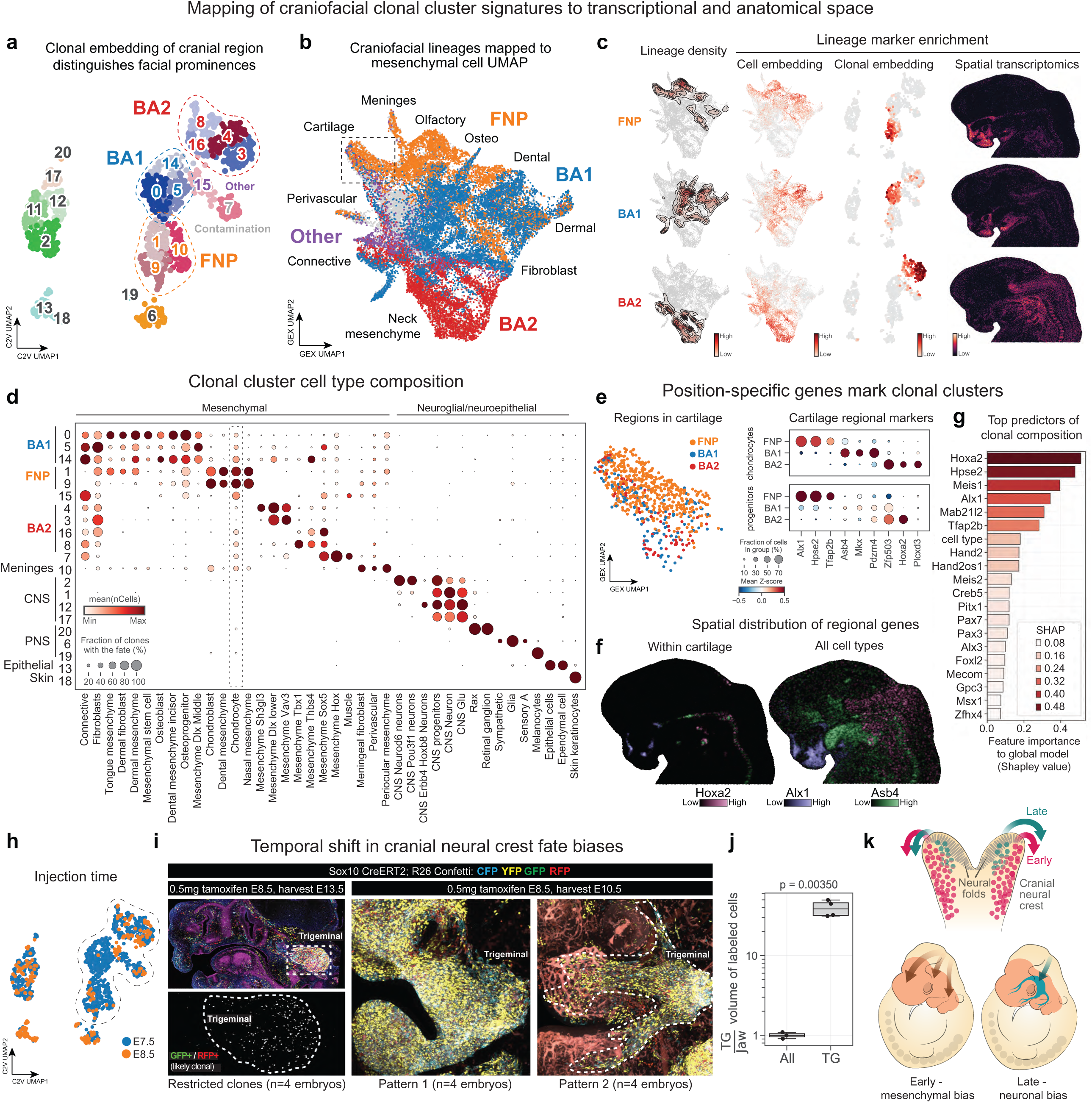
Collective multipotency of the cranial neural crest cell population. **(a)** Clone2vec UMAP embedding of E13.5 barcoded craniofacial cells, color-coded by clonal clusters (clustering resolution 2). The dotted lines indicate the major discovered mesenchymal lineages. **(b)** Cell UMAP embedding of E13.5 barcoded craniofacial mesenchymal cells, color coded by the major lineages. Dotted box indicates the position of cartilage cells (later analyzed in (e). **(c)** Mapping of nose (top), jaws (middle), and neck (bottom)-related clonal lineages by density plots for each lineage (left) and feature plots of aggregated lineage-specific gene expression scores on cell embeddings (second row), clonal embeddings (third row) and E13.5 Stereoseq mouse data (right). **(d)** Dot plot summarizing the fate distribution of each clone type in (a). Dotted line highlights the chondrocyte cluster. **(e)** Top left; isolated chondrocyte cell type colored by lineage. Right; Differential expression analysis of chondrocytes revealed lineage-enriched genes within mesenchymal cells (top dot plot) and chondrocytes (lower dot plot). **(f)** Expression of Alx1, Asb4, Hoxa2 in cartilage (left) and in general (right). **(g)** Elbow plot showing the Shapley value for the top 20 most important features (overall cell type ranked seventh) in a CatBoost model trained to predict cell type composition and position of facial clones. **(h)** Embedding in (a) color-coded by timing of injection. **(i-j)** Confocal micrographs of Sox10-CreERT2; R26R-Confetti embryos. Left; Sagittal section through the head of the embryo traced from E8.5 to E13.5. ERBB3 is labeled in magenta and 2H3 is labeled in dark blue. Cyan, yellow, green, and red colors derive from the confetti fluorescent protein labeling. Bottom left; an inset of trigeminal ganglia (outlined with the dotted line) showing rare double-positive GFP/RFP cells. Right panels; maximum intensity projection of whole mounts of Confetti-traced E10.5-E11.0 embryos traced from E8.5-E9. The proportion of observed tracing in the jaw versus trigeminal ganglia is quantified in (j). **(k)** Model of evolving fate biases in neural crest based on the timing of emigration from the neural tube.

Next, we focused on the persistence of positional factors in differentiated cells. Differential expression analysis of specific chondrogenic signatures within all three major lineages revealed key transcription factors, which were sufficient to reconstitute the expected positional identity (Figure 4e). We then confirmed the cartilage positional signatures (including previously reported markers *Alx1/Asb4/Hoxa2* ^34,35^) were also observed in progenitor cells originating from the same clones (Figure 4e). The anatomical position of these signatures was validated by re-analysis of Stereo-seq data (Figure 4f).

Then, we explored the idea that the detected gene signatures represent position-specific programs modulating progenitor cell fate choices and plasticity. Indeed, fate profiles of cranial clonal clusters revealed that Hox-negative BA1 (branchial arch 1) progenitor-derived clones were highly multi-fated, containing fates from nearly all mesenchymal clusters, whereas Hoxa2^+^ clones from BA2 (branchial arch 2) lacked several mesenchymal cell types (Figure 4d). CatBoost regression-based numeric estimation of gene importance for predicting clone composition and location resulted in a list of top factors including known spatial regulatory factors (Figure 4g). Overall, these results demonstrate how analysis of continuous clonal variation can suggest candidates for positional programs within the developing embryo.

### Cranial neural crest generates neuronal and mesenchymal fates from partly overlapping but mostly distinct progenitor pools

Next, we used clonal embeddings to capture temporal influences on craniofacial clonal multipotency. The cranial neural crest cells (CNCCs) that build our faces emigrate in waves from the dorsal neural fold but encounter different microenvironmental contexts depending on their exact timing of delamination, which makes for a good system to study temporal modulation of cell fate choices in vivo^36,37^. As a population, the cranial neural crest is highly multipotent, giving rise to facial skeletal and other mesenchymal structures as well as to the peripheral nervous system (trigeminal and sympathetic ganglia, including the glial cells), but whether such broad multipotency can be attributed to individual CNCCs remains unclear^37–40^. Additionally, we sought to determine whether clones belonging to distinct waves of CNCCs exhibit distinct fate biases. We hypothesized that in mice, earlier CNCC waves form mesenchyme, while later waves of CNCC instead form neuroglial derivatives, as this has been demonstrated via transplantation experiments in the chicken embryo^40^ and suggested by population-level lineage tracing studies in the mouse^41^.

Examining clonal variation of CNCC lineages, we found that facial mesenchymal derivatives and trigeminal neuroglial cell types were overwhelmingly separated into distinct clonal subpopulations (clusters 6+19 versus clusters 0-1,3-5,7-10,14-16, Fig. 4d) and sparingly few clones shared trigeminal neuronal and mesenchymal fates. Furthermore, these two clonal subpopulations were distinguished by sequential E7.5 + E8.5 double injection, with neuroglial cell types labeled predominantly at later E8.5 injections (Fig. 4h). This result suggests that early and late-migrating CNCC waves indeed eventually become differentially biased. To test the temporal separation between mesenchymal and neuroglial trigeminal cell types in a complementary setting, we performed lineage tracing experiments in *Sox10-CreERT2/R26Confetti* embryos, which enable clonal tracing based on multicolor labelling of stochastically recombined cell populations (Fig 4I). Recombination was triggered by tamoxifen injection at slightly different developmental times (∼12h differences at E8.5 stage). The resulting clonal tracing showed that, although early labeled clones contribute broadly to all facial tissues, later-labeled clones were confined to trigeminal ganglion and outgoing nerves, revealing a systematic difference in the cell type composition depending on the timing of recombination (Fig. 4j).

This result is consistent with a temporal model in which different migratory waves of CNCC undergo differential fate biasing (Fig. 4k), although it is not yet determined whether this biasing is caused by cell-intrinsic features prior to migration, or by differences in their migratory path and post-migratory environments. Overall, this finding demonstrates how clonal variation can help to resolve subtle temporal effects on lineage separation and fate biasing among nearly uniform progenitor cell populations.

### Perturb-TREX: Joint clonal tracing with multiplexed signaling receptor perturbations

The clonal analyses in this study led to predictions about programs modulating progenitor fate, which eventually need to be tested experimentally. Here, mammalian development poses a challenge, as disruption of pathways critical for cell fate determination, even with conditional genetic perturbations, can result in major distortions of the tissue and lead to widespread indirect effects. Mosaic perturbations of individual clones, however, are unlikely to cause such tissue distortions, presenting a valuable opportunity to study the role of genes with strong phenotypes *in vivo*. Because somite axial positional identity was so tightly linked to clonal fate distributions, we wondered how sensitive that pattern was to modulation of the cell signalling environment.

To examine the role of extrinsic signals in early trunk development, we combined TREX tracing with multiplexed CRISPR/Cas9 gene editing of individual clones (Fig. 5 and Extended Data Fig. 17)^42^. Taking advantage of detailed scRNA-seq data on differentiating neural crest and other cell lineages, we extended the barcoded TREX vectors with a library of guide RNAs targeting receptors of signals associated with early cell fate bifurcation steps (Fig. 5a)^11^, as well as a GFP control guide RNA. While designing the library, we noticed from our SCENIC analysis of osteochondral clones (from Fig. 3) that Hedgehog signaling inhibitor *Ptch1* was a predicted transcriptional target of skeletal regulator *Runx2*, which itself is a known target of Hedgehog signaling. We thus included two gRNAs targeting the Hedgehog pathway (*Hhip* and *Ptch1*) as an opportunity to study the role of extrinsic signals during axial patterning of somitic cell subtypes. These multiplexed lentiviral libraries were injected into murine embryos at E7.5 or E8.5 and examined with scRNA-seq and clone2vec at E13.5 (Fig. 5b). While, as expected, none of the analyzed embryos showed major developmental abnormalities, significant fate deviations were observed for mosaic perturbations at the clonal level, resulting in ectopic clonal clusters (Fig. 5c). These clusters seemed to consist mostly of clones crispant for the gene *Ptch1*, which encodes the sonic hedgehog (SHH) receptor ^43^. Indeed, *Ptch1* gRNA^+^ clones were significantly enriched within the ectopic clusters (Fig. 5d), which represented some variation on neural tube clones (clonal cluster/lineage 12), ventral mesenchyme (lineage 9), and somitic mesenchyme clones (lineage 11).

**Figure 5.**
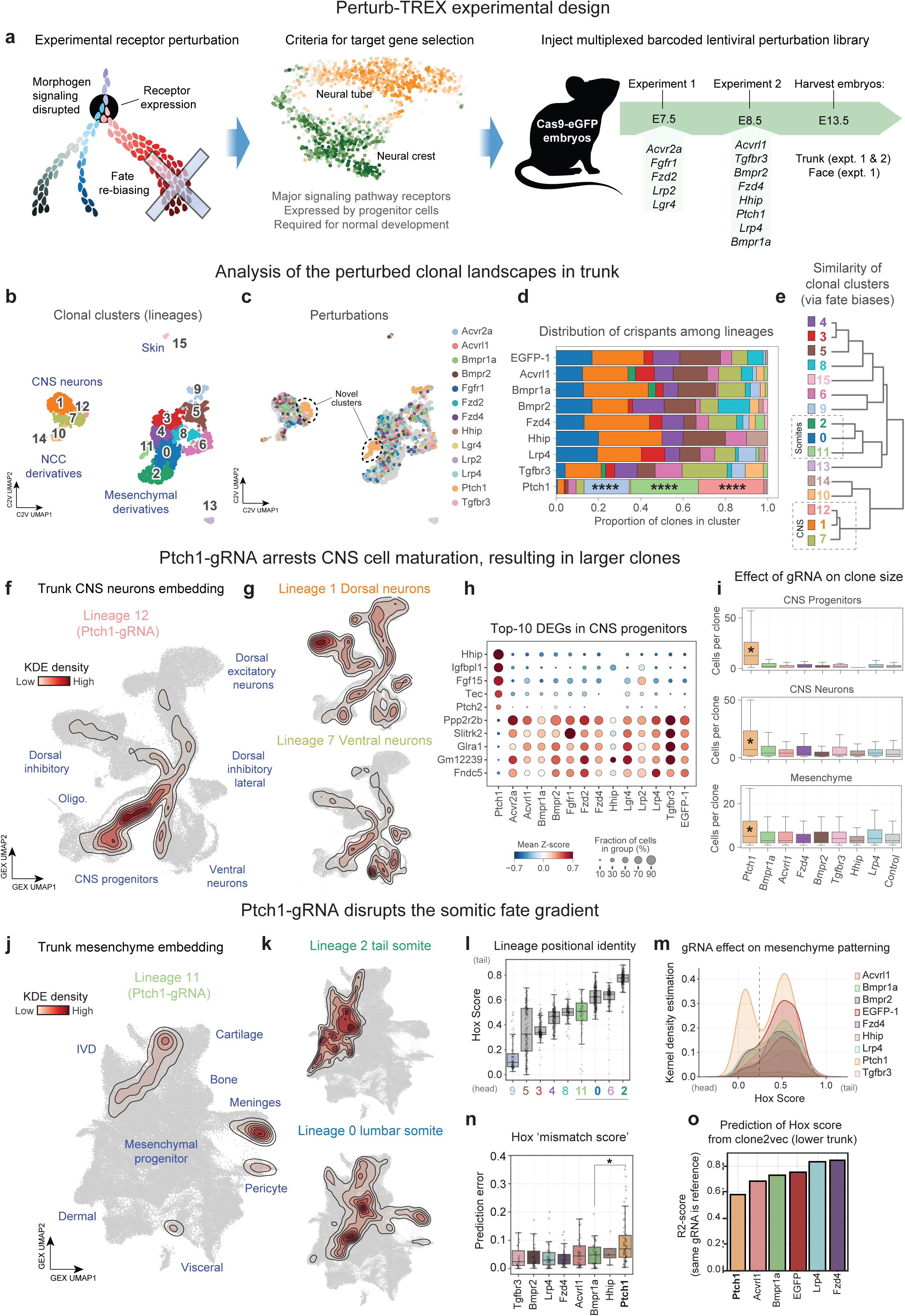
Novel clonal behaviors caused by genetic perturbations. **(a)** Experimental design. **(b)** Clone2vec UMAP embedding of E13.5 control and perturbed barcoded trunk cells, labeled and color-coded by clusters (clustering resolution 1). **(c)** Same embedding as in (b), instead color-coded by perturbation. Dotted lines indicate clonal clusters highly enriched with Ptch1 gRNA+ clones. Note cluster 9 is also highly enriched with Ptch1 gRNA+ clones. **(d)** Bar chart showing the proportion of clone types constituting each gRNA+ population, statistical significance was calculated with scCODA. **(e)** Hierarchical clustering of clone types based on Euclidean distance between centroids in clone2vec space. Dotted boxes indicate the novel clone types. **(f-g)** Density plot of the Ptch1-enriched neuronal clone type (f), and the two clone types most closely related to it (g), based on the clustering in (e). **(h)** Dot plot displaying the top differentially expressed genes by the Ptch1 gRNA+ CNS neural progenitors. **(i)** Box and whisker plot quantifying the distribution of the number of cells per clone within each gRNA+ population (line = median; box = IQR; whiskers = [Q1 - 1.5IQR; Q3 + 1.5IQR]). **(j-k)** Density plot of the novel Ptch1-enriched mesenchymal clone type (j) and of the two clone types most closely related to it (k), based on the clustering in (e). **(l)** Bar and whisker plot showing positional identity of each clone type, quantified by the Hox genes present in each clone. Higher score indicates a position closer to the tail. The label for each clone type is color coded according to the color scheme in (b) and (e). Blue and green bars represent Ptch1-gRNA-enriched mesenchymal clone types (9 and 11, respectively). **(m)** Kernel density estimation plot showing the distribution of Hox scores per clone across different perturbation conditions. Note the bi-modal distribution in Ptch1 crispant clones. **(n)** Error rate in predicting Hox score from nearest-neighbor clones (from clone2vec space). Low values indicate that the clones of a given condition consistently have a similar expression profile for Hox genes relative to neighboring clones, whereas high values indicate clones having different positional identities compared to cells with similar fate biases. **(o)** R2-score (proportion of variance) for each gRNA, reflecting the Hox score predictability of gRNA+ clones, based on nearest neighbor clones (from clone2vec space) bearing the same gRNA. High score indicates high predictability (low variance) in positional identity within the condition, for clones with similar clonal fate biases. Note the low score (high variance) within Ptch1 clones.

Hierarchical clustering to determine lineage similarity was used to find points of comparison for the ectopic clusters (Fig. 5e). As an internal control, we examined the developing CNS, as there are well-known roles for Hedgehog signalling during spinal neuron maturation^43^. *Ptch1*-crispant clones were enriched in progenitor populations (Fig. 5f-g) compared to similar CNS lineages. *Ptch1* crispant clones also upregulated a unique gene expression signature (i.e. *Hhip, Fgf15, Ptch2, Plagl1)* related to cell cycle and Hedgehog signaling (Figure 5h), and significantly increased cell counts per clone (Fig. 5i). As expected, *Ptch1* editing in neuroectodermal clones results in an accumulation of stalled intermediate progenitors prior to their cell cycle exit and neuronal differentiation, indicating that Ptch1-mediated inhibition of Hedgehog signaling is needed for CNS neural cells to mature beyond a transit-amplifying progenitor state. Based on these results, we concluded that *Ptch1*-gRNA clones are indeed undergoing a gain-of-function in the Hedgehog pathway, and next turned to analyse the effects of Ptch1 deletion on the multipotent somitic mesenchymal fate gradient.

Somite clones lacking *Ptch1* exhibited a complex phenotype, and the ectopic clone type resulting from elimination of *Ptch1* from mesoderm-derived mesenchyme was not found in any other analyzed conditions (Fig. 5j-k). As predicted, based on the described feedback between *Ptch1* and the *Runx2* gene, the *Ptch1-*gRNA clonal cluster was almost entirely depleted of bone cells. Instead, this distinct clone type was enriched with vertebral chondrocytes and meningeal cells (much like the lumbar path 8 in Figure 3) but was also strikingly and specifically depleted of undifferentiated mesenchyme (Fig. 5j, compared to control clusters in Fig. 5k). Since apoptosis-related gene expression did not differ across gRNAs (indeed, *Ptch1* clones are larger, Fig. 5i), an enticing explanation could be that unlike in the CNS, gain-of-function of Hedgehog signaling in mesoderm might increase differentiation tempo, depleting the mesenchymal progenitor pool.

Given that somites occur along the body axis, we were surprised that *Ptch1* editing resulted in a single perturbed somite lineage and questioned whether this was driven by mis-patterning or differential local sensitivity. We compared the Hox score of mesenchymal-biased lineages including *Ptch1* gRNA-enriched clusters 9 and 11 (Fig. 5l), finding that they drastically differed in positional identity, suggesting an effect of *Ptch1* editing on mesodermal axial patterning. Indeed, analyzing Hox score per gRNA condition revealed a bimodal distribution of Ptch1 clones but no other condition including EGFP controls. Cluster 9, with anterior identity (and found primarily in middle trunk samples), was primarily biased towards duct mesenchymal cell types forming from the ventral part of the somite (Extended Data Fig. 17e, see location of gene expression score in Extended Data Fig. 14h) suggesting that *Ptch1* perturbation caused ventralization of anterior but not posterior mesoderm. On the other hand, the somite Cluster 11 (posterior identity, found primarily in lower trunk sample) contained clones from a wide range of Hox scores, some significantly anterior compared to nearest neighbor lineages from lumbar and tail somites (Figure 5l), suggesting that the axial pattern of clonal fate biases was disrupted. Accordingly, Hox scores of *Ptch1* gRNA^+^ clones were significantly less predictable than clones bearing other gRNAs, when using any nearest neighbor clone as reference (Figure 5n), and even when using the same gRNA as reference (Figure 5o).

Taken together, a highly predictable embryo-wide gradient of clonal fate biases is observed under natural conditions. This clonal organization can be overridden by the activation of cell signaling pathways, via either a potential interaction with the patterning mechanism, or strong influences of Hedgehog gain-of-function on cell state irrespective of positional identity. Future studies focusing on the lineage-specific interactions of *Ptch1* with cell differentiation programs should be performed using detailed comparison of the regulatory programs acting differentially across lineages coupled with combinatorial perturbations along a time series.

### Principles of cell fate biasing in neural crest revealed via clonal analysis

We next focused on the trunk neural crest, which has been debated as being either a mixed population of restricted progenitors, or a population of highly multipotent stem cells^12,36,44–46^. Clonal variation analysis of wildtype embryos revealed that all detected neural crest sub-lineages were highly fate biased. This was observed for sympathetic, melanocyte, somatosensory, enteric, and glial progenies (Figure 6a-c). Importantly, as expected, none of the neural crest lineages were clonally linked with mesenchyme (Figure 6c), and very few crest-mesenchymal bipotent clones were observed (aside from clones arising from the aforementioned NCC-MPs). Thus, although vagal neural crest gives rise to both cardiac mesenchyme and enteric neurons, this multipotency is more accurately attributed to the collective population, and not to individual neural crest cells. Instead, our results suggest that prior to clonal expansion, labeled neural crest progenitors acquire strong fate biases, corresponding to the biases described for different migratory waves of neural crest^36,47^. Such progenitor differences may be too subtle to be observed from transcriptional analysis alone^48^. For instance, we find that the progenitor cells in clones which are destined to give rise to either the *Phox2b^+^*autonomic nervous system, or the enteric nervous system within the developing gut, both map to the same overall transcriptional state (Figure 6d-e). The knowledge of clonal identity enables a more powerful way of identifying biasing programs by directly contrasting the progenitors with distinct fates. In this way we find that these early progenitors show robust upregulation of a set of biasing genes, including *Hand2* - a transcription factor known to be involved in neural crest enteric differentiation^49^. Since such genes reflect early progenitor biasing mechanisms that appear prior to overt differentiation, their modulation may be helpful for cell type generation protocols and applications intended to cure autonomic and enteric dysfunctions, including Hirschsprung’s disease^50^.

**Figure 6.**
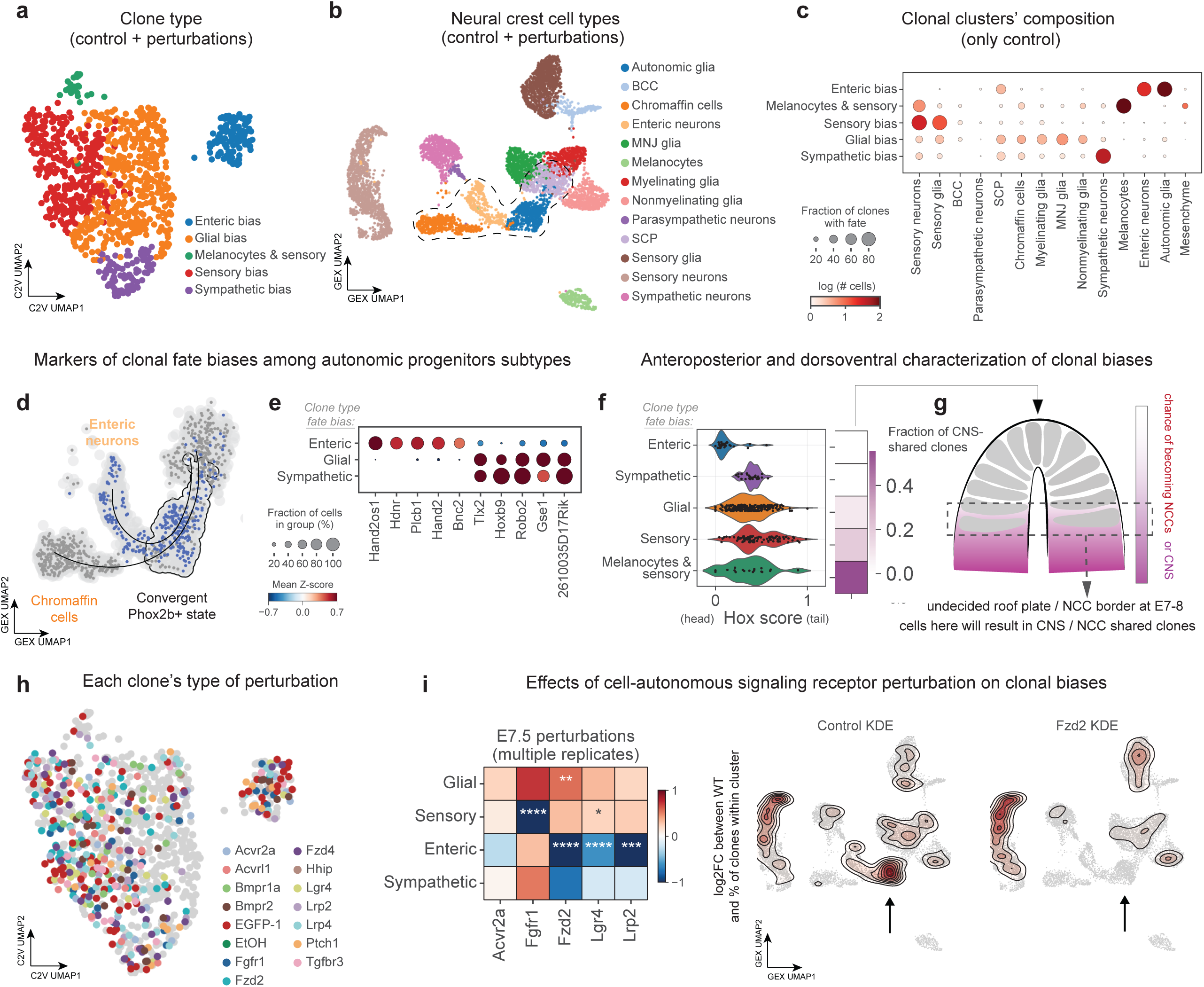
Multipotency landscape of trunk neural crest in normal and perturbed conditions. **(a)** Clone2vec UMAP embedding of E13.5 control and perturbed barcoded trunk neural crest-related cells (with more than 2 cells), color-coded by clone types. **(b)** UMAP cell embedding of the same group of cells shown in (a), color coded by the cell type. Dotted line shows enteric and chromaffin trajectories. **(c)** Dot plot showing cell type distribution of each clone type. Dot size corresponds to the proportion of clones including a given fate; Dot color corresponds to the average number of cells with that fate per clone. **(d)** Cell UMAP of the diverging enteric and chromaffin trajectories. Highlighted region indicates the common Phox2b+ glial progenitor state. **(e)** Dot plot showing gene expression profiles specific to enteric versus glial and sympathetic lineages within that shared state. **(f)** Predicted anteroposterior positions based on Hox gene expression across all clones within each clone type (left) compared to the proportion of clones shared with CNS cell types shown as a vertical heatmap (right). **(g)** Model of neural tube development in which dorsal NCC progenitors emigrate prior to ventral cells. This differential timing spent in the tube could affect the number of cell divisions, resulting in different proportions of CNS cells in the clone). This model results from the plotted data in panel f showing the fraction of neural crest clones shared with CNS. **(h)** Embedding as in (a) color-coded by perturbation. **(i)** Left; heat map showing fate distributions shifted by each perturbation. (Beta-regression LRT-based p-values with FDR multiple testing correction was used for significance estimation). Right; Density plots of control (left) and gRNA+ (right) clones on the NCC cell UMAP embedding. Arrows indicate the specific loss of enteric fates.

Because of the findings in the cranial neural crest and somites, we were also curious about the extent to which anterior-posterior (or dorsal-ventral) position, and clonal fate biases, could be used to inform one another in this lineage. We examined positional distribution of the neural crest multipotency along the anterior-posterior and dorsoventral axes. Based on the *Hox* gene expression we find the expected strong anterior bias of the enteric-biased clones^51^, as well as paucity of sympathetic clones in the posterior sacral and post-sacral parts of the body (Figure 6f). The remaining clonal clusters showed broad distributions along the body axis. As a proxy for the neural crest cell progenitor position along the dorsoventral axis within the neural tube, we examined the co-occurrence of neural crest and neuronal CNS fates within the same clones. Since neural crest is formed by successive waves of migration of cells from the most dorsal part of neural tube, increasing contribution of clones to CNS neurons would suggest a more ventral position - closer to the border between the prospective crest domain and the future roof plate of the neural tube^44^. We find that the clones biased towards somatosensory neurons and melanocytes shared a much higher proportion of progeny with CNS neurons, consistent with a model in which the originally transduced progenitor cells were positioned more ventrally (Figure 6g). Given that sensory and pigment fates are thought to arise from later-migrating waves of neural crest, these results are reminiscent of previous spatiotemporal fate mapping of the dorsal neural tube in chick embryos^36^.

Finally, to better understand the signals biasing neural crest clones, we examined the compositional changes associated with CRISPR/Cas9 perturbations of different receptor genes. Our previous work^11^ suggested that neural crest cells undergo transcriptional bifurcation that determines autonomic or sensory fates. We hypothesized this bifurcation point would be sensitive to the cell signaling environment, leading to reciprocal effects on autonomic versus sensory clonal biases. Receptor perturbation caused significant and opposing shifts in clonal composition of autonomic vs somatosensory progenies in the neural crest lineage, suggesting that the corresponding receptors and signaling pathways are likely involved in defining proportions of fate outcomes (Figure 6h-i). For instance, *Fzd2, Lrp2*, *Lgr4-*crispant clones had reductions in autonomic and enteric biases, accompanied by increases in sensory and glial fates. The perturbation of *Fgfr1* showed exactly the opposite effects on clonal biases and composition. These results expand our understanding of the role of signaling receptors in neural crest specification by providing evidence supporting the existence of a sensory-autonomic clonal lineage bifurcation^11,52^ and its modulation by putative FGF and WNT signaling molecules.

## Discussion

Context-dependent regulation of cell fates is a fundamental aspect of embryogenesis, essential for the proper formation of tissues. Tracing of clonal structures can reveal the spectrum of such fate modulation and provide important clues about the mechanisms involved. Embryogenesis, however, presents a particular challenge for modern lineage tracing approaches, since the large number of generated cell types and the diverse position-specific nature of modulation demands both increased resolution and scale of tracing. Furthermore, even though different developing skeletal structures are often made from the same few cell types^53,54^, these structures acquire distinct shapes and functions due to the fine regulation of cell fate proportions of cell progenitor populations^55,56^, which develop from spatially distinct subsets of the same cell lineage^57^. Insights into the continual variation of clonal structures can help explain how these morphogenesis-related differences gradually arise in a coordinated and context-aware manner.

In the past, low-throughput clonal lineage tracing analysis has been used extensively to successfully address hypotheses about cell multipotency and differentiation^45^. Although these studies have established the classic paradigms about lineage splits in embryogenesis, they were not sufficiently powered to investigate the continual variation of clonal fates. Unless paired with live imaging, fluorescence-based techniques generally suffer from ambiguities when defining a clone^37^. A resolution that would enable biological discoveries has been difficult to achieve on an embryo scale, and to date, most high-throughput tracing studies have focused on specific cell lineages such as hematopoiesis or forebrain development, instead of the numerous connective tissue derivatives of mesenchymal cells ^15,58^.

Here, we present an effective practical approach for studying cell fate specification: 1) quantitative *in vivo* fate mapping of thousands of individually barcoded clones across developmental timepoints, 2) computational analysis of gene programs associated with clonal fate biases, and 3) mosaic multiplexed perturbations. Applying this, we generated a large-scale clonal atlas of ectoderm and mesoderm, which facilitated studying gene programs and spatiotemporal factors that influence continuous and discrete clonal variation in the embryo.

However, even the most state-of-the-art high-throughput lineage tracing methods still sample only a small fraction of cells in the embryo. The sparseness of the resulting clonal data represents a major challenge for subsequent analysis. This has prompted us to develop a machine learning approach – clone2vec^20^. Starting with a continuous representation of expression state, clone2vec embeds sparse clonal data into a low dimensional space where clones with similar or compatible fate profiles are grouped together^20^. In contrast to the methods aimed at analysis of factors associated with specific cell fate choices (CoSpar^59^ or CellRank^60^), clone2vec aims to provide an effective overview of clonal variation, enabling exploration of recurrent patterns and subsequent analysis of associated molecular features. The continuous representation of the expression space enables a shift away from a cell type-centric perspective of clones’ classification, allowing for a nuanced description of state patterns in clone distribution. This approach, as we show, can recover complex clonal fate patterns in a manner highly robust to dropouts^20^, and permits generation of new hypotheses. In addition to separating distinct progenitor lineages (*e.g.* mesenchymal and neural), clone2vec embeddings capture continuous clonal variation within these lineages^20^.

In contrast to tree representations^6–8^, clonal embeddings enable analysis of gradual variation of clonal fates and inference of regulatory processes modulating the proportions of cell types. Clones within different spatiotemporal contexts display distinct but overlapping transcriptional signatures^61^. For example, the trunk and cranial mesenchymal cells both form large unstructured clusters of cells in the transcriptional embedding, posing a challenge for the traditional methods of trajectory reconstruction and other analysis. Lineage tracing and clone2vec analysis, however, effectively partitions such a population into a series of clonally connected sub-clusters, revealing the primary drivers of variation (Figure 3a). Such nuanced clonal patterns cannot be easily discerned using traditional approaches such as hierarchical clustering (Extended Data Fig. 11n).

This approach led to the finding that the anterior-posterior axis is strongly predictive of clonal fate biases, on an apparently continuous basis in the paraxial mesoderm, while in cranial neural crest clones this effect appeared discretised by pharyngeal arches. In the somite lineage, our early-labeled mesodermal progenitors likely contributed to multiple somite subdomains; thus, labeling at later stages would instead be expected to yield clonal clusters that more directly mirror somite sub-regional anatomy. Conversely, an A-P axial gradient in fate biases was not evident from trunk neural crest, aside from well-known anatomical stream structures (e.g., enteric, adrenal) and contributions from NMPs. While earlier-stage labeling could reveal analogous positional effects, such experiments would depend on further advances in the labeling method. Instead, trunk crest lineages seemed to differ by their clonal proximity with CNS, which, given the rapid rates of cell division in the embryo, we interpret as being linked to the time spent in the neural tube. This is expected to be driven by differences in the initial dorsal-ventral position in the neural tube^36^, which in turn could be plausibly mapped to specific mediolateral positions at the neural plate border. Future studies in which lentiviral barcoding is paired with promoter-specific recombination could test such a model by fate mapping individual ectodermal subdomains.

The analysis of transcriptional variation between clonal clusters can reveal more subtle spatial programs that regulate clonal structures. For instance, we observed position-dependent variation in skeletogenic mesenchyme, and identified transcriptional programs downstream of the Hox code that do not appear to be essential for chondrogenic differentiation, but rather modulate secondary features such as a probability of transiting into osteogenic fate or maturation dynamics of progenitor cells. Such an approach should be useful for analyzing position-dependent modulation of progenitor potency in other contexts such as during tissue regeneration^62^. Further, distilling cell-type versus spatial differentiation programs for more cell types might aid optimization of efficient and parsimonious *ex vivo* differentiation methods needed for regenerative medicine.

In addition to spatial modulation of clonal dynamics, we also observed heterochrony of clonal fate biases by introducing static barcodes at precise developmental stages. This can be contrasted with other methods employing dynamic barcodes^8,9^, in which the timing of lineage bifurcations can be difficult to pinpoint, complicating efforts to link lineage decisions to the corresponding signals and developmental stages. Our comparison of cranial neural crest cells traced from two successive time points showed temporal segregation of facial skeletogenic mesenchyme and trigeminal ganglion from the cranial neural crest, which was noted in chick ^40^ and suggested previously based on population-level lineage tracing experiments in mouse embryos ^41^. Our finding suggests that temporal windows shaping cell potency are essential for building an integrated head, as neural crest gives rise to structures as distinct as peripheral nerves and facial bones^63^. Furthermore, the wide range of mesenchymal fates in facial clones detected in this study, compared to previous studies of traced neural crest cells, indicates a steep decline in developmental contributions within a short time span^37^. Finally, though we cannot conclude that CNCC are fate restricted per se, the absence of neural-mesenchymal bipotent clones in both studies is consistent with multiple progenitor pools, given that the different CNCC waves seem to emerge from separate regions of ectoderm^41^. Overall, the diversity and range of facial shapes produced during vertebrate craniofacial morphogenesis likely benefits from layers of dynamic temporal control that complement positional cues^64,65^.

The coupling of clonal lineage tracing with mosaic perturbations facilitates studying cell fate regulation across spatiotemporal niches. Perturbing the signalling environment resulted in reciprocal fate biasing of neural crest clones by FGF and WNT pathways, suggesting a competitive interaction between sensory and autonomic fates that was previously hinted at by transcriptomic profiling^11^. Furthermore, disrupting signaling axes such as Hedgehog (via *Ptch1*), results in formation of ectopic clonal clusters in which cell differentiation appears accelerated or retarded, suggesting a new opportunity to study tempos controlling development. Since mosaic perturbations impact only a very small fraction of cells in the tissue, this approach can be used to evaluate the impact of signaling axes that would otherwise severely distort the tissue or lead to unviable states. We anticipate a future with mosaic multiplexed perturbation studies aimed at quantitatively investigating cell fate regulation in native tissue environments under both normal and diseased conditions.

In addition to insight into signaling landscapes and positional cues, high-throughput analysis of clonal biasing in the mammalian embryo uncovered previously unrecognized lineage relationships or fate restrictions. For instance, that NMPs – multipotent mesoderm-neuro competent cells^13,21^ - coexist in the tail region with mesoderm-restricted progenitors. Our results yielded transcriptional signatures that separate the E13.5 derivatives of NMPs from that of their fate-restricted neighbors (*i.e. Ran*, *Hoxb13*) that could be helpful in determining whether a cellular memory of NMP multipotency exists. This study also identified a population of murine NMPs that contribute neural crest cells but not spinal cord neurons (so-called NCC-MPs)^22^. While classical models assume that neural crest arises from the neuroepithelium, our findings suggest that a subset of NMPs gives rise to neural crest independently of the CNS, which might represent a novel mechanism of neural crest induction. This implies that axial progenitor pools are more diversified than previously thought, and their regulation will influence the balance between CNS, neural crest, and mesodermal cells during secondary neurulation and body extension. While our results provide a gene signature to distinguish them (*i.e. Tspan18, Pde8a*), further studies will be needed to determine if neural crest cells derived from NCC-MPs bear functional differences from other sacral crest cells, and how differences in NCC-MP behavior affect spinal cord development and function.

On a larger developmental scale, our analysis reveals a picture of global fate restrictions at the stages of late gastrula and early neurula, which altogether opposes the idea of early plasticity of different germ layers as was suggested from classic experiments in vitro^66^. According to our data, most clones do not span the competence of more than a single germ layer, except for the NMPs and a handful of exceptional clones from the cranial neural crest. This does not rule out that cells from these stages might exhibit different multipotency profiles when subjected to different environments, as in the case of transplantation experiments that can reveal greater cell plasticity^1,40^.

Overall, we hope the presented approach will be helpful for further studies aimed at systematically dissecting the spatial and temporal correlates of clonal variation in different cell systems. The current clonal atlas, including its interactive interface (browsable online at https://clones2cells.streamlit.app) should provide a useful resource for a wide range of questions related to the formation of boundaries between developing embryonic structures^67^. Similarly, clone2vec analysis should aid in the identification of fine-grained positional and temporal codes. While the Hox code and other crude positional systems provide a general framework for the development of body axes and programming the large body domains^25^, the fine-grained signals that guide specific cell behaviors within very local tissues are much harder to study due to the complexity and redundancy of embryonic signaling systems^68^. Clone2vec may assist investigations of how fine-grained positional information informs cell diversity within individual organs like the heart or lungs. This might shed light on other unresolved questions, for instance, the enigma of the anterior-posterior allocation of brain nuclei^69^, or finer patterning of mesodermal populations building individual muscles^70^. Further, joining clone2vec clonal analysis with mosaic perturbations enables multiplexing simulated conditions that mimic human genetic abnormalities, including neurodevelopmental disorders and pediatric cancers (i.e. neuroblastoma). This might help reveal clonal compensatory behavior in modelled congenital diseases, as well as in regeneration, potentially revealing clonal mechanisms of embryonic resilience^71^.

Despite the potential, we realize the limitations of the study. Comprehensive and targeted validation of the cell-fate regulatory interactions predicted by the current study will be an important next step. The presented multiplexed tracing and perturbation approach can be widely applied for focused analysis of specific tissues and compartments, however, the potential to refine these techniques is substantial. Current limitations include the constrained developmental time frames and specific cell types that can be effectively targeted using existing barcoding techniques. Inference of differentiation paths or the state of progenitor cells based on one measured endpoint poses well-known fundamental challenges, particularly when the underlying process is asynchronous and one cannot observe, for example, both progenitor and differentiated cells from the same clone in the snapshot provided by scRNA-seq data. While we show that successive barcoding rounds can help, incorporating dynamic barcoding strategies such as CRISPR scarring^6–9,72^ is likely to improve path reconstruction, and systematic variation of experimental endpoints might assist progenitor state inference.

## Acknowledgements

General acknowledgements

We thank Olga Kharchenko for the lovely illustrations. We thank Ruslan Soldatov for the insightful discussions. We thank Daniel Wies, Adrian Martinez Martin, and Polina Kameneva for their technical assistance. We acknowledge the National Genomics Infrastructure, Biomedicum Flow Core, Biomedicum Imaging Core, Advanced Light Microscopy Core at SciLifeLab, and Biomedicum Comparative Medicine for access to their facilities and services during the project.

## Author contributions

Conceptualization: IA, PVK, AGE, SI

Investigation: IA, PVK, AGE, SI, MR, JF, EA, BS, JH, JF

Validation: AGE, SI, AM, BS, EA, JH

Computational investigation and analysis: SI, AGE, MR, PVK

Writing – original draft: IA, PVK, AGE, SI, EA, AC, BS, JF, MR, AM

Funding and resources acquisition: IA, EA, AGE, JF, PVK

Supervision and project administration: IA, EA, AGE, JF, PVK

## Materials and Methods

### CloneID and gRNA-CloneID library construction

Barcoded lentiviral libraries were generated based on Ratz et al^15^. Briefly, a lentiviral backbone was constructed containing human elongation factor 1 alpha (EF1a) promoter driving expression of either tandem dimer tomato (tdTomato) or green fluorescent protein fused to histone 2b for nuclear localization (H2B-EGFP). For perturbation libraries, a gRNA scaffold using capture sequence “cs1” within the hairpin loop was inserted to LV-U6-tdTomato-NLS constructs using Gibson assembly. Then, the resulting vector was digested using BsmBI and separately annealed gRNA oligonucleotide sequences (which were chosen from the validated sequences in the Broad Institute’s Brie CRISPR knockout library) were cloned into the gRNA scaffold by ligation. The resulting vectors were amplified by transformation by heat shock into ONE-SHOT STBL3 competent cells, isolated using a miniprep, and sequence-verified by Sanger sequencing. Next, control backbones (or equimolar mixtures of gRNA-containing backbones) were used for insertion of an amplified random 30N oligonucleotide library via Gibson assembly. The library was further amplified via electroporation of Endura electrocompetent cells. The barcoded plasmids were then subjected to third-generation lentiviral packaging with a VSV-G envelope and concentrated by ultracentrifugation by KI Virus Tech facility or by GEGTech (Paris, France). The resulting ultra-concentrated viruses had high functional titers greater than 10^9^ TU/ml.

### Animal experimentation

All experiments were approved by the Swedish Animal Welfare Body Jordbruksverket De regionala Djurförsöksetiska nämnderna and Stockholms Djurförsöksetiska nämnd, permit number N15907-2019/18314-2021. Animals from non-genetically modified mouse lines CD-1 and C57BL/6 were purchased from Charles River Laboratories and Janvier. Animals were kept in an SPF animal facility with standardized conditions (24 °C, 12:12-h light–dark cycle, normal humidity, food and water ad libitum). CD1 or Cas9-GFP mice were placed together into time-controlled matings, and the next day the female was plug checked – if found positive, this was counted as embryonic day (E) 0.5.

### Ultrasound guided injections (single injections)

Injections for lentiviral gene delivery were performed according to previously published protocols^14^. A glass capillary was pulled, and the tip was grinded to create an ultra-sharp needle. At embryonic day E7.5-E8.5, the females were anesthetized with isoflurane (4% induction, maintained at 1.5-2.5%) and placed on a heated plate for surgery. Depth of anesthesia was tested through pinching the paw. Eye gel was applied to prevent dehydration. Pain relief e.g. Buprenorphine 0.05-0.1 mg/kg was administered via subcutaneous injection shortly before the first incision. All instruments were autoclaved and disinfected with chlorhexidine alcohol before surgery. The skin was disinfected with chlorhexidine alcohol, and the hair on the abdomen was removed with a shaver or hair removal agent such as Veet. A small incision was placed in the abdomen and the uterus was exposed and pulled through an elastic bottom of a modified petri dish with PBS. The uterus was kept moist with warm NaCl solution (37C). Embryos were visualized using a Vevo2100 ultrasound imaging system. Using the glass capillary, lentivirus (from 23-206 nL) was injected through the uterine wall into the amniotic cavity. After the embryos were injected, the uterus was brought back into the abdomen and the incisions in the muscle wall were sewn together with absorbent suture thread, and the skin was closed with suture thread or staples. The mice were taken off isoflurane and allowed to wake up in their own cage on a heating pad under supervision. Postoperative pain relief was given in the form of a subcutaneous injection of Buprenorphine (0.05-0.1 mg/kg). For embryo retrieval, the mother was euthanized via CO2 overdose and cervical dislocation, and the embryos were collected at E13.5 and euthanized via decapitation.

### Ultrasound-guided double injections

To determine temporal aspects of general fate restrictions in vivo, two sequential uterine injections in pregnant CD1 mice were performed. A first injection of 206 nL (corresponding to ∼313000 predicted unique CloneIDs) pMR532-LV-EGFP-bc lentivirus was done on mouse embryos from embryonic day 7 (TS11). A second injection of 69nL (corresponding to ∼72,000 predicted unique CloneIDs) of pMR671-LV-TOM-bc was performed at an interval of 12-24 hours after the first. There was no observable scar tissue formation on the day of the second injection. The ultrasound-guided injection itself took place as described in the above section. The embryos survived without visible developmental delays after the single or double injections. Embryos were euthanized by decapitation at E13.5, and cells from the most highly labeled embryos were harvested for downstream analysis as described above, while the remaining littermates were fixed overnight in paraformaldehyde at 4C for histological analysis of cell labeling distribution. During and after the second injection, the mice were given the following pain relief protocol: 5 mg/kg of the long-acting anesthetic Meloxicam, 2 mg/kg of the local anesthetic Bupivacaine, and 0.1 mg Buprenorphine. We tested this protocol for pain relief after double injections 24 hours apart. Together with the veterinarian, we used 3 females to evaluate the effectiveness and impact of the protocol. We monitored the females for pain and recovery from the procedure, assessed the healing of the surgical wound, and checked the status of the fetuses toward the end of the procedure. All females recovered quickly after the first and second injection (within 10 min of placing the cage on a warm pad, the females were walking around the cage and eating normally). The animals showed minimal signs of discomfort or pain after the procedure, on the injection days as well as the day after.

### Single cell dissociation

The embryos were removed from the uterus, sacrificed, and placed into a petri dish containing Hank’s Balanced Salt Solution (HBSS) on ice. Embryos were screened for expression of tdTomato or EGFP using a benchtop fluorescence stereoscope. Limbs, liver, and the ventral skin were removed. Body regions, primarily lower trunk, middle trunk, and cranial regions, were dissected separately and placed into a small Petri dish containing 2.5 mL of a cell dissociation buffer consisting of 2mg/mL Collagenase P (Roche, Sigma-Aldrich), 1X TrypLE, and 1uL/mL DNASe I, fully dissolved in HBSS (without phenol red, including Ca^2+^ and Mg^2+^). Tissue was chopped finely with scalpel and forceps under control of stereomicroscope and transferred to 15mL falcon tubes. The tissue pieces were triturated 10X using a transfer pipette, and then 10X using a 1mL pipette. Samples were then incubated for 20 minutes at 37 C in a preheated shaking incubator set at 150 RPM, with the tubes tilted to ∼60 degrees. During this time every 6-8 minutes, the samples were vortexed and triturated using a 1mL pipette. After the tissue clumps disappeared, 12.5 mL of 2% Fetal Bovine Serum (FBS) diluted in ice-cold HBSS (without phenol red, without Ca^2+^ and Mg^2+^, H6648 SIGMA) was added. The samples were centrifuged in a cold centrifuge (4C) for 15 minutes at 500 g. The supernatant was carefully removed and the pellet was resuspended in 1mL HBSS + 2% FBS and kept on ice thereafter. After centrifugation and resuspension, the cells were filtered into round-bottom polypropylene test tubes (Falcon) using a cell strainer with a 40 micrometer woven polyethylene terephthalate mesh (pluriSelect pluriStrainer SKU 43-10040-40) for downstream cell sorting.

### FACS (Fluorescence activated cell sorting)

FACSDiva software (BD), connected to FACSAria Fusion or FACSAria III flow cytometers (BD) equipped with 100 micrometer nozzle and 1.5 neutral density filter, were used to isolate the fluorescently labeled cells. During sorting the samples were refrigerated to 4 C and regularly agitated by spinning. The event rate was kept below 4000 events/second. Debris, dead cells and doublets were excluded by gating forward-scatter and side-scatter area versus width. Log EGFP and tdTomato fluorescence was detected by excitation with 488 nm and 561 nm lasers, with emissions filtered by 530/30 nm and 610/20 nm bandpass filters, respectively. FP-negative cells served as an internal negative control for FACS sorting. Sorting precision mode was set to “four-way purity”. FP-positive cells were sorted directly into Low-binding tubes containing 50uL of HBSS + 2% FBS. For each sample, all the sorted FP^+^ cells were centrifuged in a cooled centrifuge at 500g for 5 min and resuspended for cell counting with a hemocytometer. From FACS the proportion of labeled cells was ∼5% on average. Because tracing every cell in the embryo across organogenesis would not be feasible (because of the large number of cells in a single E13.5 embryo) we prioritized measuring fewer but more “complete” clones. We observe consistent cell types and clone types across embryos, indicating that a broad range of accessible progenitor types was sampled in all experiments.

### Single cell transcriptomics

Cells were loaded into the Chromium machine (aiming for ∼10,000 cells per reaction) using the V3.1 single-cell reagent kit (10X Genomics). Following GEM creation and cell lysis, cDNA was synthesized by reverse transcription, then amplified by PCR for 12 cycles as per the manufacturer’s protocol (10X Genomics). A portion of the amplified cDNA was used to generate Gene expression libraries and then indexed using the Single Index Plate T Set A. CloneID enrichment from cDNA libraries was additionally performed based on ^15^ to increase the retrieval of the barcodes by ∼5-10%. Briefly, two nested PCR reactions were performed using Q5 error-correcting DNA polymerase to amplify the CloneID-containing region, and a third reaction was used for sample indexing using the Single Index Plate T Set A. In between each reaction, amplicons were purified by 0.8X AMPure beads using a DynaMag Magnetic separator. Indexed GE and CloneID-enrichment libraries were quantified using BioAnalyzer and/or Qubit, and then pooled prior to Illumina sequencing on a NovaSeq 6000 S2-200 v1.5 flow cell with read setup 28-8-0-166.

### Confetti lineage tracing experiment

Transgenic mouse line R26R-Confetti^37,55^, homozygous for the confetti allele, was crossed with Sox10-CreERT2 or PLP-CreERT2 mice. At embryonic day E9 they were injected with 0.02 mg/g tamoxifen (TM) intraperitoneally. Approximately 42 h post TM injection, the pregnant mice were sacrificed and embryos still in the uterus were collected in PBS on ice. Then, under the stereo microscope (ZEISS Discovery V8) and in PBS, the muscle layers of the uterus were ripped carefully with micro-dissecting forceps for the embryos to be extracted. Cre-positive R26R-Confetti embryos were identified on the basis of multicolor fluorescence using a stereo microscope fluorescence adapter. E10.5 R26R-Confetti Cre^+^ mouse embryos were processed to be analyzed histologically or in whole mount. Embryos were fixed in 4% paraformaldehyde (PFA) solution in phosphate buffer saline (PBS) overnight at 4C on rotation. Afterwards, fixed samples headed for histological analysis were washed three times in 1x PBS for 5 min each and were immersed in 30% sucrose in PBS overnight at 4oC on rotation. The next day, they were embedded in Optimal Cutting Temperature (OCT)-compound on dry ice and stored at -20C until cryosectioning. An NX70-Epredia cryostat was used to cut sections with a 14μm thickness which were serially collected onto Superfrost® Plus slides (Thermo Scientific, J1800AMNZ/ground 90°). Prior to storing at -20C, they were left to dry on the bench for 2 hours. Fixed embryos headed for whole mount analysis were washed in PBS before immersed in CUBIC-Reagent I clearing solution for 1 day at 37 C, gently shaking. Then, they were carefully washed with PBS at room temperature (RT) prior to being immersed in CUBIC Reagent II clearing solution at RT for approximately 6 h. The embryos should be totally transparent in order to be placed onto glass bottom dishes (MatTek Corporation) for imaging. The imaging media is CUBIC-Reagent II. This procedure was also used for the whole mount imaging short-term E8.5 lentiviral transductions in E10.5 embryos, except the samples were imaged using a light sheet microscope (Zeiss Z.1) equipped with a 5X objective. Zen Black software was used during acquisition, and manual segmentation in Bitplane Imaris software (v9) was used to reveal the clonal labeling patterns beneath the skin.

### Immunohistochemistry

For histological analysis, tissue sections on slides were submerged in Target Retrieval Solution (DAKO, S1699) for antigen retrieval and brought to boiling point using a microwave oven. The solution was then left to cool down for 40-60 min. Sections were washed in PBS containing 0.1% Tween-20 (PBST). A hydrophobic barrier was created with Super PAP Pen (Invitrogen, 008899) surrounding the tissue. The sections were incubated in a humidified chamber overnight at 4 C with primary antibodies diluted in PBST. The next day, sections were washed in PBST and incubated with secondary antibodies diluted in PBST for 45-90 min. Afterwards, slides were washed with PBST, excessive liquid was removed and the slides were mounted using Mowiol (Merck, 81381) as mounting medium. Primary antibodies that were used against mouse at the indicated dilutions: goat anti-Sox10 (1:800, R&D systems, AF2864), mouse anti-neurofilament medium chain (2H3) (1:100, DSHB, AB_2314897), goat anti-CD31 (1:400, R&D systems, AF3628), mouse anti-Myosin Heavy Chain (MyHC) (1:50, R&D systems, MAB4470), mouse anti-Tyrosine Hydroxylase (TH) (1:1000, Sigma-Aldrich, T2928), chicken anti-GFP (1:500, Aves Labs, GFP-1020), rabbit anti-tRFP (1:500, Invitrogen, PA5-79658), rabbit anti-Sox9 (1:500, Millipore). Primary antibodies were detected with Alexa Fluor 405-, 488-, 555- and 647-conjugated secondary antibodies produced in donkey against mouse, rabbit, goat or chicken antibodies (1:1,000, Molecular Probes, Thermo Fisher Scientific).

### Microscopy and image analysis

Confocal microscopes were used for the imaging of either tissue sections or whole mount samples. Whole mount imaging of cleared E13.5 transduced embryos was performed using light sheet microscopy on a Zeiss Z.1 microscope using a 5X objective. For the Sox10-CreERT2 Confetti labeling, the ZEISS LSM 980 with Airyscan 2 confocal was used due to its capacity to effectively distinguish the overlapping emissions between GFP and YFP. The images were acquired at a 20x objective and 2056x2056 resolution with the use of a z-stack. The imaging of the sections from lentiviral-injected samples was conducted at ZEISS LSM 800 and 800 Airy with a 20x objective and 1024x1024 resolution. ImageJ 1.53t software was used for the analysis of tissue sections. The different color combinations in Sox10-CreERT2 Confetti sections were counted manually with the “Cell Counter” plugin. The determination of DRG units was based on the number of cells in each DRG per embryo. The lentiviral-injected labeled cells in tissue sections were counted with the Cell Counter plugin in compartments of interest like DRG and NT. Bitplane Imaris version 10.0 software was used for the analysis of whole Sox10-CreERT2 Confetti embryos. The four channels were first masked based on the “Surface” to remove background, and a second surface mask was used to distinguish double positive cells.

### scRNA-seq Data Preprocessing and Clone Identification

In the course of the experiments, one to three libraries were generated for each sample: (1) 10x v3 Gene Expression, and (2-3) viral barcode-containing cDNA libraries (two in the case of double injection). Detailed descriptions of each sample, corresponding libraries, and FP sequences are provided in the Github repository of this project (please see Code Availability Statement). Reads from experiments (1) and (2) were aligned to a chimeric mm39 and FP genome reference using Cell Ranger 7.0.1. Subsequently, Gene Expression matrices underwent additional filtering based on sequencing depth, UMI distribution per cell, genes per cell, and the percentage of mitochondrial expression per cell. Specific thresholds are detailed in the Github repository (Code Availability Statement). Clonal barcodes were identified using an algorithm suggested in^15^ with modifications available at https://github.com/serjisa/TREX.modified. Firstly, barcode sequence corrections were performed individually for each cell, enhancing robustness against artificial merging but slightly increasing clonal dropout rates. Secondly, cells were grouped into clones only if they shared identical sets of viral barcodes, precluding the merging of cells with different barcodes. Clonal identification was performed simultaneously on all samples corresponding to the same mouse. Doublet filtering was conducted based on gene expression and clonal information before grouping cells into clones. Scrublet v.0.2.3 ^73^ with manual thresholds was employed for manual doublet filtering. Additionally, cells with two or more viral barcodes, where such combinations existed in only one cell, were considered doublets. Despite these measures, a few clusters likely representing doublets based on marker expression were observed in the final embedding; these were retained in the analysis and labeled as “Doublets.” Reference gene expression embeddings were constructed using scanpy v.1.9.3^74^ in several steps. Gene expression embeddings from all control samples (both head and trunk) were created to identify major cell type domains (Mesenchyme, Neurons, and Other cells). Genes present in fewer than five cells were filtered out. Subsequently, counts per cell were linearly transformed to sum to 10,000 counts per cell, followed by log-transformation of (counts + 1)^75^. The top 3,000 highly variable genes were selected, and PCA was performed on gene-wise scaled expressions. Using the top 30 PCs, dataset horizontal integration was performed with the harmonypy v.0.0.5 package^76^. These corrected PCs were used to construct a kNN graph based on Euclidean distances, followed by community detection with the Leiden algorithm^77^ (leidenalg v.0.9.1) and visualization with UMAP^78^ (umap-learn v.0.5.3). These processing steps were repeated for each of the larger cell subclusters in the head and trunk individually, with an additional step to mitigate the effects of the cell cycle on clustering results. Cell cycle phases were assigned to each cell as described in^79^, and a “reference” was built with cells in interphase. This reference was used for label and coordinate transfer using Symphony^80^ (symphonypy v.0.2). The joint latent representation post-assignment was used for further analysis.

### Lineage Relationships and clone2vec Algorithm

For plotting correlational relationships between cell types, Pearson correlation-based hierarchical clustering was employed. To create clonal latent embeddings, only clones with at least 10 cells were selected. Cells from these clones, termed “whitelisted cells,” were used to find 15 nearest neighbors among the whitelisted cells based on batch-corrected Harmony coordinates using the pynndescent v.0.5.7 package. Each whitelisted cell and its 15 nearest neighbors formed a “bag of clones,” consisting of 15 pairs of context for the target clone. Using a Skip-Gram architecture^19^, a neural network was trained to predict clonal labels of context cells based on the clonal label of the target cell. Clonal labels were one-hot-encoded, and a fully connected neural network with one hidden layer (size = 10) was trained with a negative log-likelihood loss. The matrix describing the linear transformation between the input and hidden layers was used as the resulting embedding, capturing similarities between clones. This embedding enabled the construction of a kNN graph of clones, community detection, and UMAP visualization. Clonal embeddings were constructed for both E7.5 and E8.5 clones simultaneously. Cells with multiple clonal labels from E7.5 and E8.5 were duplicated prior to “bag of clones” construction to count them twice with different clonal labels. For per-clone gene expression analysis, the expression of each cell within a clone was averaged to obtain the final per-clone expression. For embedding exploratory purposes, PCA was performed on the clone2vec embedding space, and gene expression was used to identify associations with clone2vec PCs. Cell cluster names were manually determined by expression of marker genes found in the literature. In case of mesenchymal cells for which the exact identity was unclear, clusters were given names based on transcriptional proximity to well-known cluster or cluster combinations (an example of this would be ‘osteochondral mesenchyme’ situated adjacent to chondrocyte and osteoblast clusters). Then, clonal cluster names were manually assigned by inspection of the cell states and cell types that were present in them, and literature searches for embryonic progenitor cell types that would give rise to each of those cell type ensembles. This information was used in combination with analysis of the gene expression patterns indicating anatomical position, and spatial validation in mouse embryos by mapping clone type-specific gene signatures onto publicly available Stereo-seq datasets^81^.

### Gene Importance Analysis

To identify genes associated with clonal composition, we built models to predict either clone2vec latent embedding coordinates or cell type proportions within each clone. For this purpose, we used a gradient boosting model, as its applicability for gene regulatory network (GRN) inference has been demonstrated^82^. Specifically, we employed the relatively overfitting-robust CatBoost algorithm^83^, following the strategy described below. First, we created pseudo-bulk profiles for each clone within each cell type to mitigate learning of cell type–specific gene expression. To further reduce this effect, we included cell type as a categorical predictor alongside gene expression. Clones were then split into two subsets: a training set (80% of clones) and a validation set (20%) to monitor overfitting during training. The validation set was selected using a geometric sketching approach^84^, choosing clones that best represent the clone2vec space. CatBoost regression (for latent coordinate prediction) or classification (for clone composition prediction) was applied using MultiRMSE or MultiCrossEntropy loss functions on both the training and validation sets. Training was stopped if the validation loss did not improve for more than 100 iterations, and the best model was selected for further analysis. The final model was used to estimate feature importance for each prediction across the dataset using Shapley values^85,86^. To quantify the overall importance of each gene in clonal behavior, we averaged the L2-norm of the SHAP vectors from the clone2vec model across all generated pseudo-bulks. To investigate gene associations with cell types, we extracted SHAP value modules from the cell type model and assigned them the sign of their correlation with gene expression. This sign correction allowed us to distinguish between positive and negative markers of cell type proportions. Thus, the clone2vec model captures the overall variance in clonal behavior, while the cell type model provides a mechanistic interpretation of gene effects at the cell type level. Both models were trained using the same train/validation split for consistency. For visualization, sign-corrected SHAP magnitudes from each cell type can be min-max normalized with respect to their sign.

### Cartilage Trajectory Analysis

To reconstruct differentiation trajectories within individual clonal clusters, cells were re-embedded, and clusters linked to the cartilage/bone trajectory were selected from clonal clusters 2 and 8. Velocity analysis with UniTVelo (v.0.2.5.2) confirmed the differentiation directionality within the selected cells. Bifurcation points were identified using the scFates package (v.1.0.7)^87^. Pseudotime was inferred using the diffusion pseudotime approach, and cells with pseudotime value less than 0.3 were considered as progenitors^88^. Alignment between two differentiation trajectories was performed with the DTW Python package (v.1.3.0)^89,90^. The significance of the association between gene expression and pseudotime was determined using GAM fitting and LRT, comparing models with and without pseudotime-associated splines. Differences in associations with pseudotime across trajectories were identified by comparing GAMs with and without clonal cluster assignment as a covariate. Genes significantly associated with pseudotime (p < 10^−25^) and not differing between clonal clusters (FDR > 0.05) were considered “common”, at the same time genes with significant lineage association (FDR < 10^−5^) and fold-change > 1 between lineages were considered as “unique”. Bone-related signature was gotten by differential expression analysis between cartilage and bone in clonal cluster 2 (top-20 highly expressed in cartilage trajectory genes with FDR < 10^−20^ and logFC > 4) using Student’s t-test. Analysis of “early” genes was performed with standard scFates pipeline^87^. Transcription factor (TF) activity inference was conducted using pyscenic (v.0.12.1)^30^ with arboreto (v.0.1.6). Differential activity analysis between clonal clusters followed the same procedure. Graph of the potential regulation was gotten from cisTarget step of pyscenic protocol. In all cases signature scores were calculated with sc.tl.score_genes scanpy function.

### Spatial Validation of Gene Expression Signatures

Stereo-seq data from^81^ was used to identify the spatial localization of gene expression signatures. Gene signatures were identified using Student’s t-test between groups of cells (e.g., between different clonal clusters or within the same cell type across different clonal clusters) with a minimum log2FC of 2. Expression of gene signatures within one cell type was visualized as the product of gene and cell type signatures. Hox-score was calculated as a min-max normalized weighted sum of expressions of the following Hox-genes: *Hox score*_raw_ = (1 × expr(*Hoxb4*) + 2 × expr(*Hoxb5*) + 3 × expr(*Hoxc6*) + 4 × expr(*Hoxa7*) + 5 × expr(*Hoxc8*) + 6 × expr(*Hoxd9*) + 7 × expr(*Hoxc10*) + 8 × expr(*Hoxd11*) + 9 × expr(*Hoxc11*) + 10 × expr(*Hoxc12*) + 11 × expr(*Hoxc13*)) / 66; *Hox score* = (*Hox score*_raw_ – min(*Hox score*_raw_)) / (max(*Hox score*_raw_) – min(*Hox score*_raw_)).

### Perturbation Analysis

Perturbational screening experiments were mapped onto the control reference via Symphony, as previously described (separately for cycling and non-cycling cells). Each clone from the perturbational experiment was assigned to a gRNA based on the most abundant gRNA combination within the clone. Clones with multiple gRNAs were excluded from further analysis. New clonal embeddings were constructed as described in the Lineage Relationships and Clone2vec Algorithm section. Differential compositional testing between clones in different clonal clusters was conducted using scCODA^91^ to analyze clone composition in the entire clonal embedding rather than individual cells within different cell types. An independent embedding focusing on NC-related clones (with at least three cells) was created using both control and perturbed cells. Compositional testing was performed for perturbations with multiple replicates using beta regression and LRT with FDR multiple testing correction.

## Code Availability

All code that was used in the current study is readily available on Github: (https://github.com/kharchenkolab/clonal-atlas-paper/). The clone2vec Python package is also available on Github: (https://github.com/kharchenkolab/scLiTr). Additional documentation for the scLiTr package can be found here: (https://sclitr.readthedocs.io/). The modified TREX pipeline is available here: (https://github.com/serjisa/TREX.modified). Also, the code for the clones2cells app is available via this link: (https://github.com/serjisa/clones2cells_app)

## Data Availability

Raw data has been submitted to the NCBI Gene Expression Omnibus database (GEO) and is accessible via the following link: (https://www.ncbi.nlm.nih.gov/geo/query/acc.cgi?acc=GSE269395). Data has also been uploaded to Zenodo: (https://zenodo.org/records/15397247). The data can be browsed via the clones2cells app via this link: (https://clones2cells.streamlit.app). Alternatively, browse the datasets on our CELLXGENE single-cell dataset repository: (https://adameykolab.hifo.meduniwien.ac.at/cellxgene_public/filecrawl/2024_Erickson_Isaev)

