## Extended Data Figures 1-17 for "Unbiased profiling of multipotency landscapes reveals spatial modulators of clonal fate biases"

**Extended Data Figure 1. Overview of the injections used in the general dataset.**

A breakdown of all samples injected with control barcoded virus in the study used to make the clonal atlas of the E13.5 embryo. Left: experimental timeline, including the time points of injection and harvesting, as well as the volume of the injected virus. Right: the body regions harvested from each sample, as well as notes about specific unique features of some samples.

In addition to the embryos shown in the schematic, we also generated the following TREX-scRNAseq datasets: Four injected embryos related to perturb-TREX experiments (Figures 5-6). Four injected embryos to validate the NCC-MP clone population in E14 lower trunk and tail tissue (Extended Data Figure 13).

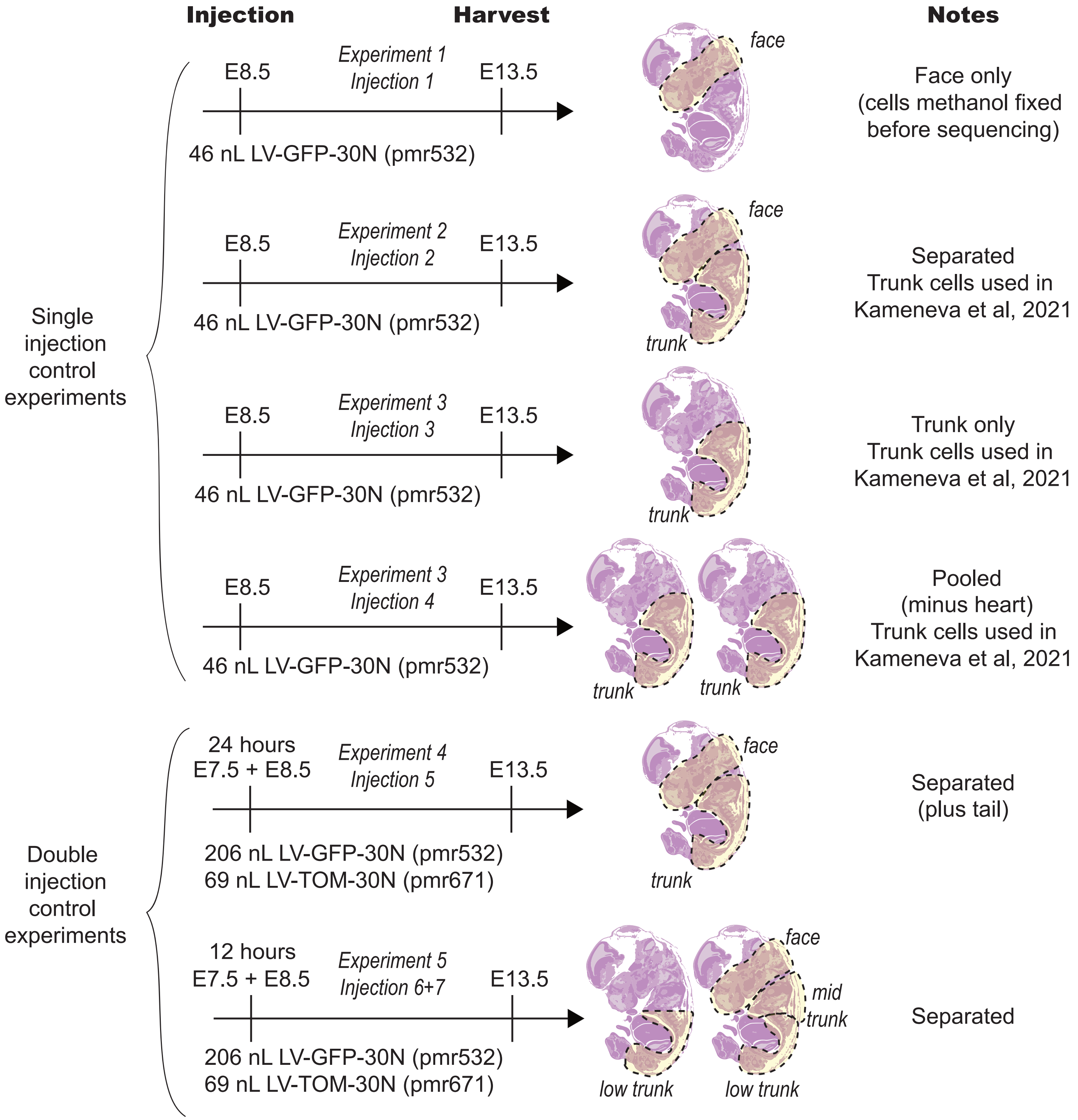

Injection
Harvest
Notes
Single injection control experiments
E8.5
Experiment 1
Injection 1
E13.5
46 nL LV-GFP-30N (pmr532)
face
Face only (cells methanol fixed before sequencing)
E8.5
Experiment 2
Injection 2
E13.5
46 nL LV-GFP-30N (pmr532)
face
trunk
Separated Trunk cells used in Kameneva et al, 2021
E8.5
Experiment 3
Injection 3
E13.5
46 nL LV-GFP-30N (pmr532)
trunk
Trunk only Trunk cells used in Kameneva et al, 2021
E8.5
Experiment 3
Injection 4
E13.5
46 nL LV-GFP-30N (pmr532)
trunk
trunk
Pooled (minus heart) Trunk cells used in Kameneva et al, 2021
Double injection control experiments
24 hours
E7.5 + E8.5
Experiment 4
Injection 5
E13.5
206 nL LV-GFP-30N (pmr532)
69 nL LV-TOM-30N (pmr671)
face
trunk
Separated (plus tail)
12 hours
E7.5 + E8.5
Experiment 5
Injection 6+7
E13.5
206 nL LV-GFP-30N (pmr532)
69 nL LV-TOM-30N (pmr671)
face
mid trunk
low trunk
low trunk
Separated

**Extended Data Figure 2. QC of the scRNAseq data and clone assignment.**

Description of the quality control for scRNAseq and clone assignment for craniofacial **(a-d)** and trunk **(e-h)** samples. **(a, e)** the highlighted body part indicates the origin of the analyzed cells. **(b, f)** Histograms show the distribution of UMIs (left), genes (middle) and mitochondrial UMIs (right) per cell. Number of cells per clone is quantified by a histogram (left panels) and a cumulative chart (right panels) for craniofacial **(c-d)** and trunk **(g-h)** clones labeled from E7.5 **(c, g)** versus E8.5 **(d, h)** until E13.5.

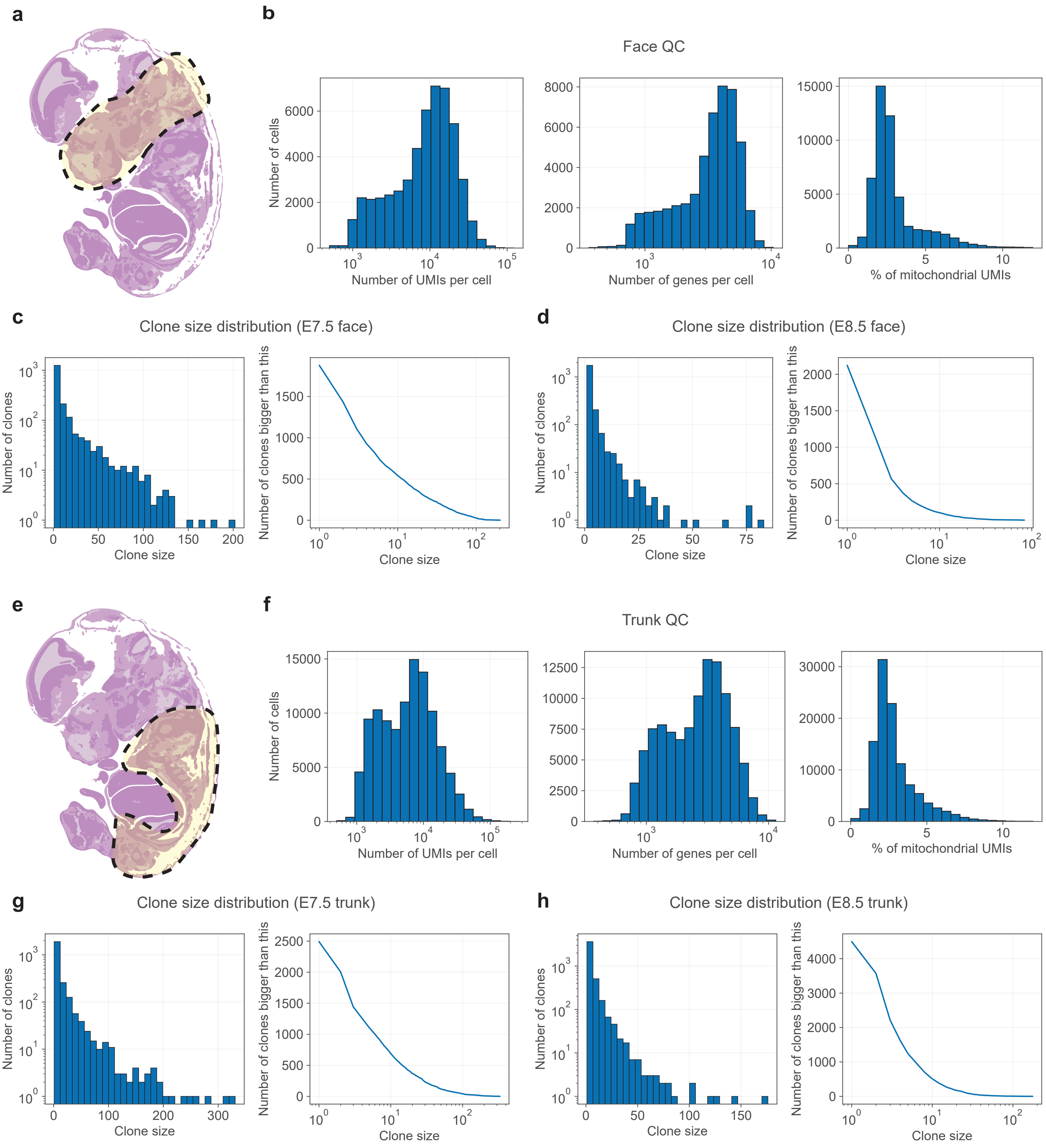
a
b
Face QC
Number of cells
Number of UMIs per cell
Number of genes per cell
% of mitochondrial UMIs
c
Clone size distribution (E7.5 face)
Number of clones
Clone size
Number of clones bigger than this
d
Clone size distribution (E8.5 face)
e
f
Trunk QC
g
Clone size distribution (E7.5 trunk)
h
Clone size distribution (E8.5 trunk)

**Extended Data Figure 3. Examples of successful and sparse clonal transduction in the trunk after optimization.** Whole mount light sheet imaging of full trunks of mouse embryos injected with LV-GFP, and LV-RFP at E8.5, and harvested at embryonic day 10.5. The skin is segmented away manually to visualize neural crest clones and polyclones in whole mounts as a maximum intensity projection (left and middle columns). A full transverse section (right column) shows the labeled skin as well as internally labeled cells in the neural crest and neural tube. The embryos were injected with various dilutions of the virus, with effective doses at 2nL **(a)** 5nL **(b)** and 12 nL **(c)** of virus. Scale bar 200 µm except: a, top left panel scale = 500 µm, (a-b); right panel scale = 150 µm.

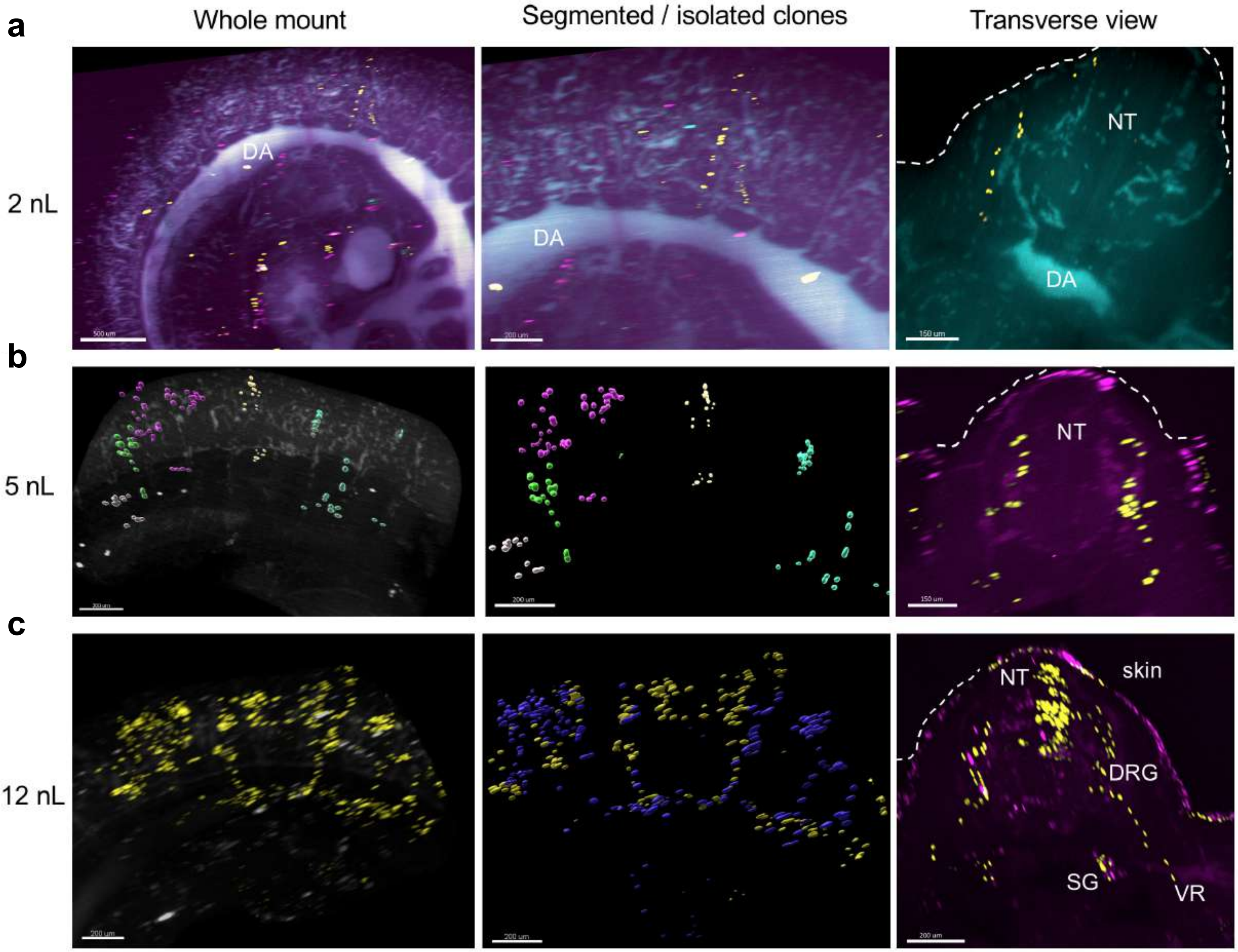
a
Whole mount
Segmented / isolated clones
Transverse view
2 nL
DA
DA
NT
DA
b
5 nL
NT
c
12 nL
NT
skin
DRG
SG
VR

**Extended Data Figure 4. Broad distribution of labeled cells in barcoded embryos.** Whole mount light sheet imaging of full trunks of mouse embryos injected with LV-GFP, and harvested at embryonic day 13.5, co-stained with tissue-type specific labels. **(a)** Transduced embryo co-stained with anti-tyrosine hydroxylase (sympathetic neurons), 2H3 (anti-neurofilament), and anti-GFP. **(b)** Example of transduced embryo co-stained with anti-Sox9 (cartilage), anti-GFP, and 2H3. Autofluorescence is shown in dark blue for visualizing other body structures. Scale bar = 1000 μm.

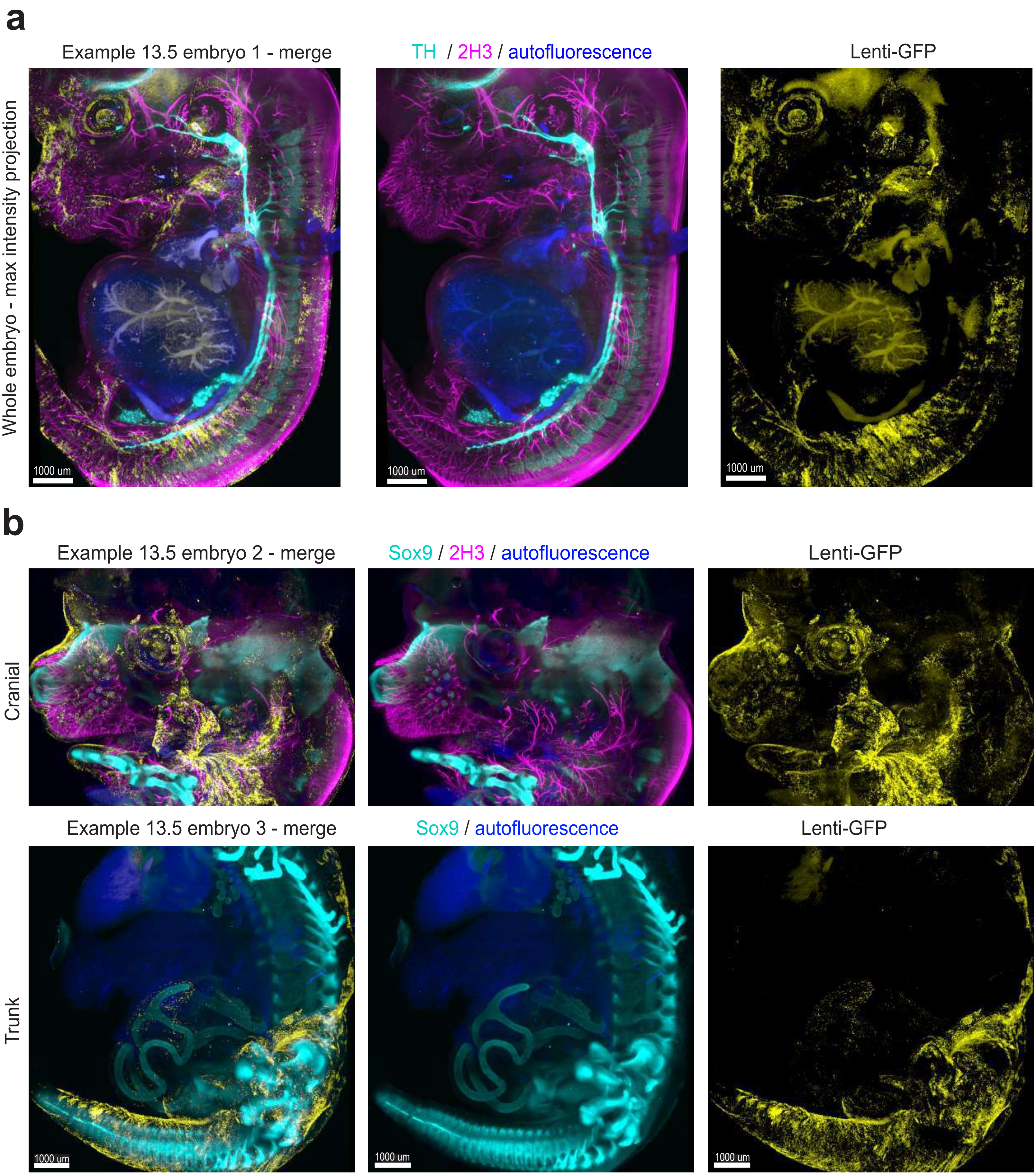
a
Example 13.5 embryo 1 - merge
TH / 2H3 / autofluorescence
Lenti-GFP
Whole embryo - max intensity projection
1000 um
1000 um
1000 um
b
Example 13.5 embryo 2 - merge
Sox9 / 2H3 / autofluorescence
Lenti-GFP
Cranial
Example 13.5 embryo 3 - merge
Sox9 / autofluorescence
Lenti-GFP
Trunk
1000 um
1000 um
1000 um

**Extended Data Figure 5. Trunk cell types recovered.**

Left column, UMAP embeddings of the major categories of cell types observed in E13.5 embryonic trunk in this study, including **(a)** mesenchymal cells, **(b)** peripheral nervous system cells, **(c)** central nervous system cells, and **(d)** all other cell types that formed separate clusters in the overall trunk dataset. Right column, dot plots display the top few gene markers for each cell type cluster. The size of the dot indicates the fraction of cells in the group expressing the gene, and dot color indicates mean expression of that gene.

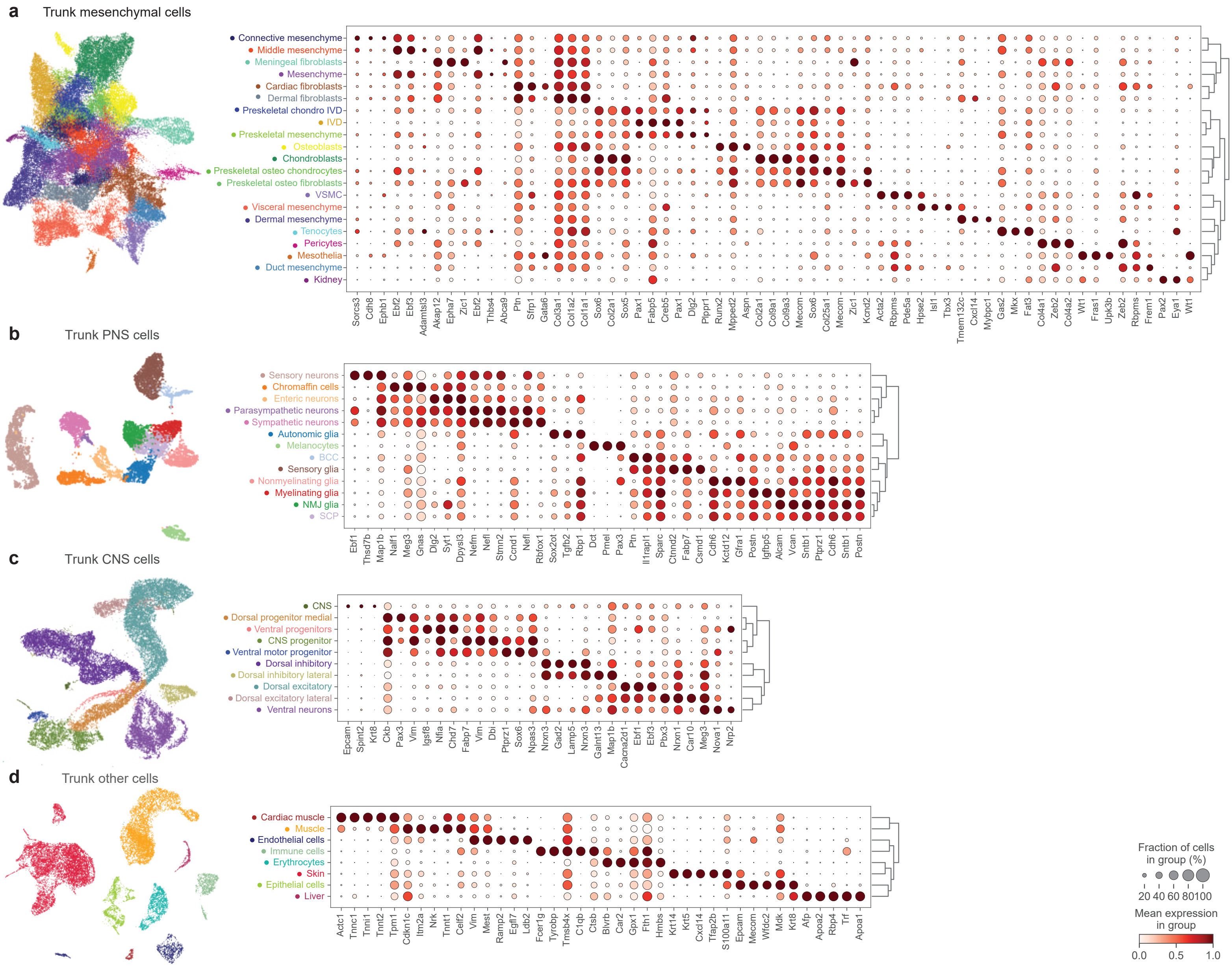

a
Trunk mesenchymal cells
Connective mesenchyme
Middle mesenchyme
Meningeal fibroblasts
Mesenchyme
Cardiac fibroblasts
Dermal fibroblasts
Preskeletal chondro IVD
IVD
Preskeletal mesenchyme
Osteoblasts
Chondroblasts
Preskeletal osteo chondrocytes
Preskeletal osteo fibroblasts
VSMC
Visceral mesenchyme
Dermal mesenchyme
Tenocytes
Pericytes
Mesothelia
Duct mesenchyme
Kidney
Sorcs3 Cdh8 Ephb1 Ebf2 Ebf3 Adamtsl3 Akap12 Epha7 Zic1 Ebf2 Thbs4 Abca9 Ptn Sfrp1 Gata6 Col3a1 Col1a2 Col1a1 Sox6 Col2a1 Sox5 Pax1 Fabp5 Creb5 Pax1 Dlg2 Plppr1 Runx2 Mpped2 Aspn Col2a1 Col9a1 Col9a3 Mecom Sox6 Col25a1 Mecom Zic1 Kcnd2 Acta2 Rbpms Pde5a Hpse2 Isl1 Tbx3 Tmem132c Cxcl14 Mybpc1 Gas2 Mkx Fat3 Col4a1 Zeb2 Col4a2 Wt1 Fras1 Upk3b Zeb2 Rbpms Frem1 Pax2 Eya1 Wt1
b
Trunk PNS cells
Sensory neurons
Chromaffin cells
Enteric neurons
Parasympathetic neurons
Sympathetic neurons
Autonomic glia
Melanocytes
BCC
Sensory glia
Nonmyelinating glia
Myelinating glia
NMJ glia
SCP
Ebf1 Thsd7b Map1b Nalf1 Meg3 Gnas Dlg2 Syt1 Dpysl3 Nefm Nefl Stmn2 Ccnd1 Nefl Rbfox1 Sox2ot Tgfb2 Rbp1 Dct Pmel Pax3 Ptn Il1rapl1 Sparc Ctnnd2 Fabp7 Csmd1 Cdh6 Kctd12 Gfra1 Postn Igfbp5 Alcam Vcan Sntb1 Ptprz1 Cdh6 Sntb1 Postn
c
Trunk CNS cells
CNS
Dorsal progenitor medial
Ventral progenitors
CNS progenitor
Ventral motor progenitor
Dorsal inhibitory
Dorsal inhibitory lateral
Dorsal excitatory
Dorsal excitatory lateral
Ventral neurons
Epcam Spint2 Krt8 Ckb Pax3 Vim Igsf8 Nfia Chd7 Fabp7 Vim Dbi Ptprz1 Sox6 Npas3 Nrxn3 Gad2 Lamp5 Nrxn3 Galnt13 Map1b Cacna2d1 Ebf1 Ebf3 Pbx3 Nrxn1 Car10 Meg3 Nova1 Nrp2
d
Trunk other cells
Cardiac muscle
Muscle
Endothelial cells
Immune cells
Erythrocytes
Skin
Epithelial cells
Liver
Actc1 Tnnc1 Tnni1 Tnnt2 Tpm1 Cdkn1c Itm2a Nrk Tnnt1 Celf2 Vim Mest Ramp2 Egfl7 Ldb2 Fcer1g Tyrobp Tmsb4x C1qb Ctsb Blvrb Car2 Gpx1 Fth1 Hmbs Krt14 Krt5 Cxcl14 Tfap2b S100a11 Epcam Mecom Wfdc2 Mdk Krt8 Afp Apoa2 Rbp4 Trf Apoa1
Fraction of cells in group (%)
20 40 60 80 100
Mean expression in group
0.0 0.5 1.0

**Extended Data Figure 6. Craniofacial cell types recovered.**

Left column, UMAP embeddings of the major categories of cell types observed in E13.5 embryonic craniofacial region in this study, including **(a)** mesenchymal cells, **(b)** nervous system cells, **(c)** all other cell types that formed separate clusters in the overall craniofacial dataset. Right column, dot plots display the top few gene markers for each cell type cluster. The size of the dot indicates the fraction of cells in the group expressing the gene, and dot color indicates mean expression of that gene.

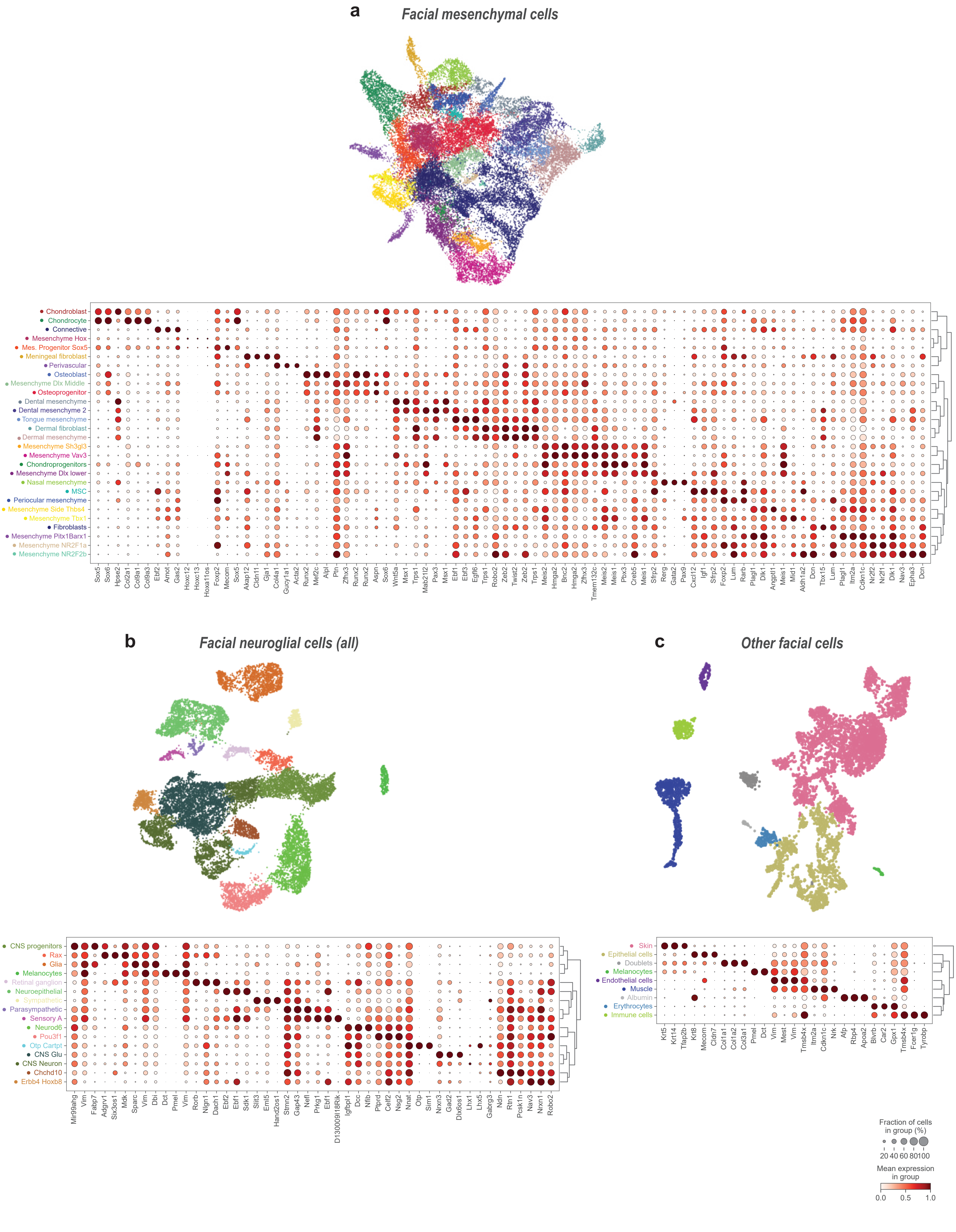

a
Facial mesenchymal cells
Chondroblast
Chondrocyte
Connective
Mesenchyme Hox
Mes. Progenitor Sox5
Meningeal fibroblast
Perivascular
Osteoblast
Mesenchyme Dlx Middle
Osteoprogenitor
Dental mesenchyme
Dental mesenchyme 2
Tongue mesenchyme
Dermal fibroblast
Dermal mesenchyme
Mesenchyme Sh3gl3
Mesenchyme Vav3
Chondroprogenitors
Mesenchyme Dlx lower
Nasal mesenchyme
MSC
Periocular mesenchyme
Mesenchyme Side Thbs4
Mesenchyme Tbx1
Fibroblasts
Mesenchyme Pitx1Barx1
Mesenchyme NR2F1a
Mesenchyme NR2F2b
Sox5 Sox6 Hpse2 Col2a1 Col9a1 Col9a3 Ebf2 Amot Gas2 Hoxc12 Hoxc13 Hoxa11os Foxp2 Mecom Sox5 Akap12 Cldn11 Gja1 Col4a1 Gucy1a1 Acta2 Runx2 Mef2c Alpl Ptn Zfhx3 Runx2 Runx2 Aspn Sox6 Wnt5a Msx1 Trps1 Mab21l2 Pax3 Msx1 Ebf1 Ebf3 Egfl6 Trps1 Robo2 Zeb2 Twist2 Zeb2 Trps1 Meis2 Hmga2 Bnc2 Hmga2 Zfhx3 Tmem132c Meis2 Meis1 Pbx3 Creb5 Meis1 Sfrp2 Rerg Gata2 Pax9 Cxcl12 Igf1 Sfrp2 Foxp2 Lum Rarb Plagl1 Dlk1 Angptl1 Meis1 Mid1 Aldh1a2 Dcn Tbx15 Lum Plagl1 Itm2a Cdkn1c Nr2f2 Nr2f1 Dlk1 Nav3 Epha3 Dcn
b
Facial neuroglial cells (all)
CNS progenitors
Rax
Glia
Melanocytes
Retinal ganglion
Neuroepithelial
Sympathetic
Parasympathetic
Sensory A
Neurod6
Pou3f1
Otp Cartpt
CNS Glu
CNS Neuron
Chchd10
Erbb4 Hoxb8
Mir99ahg Vim Fabp7 Adgrv1 Six3os1 Mdk Sparc Vim Dbi Dct Pmel Vim Rorb Nlgn1 Dach1 Ebf2 Ebf1 Sdk1 Slit3 Eml5 Hand2os1 Stmn2 Gap43 Nefl Prkg1 Ebf1 D130009I18Rik Igfbpl1 Dcc Nfib Ptprd Celf2 Nsg2 Nnat Otp Sim1 Nrxn3 Gad2 Dlx6os1 Lhx1 Lhx5 Gabrg3 Ndn Rtn1 Pcsk1n Nav3 Nrxn1 Robo2
c
Other facial cells
Skin
Epithelial cells
Doublets
Melanocytes
Endothelial cells
Muscle
Albumin
Erythrocytes
Immune cells
Krt5 Krt14 Tfap2b Krt8 Mecom Cldn7 Col1a1 Col1a2 Col3a1 Pmel Dct Vim Mest Vim Tmsb4x Itm2a Cdkn1c Nrk Afp Rbp4 Apoa2 Blvrb Car2 Gpx1 Tmsb4x Fcer1g Tyrobp
Fraction of cells in group (%)
20 40 60 80 100
Mean expression in group
0.0 0.5 1.0

**Extended Data Figure 7. Co-occurrence frequency of major fates among traced cranial clones.** Upset plot of major types of cranial clones resulting from injections either at E7.5 or E8.5 (datasets pooled). Left, set size indicates the number of clones containing each of the major fates. Top; intersection size shows the number of clones containing the specific combination of fates indicated by the connected dots.

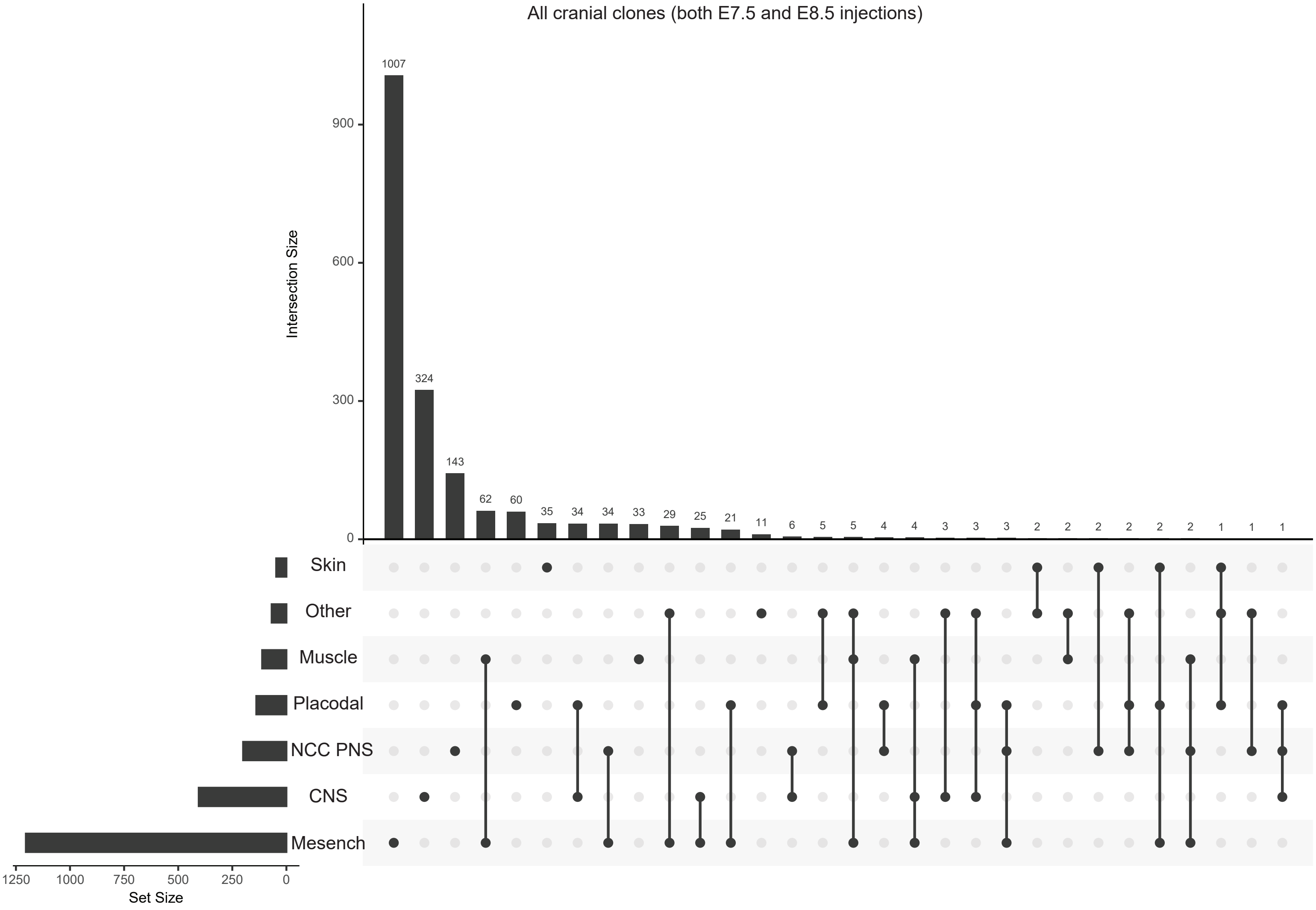

All cranial clones (both E7.5 and E8.5 injections)
Intersection Size
900
600
300
0
1007
324
143
62
60
35
34
34
33
29
25
21
11
6
5
5
4
4
3
3
3
2
2
2
2
2
2
1
1
1
Skin
Other
Muscle
Placodal
NCC PNS
CNS
Mesench
1250
1000
750
500
250
0
Set Size

**Extended Data Figure 8. Heatmaps for lineage correlation.** Heat maps showing Pearson's correlation based on clone sharing across all cell types in craniofacial **(a)** and trunk **(b)** tissue. Top, hierarchical clustering of the cell types based on the lineage correlation.

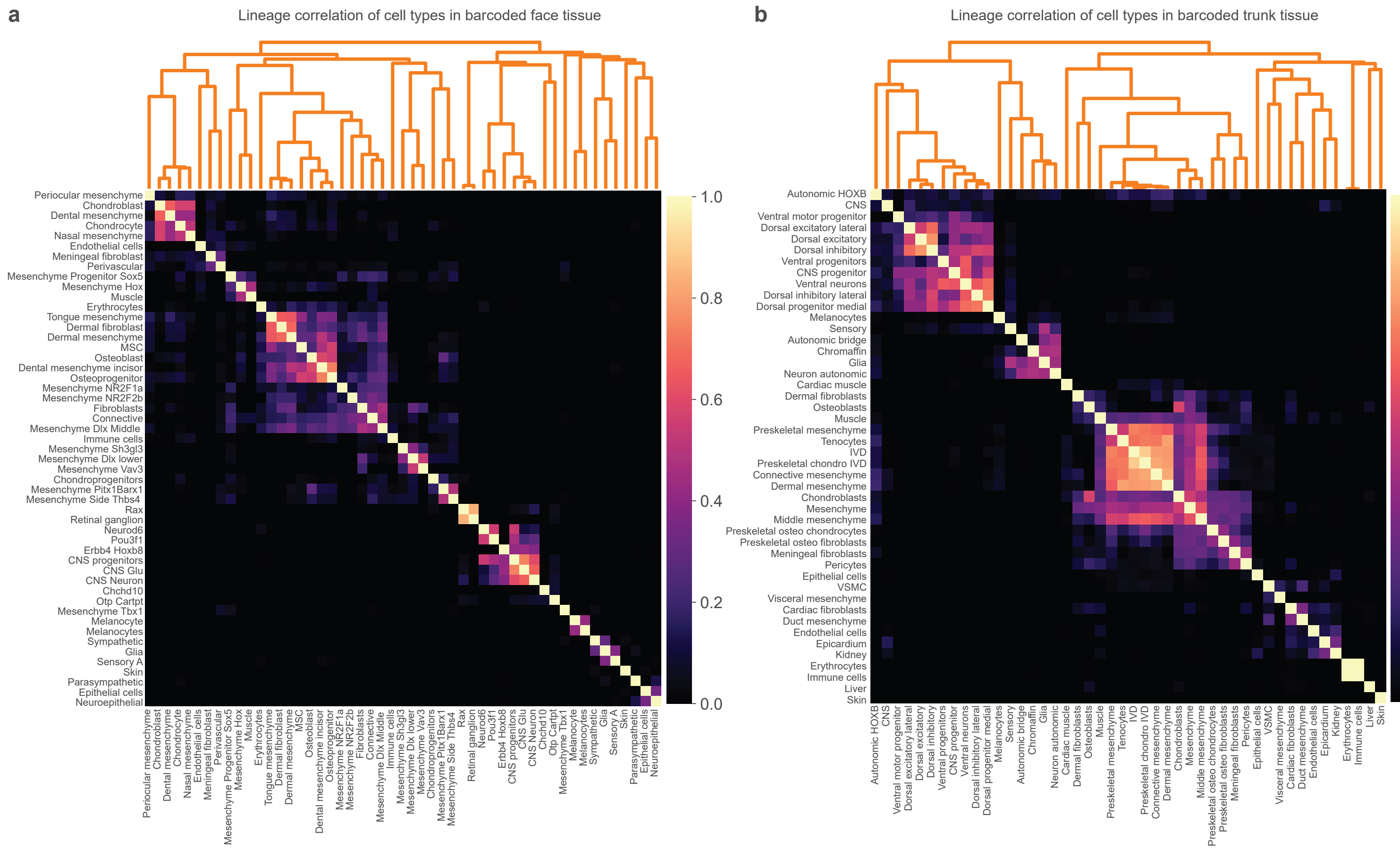
a
Lineage correlation of cell types in barcoded face tissue
Periocular mesenchyme
Chondroblast
Dental mesenchyme
Chondrocyte
Nasal mesenchyme
Endothelial cells
Meningeal fibroblast
Perivascular
Mesenchyme Progenitor Sox5
Mesenchyme Hox
Muscle
Erythrocytes
Tongue mesenchyme
Dermal fibroblast
Dermal mesenchyme
MSC
Osteoblast
Dental mesenchyme incisor
Osteoprogenitor
Mesenchyme NR2F1a
Mesenchyme NR2F2b
Fibroblasts
Connective
Mesenchyme Dlx Middle
Immune cells
Mesenchyme Sh3gl3
Mesenchyme Dlx lower
Mesenchyme Vav3
Chondroprogenitors
Mesenchyme Pitx1Barx1
Mesenchyme Side Thbs4
Rax
Retinal ganglion
Neurod6
Pou3f1
Erbb4 Hoxb8
CNS progenitors
CNS Glu
CNS Neuron
Chchd10
Otp Cartpt
Mesenchyme Tbx1
Melanocyte
Melanocytes
Sympathetic
Glia
Sensory A
Skin
Parasympathetic
Epithelial cells
Neuroepithelial
1.0
0.8
0.6
0.4
0.2
0.0
b
Lineage correlation of cell types in barcoded trunk tissue
Autonomic HOXB
CNS
Ventral motor progenitor
Dorsal excitatory lateral
Dorsal excitatory
Dorsal inhibitory
Ventral progenitors
CNS progenitor
Ventral neurons
Dorsal inhibitory lateral
Dorsal progenitor medial
Melanocytes
Sensory
Autonomic bridge
Chromaffin
Glia
Neuron autonomic
Cardiac muscle
Dermal fibroblasts
Osteoblasts
Muscle
Preskeletal mesenchyme
Tenocytes
IVD
Preskeletal chondro IVD
Connective mesenchyme
Dermal mesenchyme
Chondroblasts
Mesenchyme
Middle mesenchyme
Preskeletal osteo chondrocytes
Preskeletal osteo fibroblasts
Meningeal fibroblasts
Pericytes
Epithelial cells
VSMC
Visceral mesenchyme
Cardiac fibroblasts
Duct mesenchyme
Endothelial cells
Epicardium
Kidney
Erythrocytes
Immune cells
Liver
Skin
1.0
0.8
0.6
0.4
0.2
0.0

**Extended Data Figure 9. Diversity of fate distributions within trunk clonal clusters.** Heat maps organized by clonal cluster (indicated as "leiden") and cell types. Each row shows an individual clone, color-coded according to the proportions dedicated to each cell type (columns). To the right of each heat map, a dendrogram organizes clonal clusters by similarity. **(a)** Clones traced from E7.5 and harvested at E13.5. **(b)** Clones traced from E8.5 and harvested at E13.5.

Diversity of fate distributions within individual trunk clonal clusters

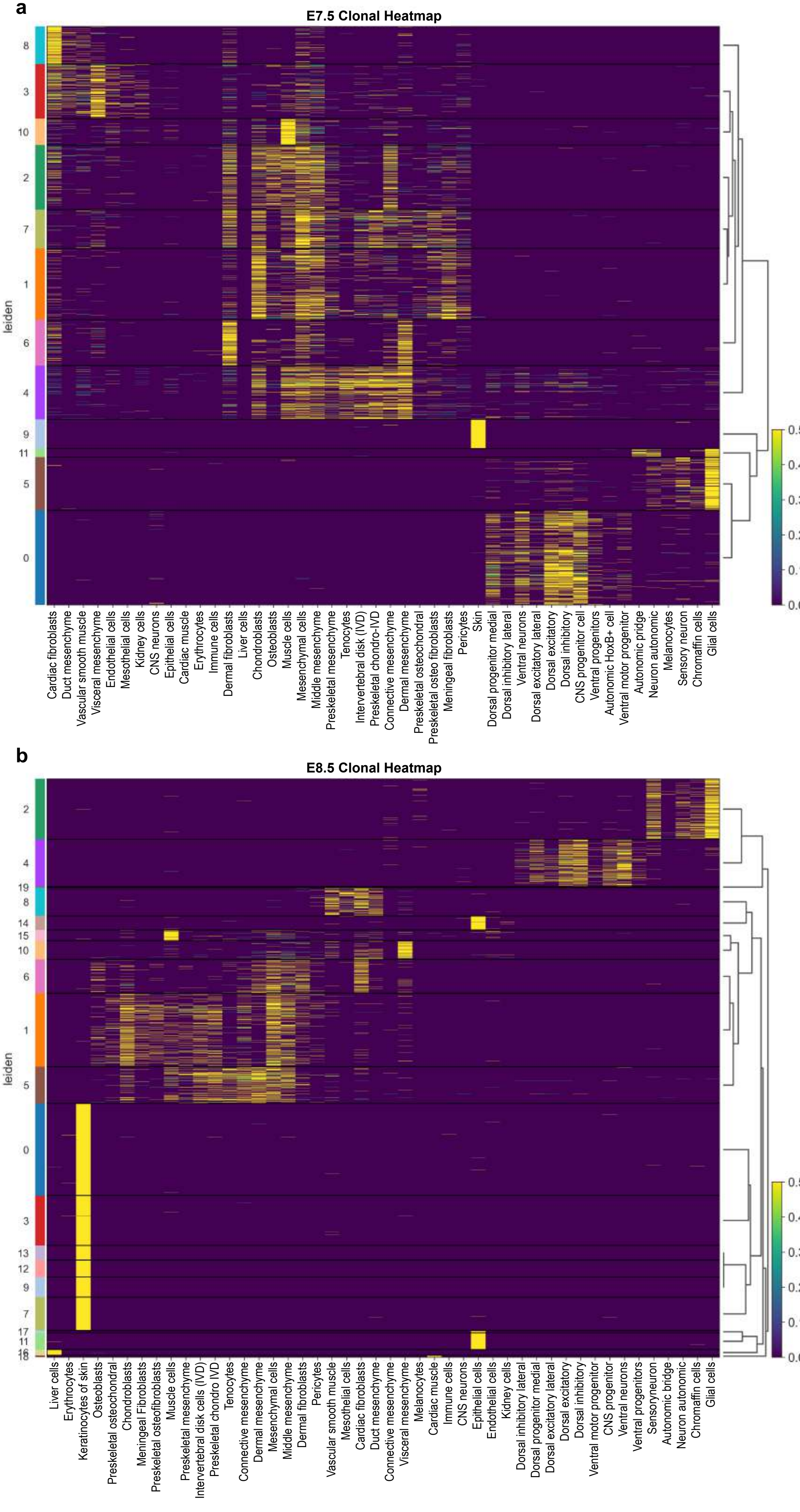

**Extended Data Figure 10. A continuous fate gradient identified in somitic mesodermal clones. (a)** Trunk clone2vec clonal embedding, color coded by discrete (skin, neural crest, and ventral mesenchyme, top) versus contiguous (somite-derived mesenchyme lineages, bottom) selections of clonal clusters.

**(b)** Principal components analysis (PCA) of the same selected clones, color coded as in (a).

**(c-d)** Trajectory inference applied to PCA embedding. (c) Black dots indicate milestones representing intermediate states along the inferred trajectory, and an arbitrary directionality of 'clonal pseudotime' is designated in (d) corresponding to the position of clones in PCA space (e-j).

**(e)** Heat map showing the expression levels of genes expressed by cells assigned to clones, arranged by their position in PCA space (according to the clonal 'pseudotime' value).

**(f-j)** Proportion of cells of the specific mesenchymal fate within each clone, including dermis (f), meninges (g), osteoblasts (h), intervertebral disk cells (i), and chondroblasts (j). Clones are sorted by their position in PCA space (according to the clonal 'pseudotime' values).

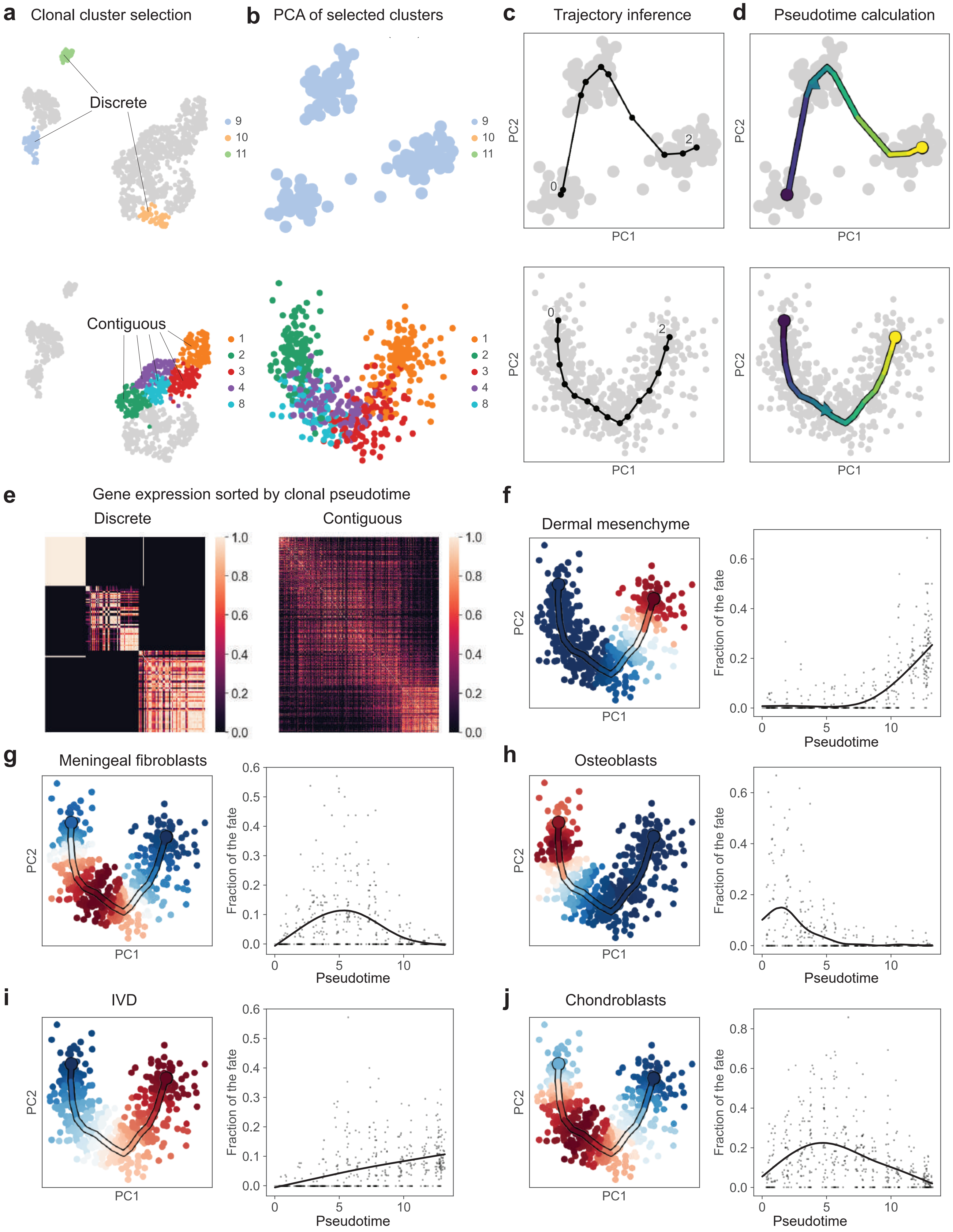
a
Clonal cluster selection
Discrete
Contiguous
9
10
11
1
2
3
4
8
b
PCA of selected clusters
c
Trajectory inference
0
2
PC1
PC2
d
Pseudotime calculation
e
Gene expression sorted by clonal pseudotime
Discrete
Contiguous
1.0
0.8
0.6
0.4
0.2
0.0
f
Dermal mesenchyme
Fraction of the fate
Pseudotime
g
Meningeal fibroblasts
h
Osteoblasts
i
IVD
j
Chondroblasts

**Extended Data Figure 11. Clonal compositional fate gradient in mesenchymal cells is driven primarily by anteroposterior patterning of somites.**

**(a)** Trunk clone2vec clonal embedding, color coded by the somite-derived mesenchyme lineages.

**(b)** Trajectory inference of the selected clones in PCA space (from Extended Data Figure 10d).

**(c)** Scatterplot showing the Hox score of the selected clones, sorted by their position in PCA space (clonal 'pseudotime' value).

**(d)** Feature plot from a reference Stereoseq dataset of an E13.5 mouse embryo, color-coded to illustrate the anatomical mapping of Hox scores.

**(e-h)** Scatter plots showing gene expression levels per clone for genes exhibiting body-wide expression gradients in somitic mesenchymal clones, including *Plagl1* (e), *Col12a1* (f), *Itm2a* (g), and *Hmga2* (h). Clones are arranged by their clonal 'pseudotime' value.

**(i-l)** Scatterplots showing per-clone expression levels of two genes with graded expression profiles, *Itm2a* (i-j) and *Cdkn1c* (k-l), plotted against the per-clone Hox score, in somitic mesenchymal lineages (i, k) versus non-somitic mesenchymal linages (j, l).

**(m)** Scatterplot showing per-clone expression levels of *Mki67* plotted against the per-clone Hox score.

**(n)** Normalized barcode heatmap with hierarchical grouping of clones based on co-occurrence of different fates within them; rows are colored by the cluster identity of the clones (resolution = 1).

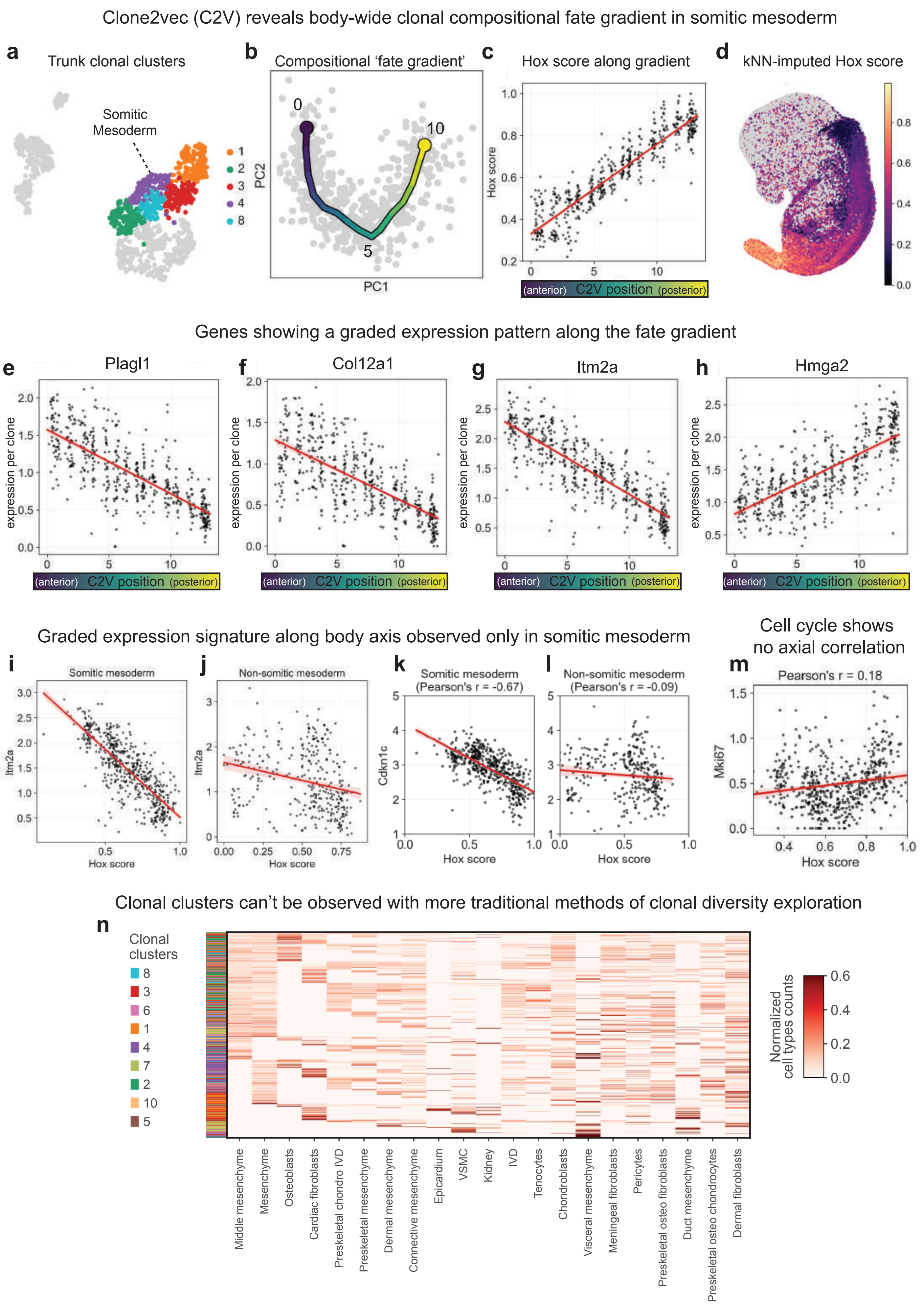

Clone2vec (C2V) reveals body-wide clonal compositional fate gradient in somitic mesoderm
a
Trunk clonal clusters
Somitic Mesoderm
1
2
3
4
8
b
Compositional 'fate gradient'
0
10
5
PC2
PC1
c
Hox score along gradient
Hox score
(anterior) C2V position (posterior)
d
kNN-imputed Hox score
Genes showing a graded expression pattern along the fate gradient
e
Plagl1
f
Col12a1
g
Itm2a
h
Hmga2
expression per clone
(anterior) C2V position (posterior)
Graded expression signature along body axis observed only in somitic mesoderm
i
Somitic mesoderm
Itm2a
Hox score
j
Non-somitic mesoderm
Itm2a
Hox score
k
Somitic mesoderm (Pearson's r = -0.67)
Cdkn1c
Hox score
l
Non-somitic mesoderm (Pearson's r = -0.09)
Hox score
Cell cycle shows no axial correlation
m
Pearson's r = 0.18
Mki67
Hox score
Clonal clusters can't be observed with more traditional methods of clonal diversity exploration
n
Clonal clusters
8
3
6
1
4
7
2
10
5
Normalized cell types counts
Middle mesenchyme
Mesenchyme
Osteoblasts
Cardiac fibroblasts
Preskeletal chondro IVD
Preskeletal mesenchyme
Dermal mesenchyme
Connective mesenchyme
Epicardium
VSMC
Kidney
IVD
Tenocytes
Chondroblasts
Visceral mesenchyme
Meningeal fibroblasts
Pericytes
Preskeletal osteo fibroblasts
Duct mesenchyme
Preskeletal osteo chondrocytes
Dermal fibroblasts

**Extended Data Figure 12. Gene expression predictors of fate biases in progenitors**

**(a, b)** UMAP of gene expression colored by a marker gene for mesenchymal progenitors **(a)** and by cells predicted as mesenchymal progenitors **(b)**. **(c)** Scatterplot comparing gene importance for cell type prediction (X-axis) versus prediction of clone2vec coordinates (Y-axis). **(d)** Left: Bar plot showing the mean absolute SHAP values of the top 25 genes contributing to clone2vec coordinate prediction. Right: The same genes, annotated by their importance for cell type prediction. **(e–f)** Analogous to panels **(b–d)**, but for neuronal progenitors.

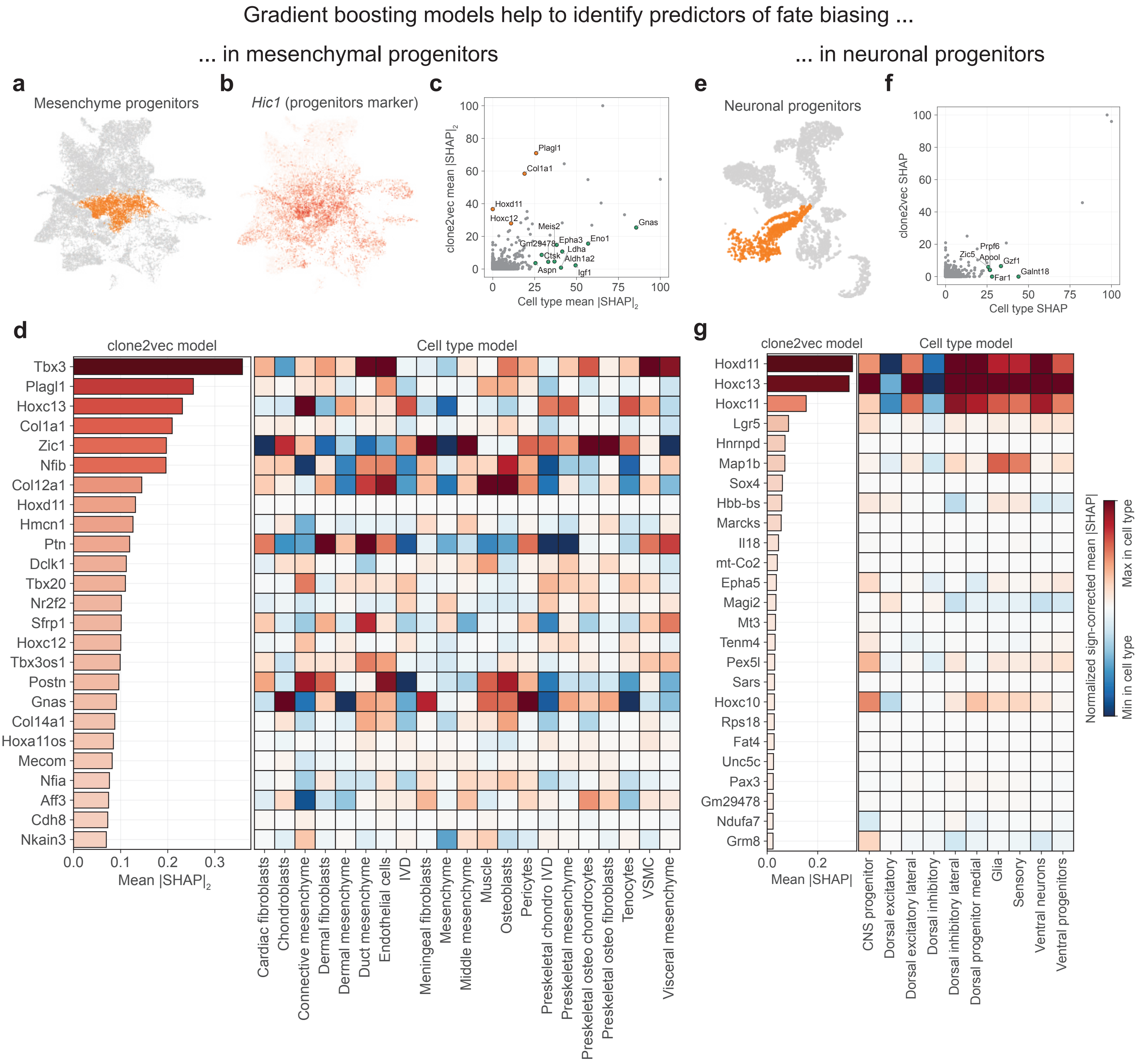
Gradient boosting models help to identify predictors of fate biasing ...
... in mesenchymal progenitors
... in neuronal progenitors
a
Mesenchyme progenitors
b
Hic1 (progenitors marker)
c
clone2vec mean |SHAP|2
Cell type mean |SHAP|2
Plagl1
Col1a1
Hoxd11
Hoxc12
Meis2
Gnas
Eno1
Epha3
Gm29478
Ldha
Ctsk
Aldh1a2
Aspn
Igf1
e
Neuronal progenitors
f
clone2vec SHAP
Cell type SHAP
Prpf6
Zic5
Apool
Gzf1
Far1
Galnt18
d
clone2vec model
Cell type model
Tbx3
Plagl1
Hoxc13
Col1a1
Zic1
Nfib
Col12a1
Hoxd11
Hmcn1
Ptn
Dclk1
Tbx20
Nr2f2
Sfrp1
Hoxc12
Tbx3os1
Postn
Gnas
Col14a1
Hoxa11os
Mecom
Nfia
Aff3
Cdh8
Nkain3
Mean |SHAP|2
Cardiac fibroblasts
Chondroblasts
Connective mesenchyme
Dermal fibroblasts
Dermal mesenchyme
Duct mesenchyme
Endothelial cells
IVD
Meningeal fibroblasts
Mesenchyme
Middle mesenchyme
Muscle
Osteoblasts
Pericytes
Preskeletal chondro IVD
Preskeletal mesenchyme
Preskeletal osteo chondrocytes
Preskeletal osteo fibroblasts
Tenocytes
VSMC
Visceral mesenchyme
g
clone2vec model
Cell type model
Hoxd11
Hoxc13
Hoxc11
Lgr5
Hnrnpd
Map1b
Sox4
Hbb-bs
Marcks
Il18
mt-Co2
Epha5
Magi2
Mt3
Tenm4
Pex5l
Sars
Hoxc10
Rps18
Fat4
Unc5c
Pax3
Gm29478
Ndufa7
Grm8
Mean |SHAP|
CNS progenitor
Dorsal excitatory
Dorsal excitatory lateral
Dorsal inhibitory
Dorsal inhibitory lateral
Dorsal progenitor medial
Glia
Sensory
Ventral neurons
Ventral progenitors
Normalized sign-corrected mean |SHAP|
Max in cell type
Min in cell type

**Extended Data Figure 13. Independent validation of NCC-MPs in tail and lower trunk mesenchyme.**

Four additional embryo samples (embryos injected at E7.5 and lower trunk/tails were harvested at E13.5-E14.5) and subject to clone2vec analysis. These are not included in the general dataset.

**(a)** Gene expression UMAP embeddings of the traced cells, color coded by the main cell types of interest (central nervous system cells, peripheral nervous system cells, mesodermal derivatives, and non-neural ectodermal derivatives).

**(b-c)** Clone2vec UMAP embeddings of the recovered clones, color coded by clonal cluster (b) and Hoxc13 expression level per clone (c). Tail mesenchyme was selected for downstream analysis in (g-m) based on specific expression of Hoxc13.

**(d-f)** Clone2vec UMAP embeddings color coded by the number of cells per clone for each cell type, including mesenchyme (d), CNS neurons (e), and neural crest-derived fates (f). Arrows point out regions of specific mesenchyme-biased clones containing crest-derived but not CNS cells.

**(g-j)** Clone2vec UMAP embeddings of the sub-selected tail mesenchyme cluster, color-coded by sub-cluster (g) or the number of cells per clone in the given fates, including mesenchyme (h), CNS neurons (i), or crest-derived cells (j). Subclusters 2 and 11, circled in (g), represent the CNS-biased NMPs versus the crest-biased NCC-MPs.

**(k-l)** Analysis of clonal fate biases in crest-biased NCC-MP clones versus CNS-biased NMP clones from the tail mesenchyme. (k-l) Bar and whisker plots showing the abundance of CNS (k) versus neural crest (l) fates in each clonal cluster. Statistical significance between the clusters was determined by a Mann-Whitney U test.

**(m)** Ternary plot of tail mesenchyme clones in the NCC-MP and NMP clusters. The position of each point is determined by the composition of the clone, where dots closer to each corner are more biased towards the labeled cell type.

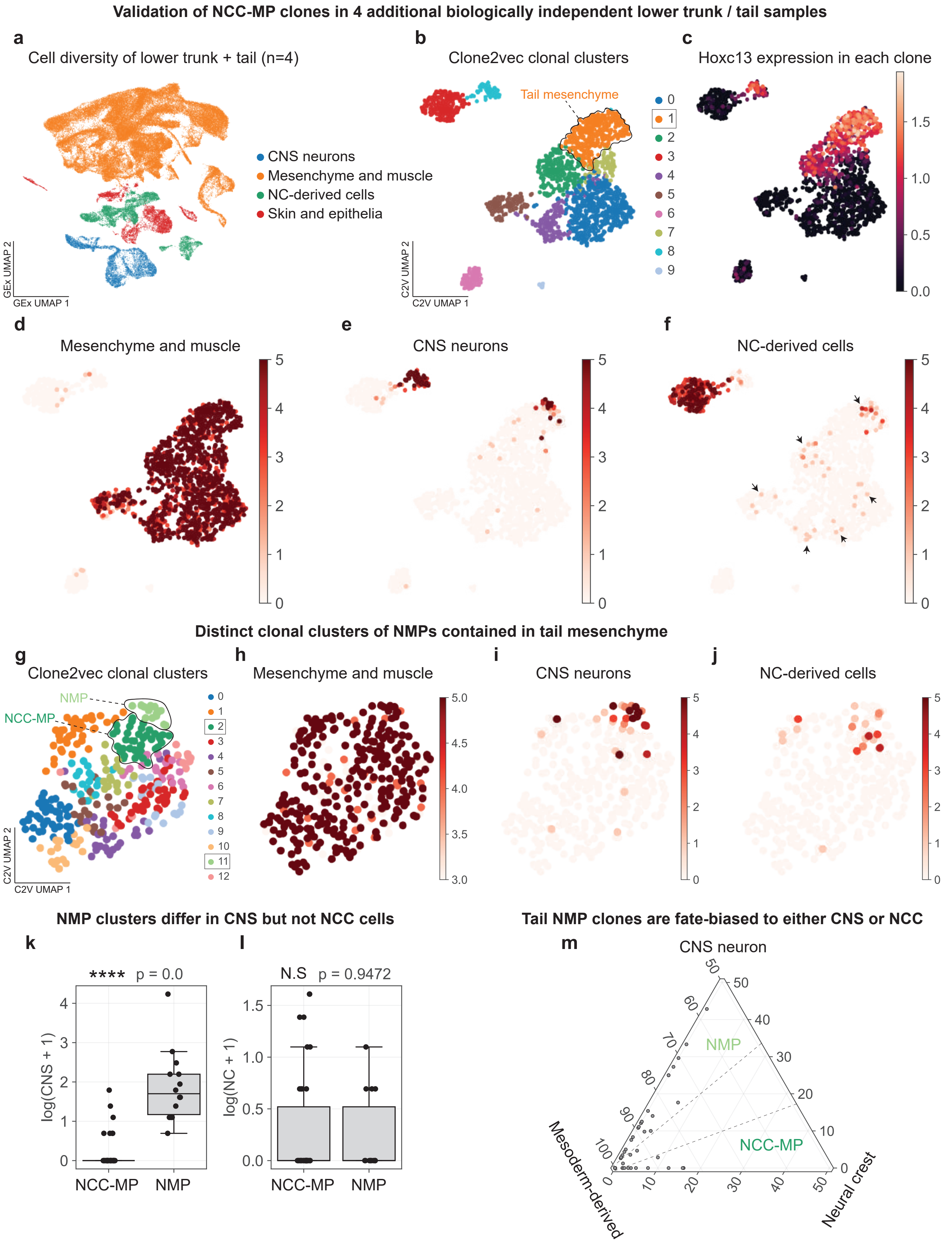

Validation of NCC-MP clones in 4 additional biologically independent lower trunk / tail samples
a
Cell diversity of lower trunk + tail (n=4)
CNS neurons
Mesenchyme and muscle
NC-derived cells
Skin and epithelia
GEx UMAP 1
GEx UMAP 2
b
Clone2vec clonal clusters
Tail mesenchyme
C2V UMAP 1
C2V UMAP 2
c
Hoxc13 expression in each clone
d
Mesenchyme and muscle
e
CNS neurons
f
NC-derived cells
Distinct clonal clusters of NMPs contained in tail mesenchyme
g
Clone2vec clonal clusters
NMP
NCC-MP
h
Mesenchyme and muscle
i
CNS neurons
j
NC-derived cells
NMP clusters differ in CNS but not NCC cells
k
**** p = 0.0
log(CNS + 1)
NCC-MP
NMP
l
N.S p = 0.9472
log(NC + 1)
Tail NMP clones are fate-biased to either CNS or NCC
m
CNS neuron
NMP
NCC-MP
Mesoderm-derived
Neural crest

**Extended Data Figure 14. Spatial validations of cell type signatures in the mouse embryo.** Feature plots showing expression of marker genes for the observed mesenchymal cell types both in gene expression space (in a cell UMAP embedding) as well as anatomical space (using publicly available Stereoseq data of E13.5 mouse embryo visualized in a sagittal section roughly through the midline). The mapped cell types include vascular smooth muscle cells **(a)**, perivascular cells **(b)**, intervertebral disk fibrocartilage cells **(c)**, osteoblasts **(d)**, meningeal fibroblasts **(e)**, chondrocytes **(f)**, mesothelial cells **(g)**, mesenchymal cells found in ducts **(h)**, and dermal mesenchymal cells **(i)**.

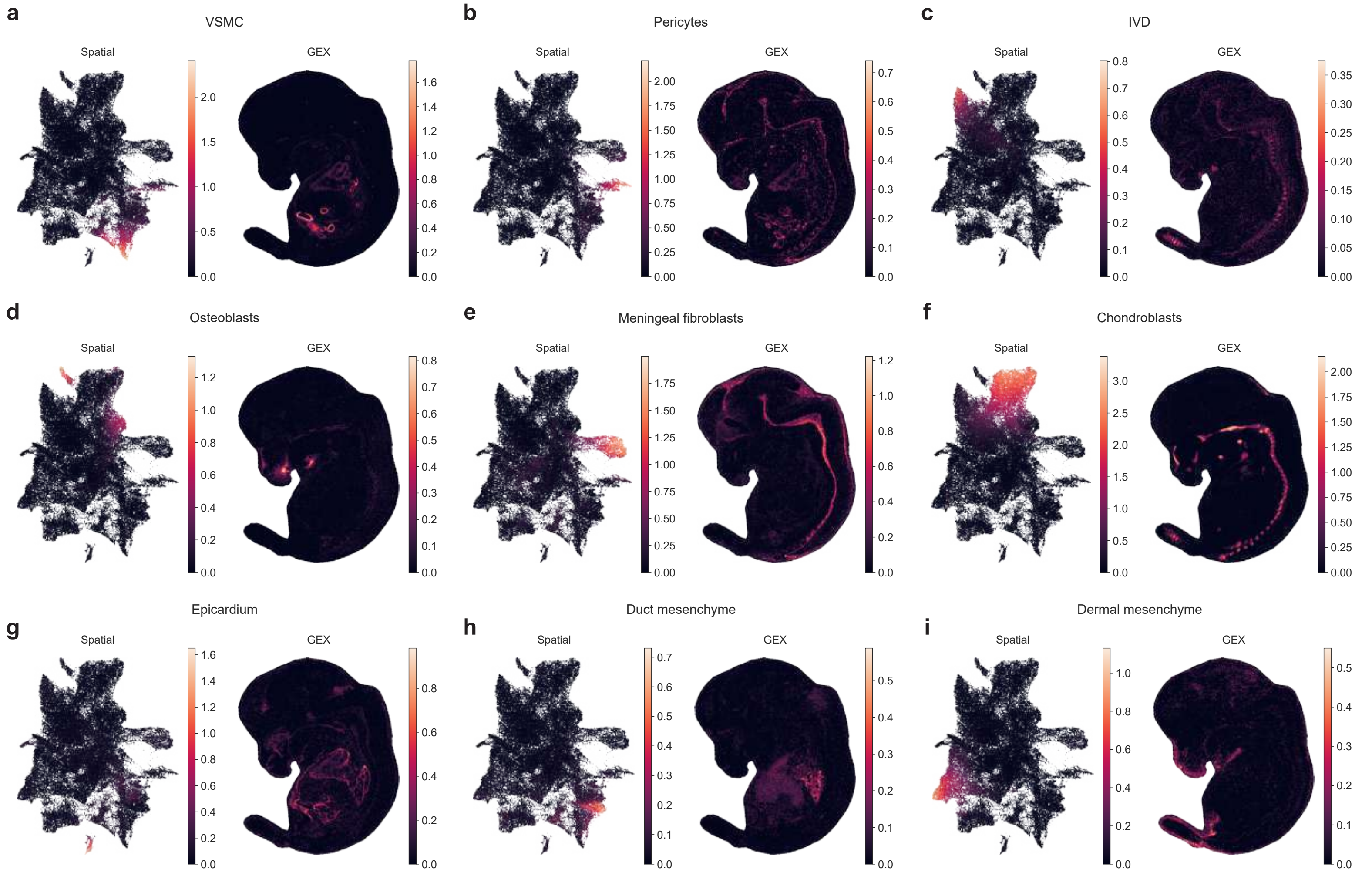
a
VSMC
Spatial
GEX
b
Pericytes
Spatial
GEX
c
IVD
Spatial
GEX
d
Osteoblasts
Spatial
GEX
e
Meningeal fibroblasts
Spatial
GEX
f
Chondroblasts
Spatial
GEX
g
Epicardium
Spatial
GEX
h
Duct mesenchyme
Spatial
GEX
i
Dermal mesenchyme
Spatial
GEX

**Extended Data Figure 15. Cartilage cell differentiation trajectory analysis 1.**

**(a-d)** Trajectory analysis of ascending and descending chondrogenesis-associated transcriptional programs common to both clonal clusters. Scatterplots indicate the strength of the program as a function of the calculated pseudotime. **(e-f)** Bone-related signature expression on a heatmap (e) and UMAPs (f). **(g)** Analysis of "early" pre-bifurcation bone-related genes (slope and significance of regression before the bifurcation points). **(h-i)** *Runx2* expression showed on UMAP for all cells from the trajectories (h) and only in the progenitor population (i). **(j)** Violin plot showing differences in Runx2 expression in progenitor population. **(k)** Regression analysis of association between Runx2 expression and pseudotime in chondrogenic and osteochondrogenic progenitors.

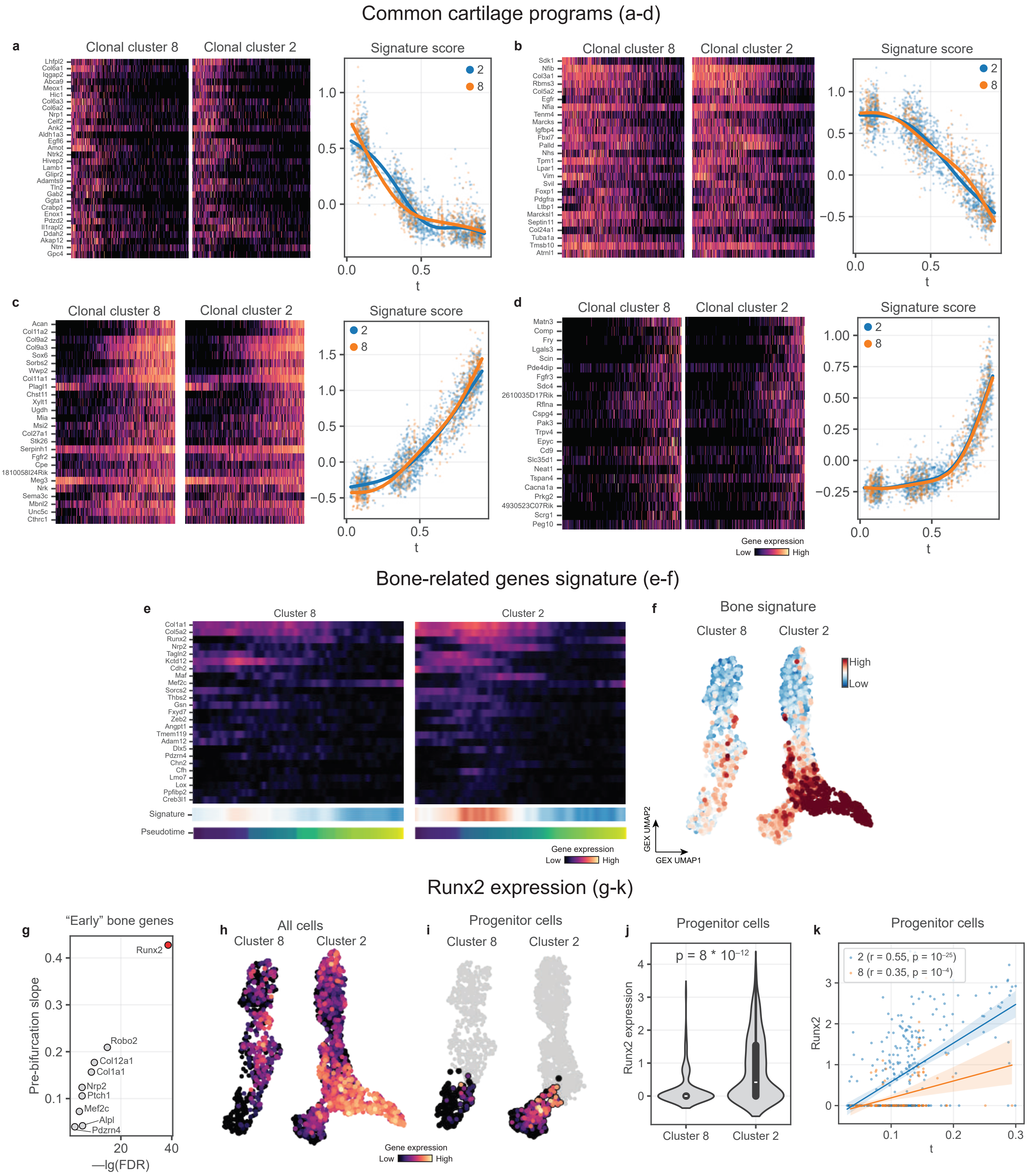

Common cartilage programs (a-d)
a
Clonal cluster 8
Clonal cluster 2
Signature score
Lhfpl2
Col6a1
Iqgap2
Abca9
Meox1
Hic1
Col6a3
Col6a2
Nrp1
Celf2
Ank2
Aldh1a3
Egfl6
Amot
Ntrk2
Hivep2
Lamb1
Glipr2
Adamts9
Tln2
Gab2
Ggta1
Crabp2
Enox1
Pdzd2
Il1rapl2
Ddah2
Akap12
Ntm
Gpc4
2
8
t
b
Clonal cluster 8
Clonal cluster 2
Signature score
Sdk1
Nfib
Col3a1
Rbms3
Col5a2
Egfr
Nfia
Tenm4
Marcks
Igfbp4
Fbxl7
Palld
Nhs
Tpm1
Lpar1
Vim
Svil
Foxp1
Pdgfra
Ltbp1
Marcksl1
Septin11
Col24a1
Tuba1a
Tmsb10
Atrnl1
c
Clonal cluster 8
Clonal cluster 2
Signature score
Acan
Col11a2
Col9a2
Col9a3
Sox6
Sorbs2
Wwp2
Col11a1
Plagl1
Chst11
Xylt1
Ugdh
Mia
Msi2
Col27a1
Stk26
Serpinh1
Fgfr2
Cpe
1810058I24Rik
Meg3
Nrk
Sema3c
Mbnl2
Unc5c
Cthrc1
d
Clonal cluster 8
Clonal cluster 2
Signature score
Matn3
Comp
Fry
Lgals3
Scin
Pde4dip
Fgfr3
Sdc4
2610035D17Rik
Rflna
Cspg4
Pak3
Trpv4
Epyc
Cd9
Slc35d1
Neat1
Tspan4
Cacna1a
Prkg2
4930523C07Rik
Scrg1
Peg10
Gene expression
Low High
Bone-related genes signature (e-f)
e
Cluster 8
Cluster 2
Col1a1
Col5a2
Runx2
Nrp2
Tagln2
Kctd12
Cdh2
Maf
Mef2c
Sorcs2
Thbs2
Gsn
Fxyd7
Zeb2
Angpt1
Tmem119
Adam12
Dlx5
Pdzrn4
Chn2
Cfh
Lmo7
Lox
Ppfibp2
Creb3l1
Signature
Pseudotime
Gene expression
Low High
f
Bone signature
Cluster 8
Cluster 2
High
Low
GEX UMAP2
GEX UMAP1
Runx2 expression (g-k)
g
“Early” bone genes
Pre-bifurcation slope
—lg(FDR)
Runx2
Robo2
Col12a1
Col1a1
Nrp2
Ptch1
Mef2c
Alpl
Pdzrn4
h
All cells
Cluster 8
Cluster 2
Gene expression
Low High
i
Progenitor cells
Cluster 8
Cluster 2
j
Progenitor cells
p = 8 * 10⁻¹²
Runx2 expression
Cluster 8
Cluster 2
k
Progenitor cells
2 (r = 0.55, p = 10⁻²⁵)
8 (r = 0.35, p = 10⁻⁴)
Runx2
t

**Extended Data Figure 16. Cartilage cell differentiation trajectory analysis 2.**

**(a-c)** Sustained early (a), transient early (b) and transient (c) gene programs that are upregulated specifically in Cluster 2 chondrocytes versus Cluster 8 chondrocytes. **(d)** Enrichment analysis for scenic-reconstructed targets of transcriptional factors within signature from (c). **(e-h)** Descending (e), transient (f) transient early (g) and ascending (h) gene programs that are upregulated specifically in Cluster 8 chondrocytes versus Cluster 2 chondrocytes. **(i)** Heatmap displaying patterns of regulon activity (top) and transcription factor expression (bottom) unique to chondrogenic clones. **(j)** Gene regulatory networks predicted within this lineage.

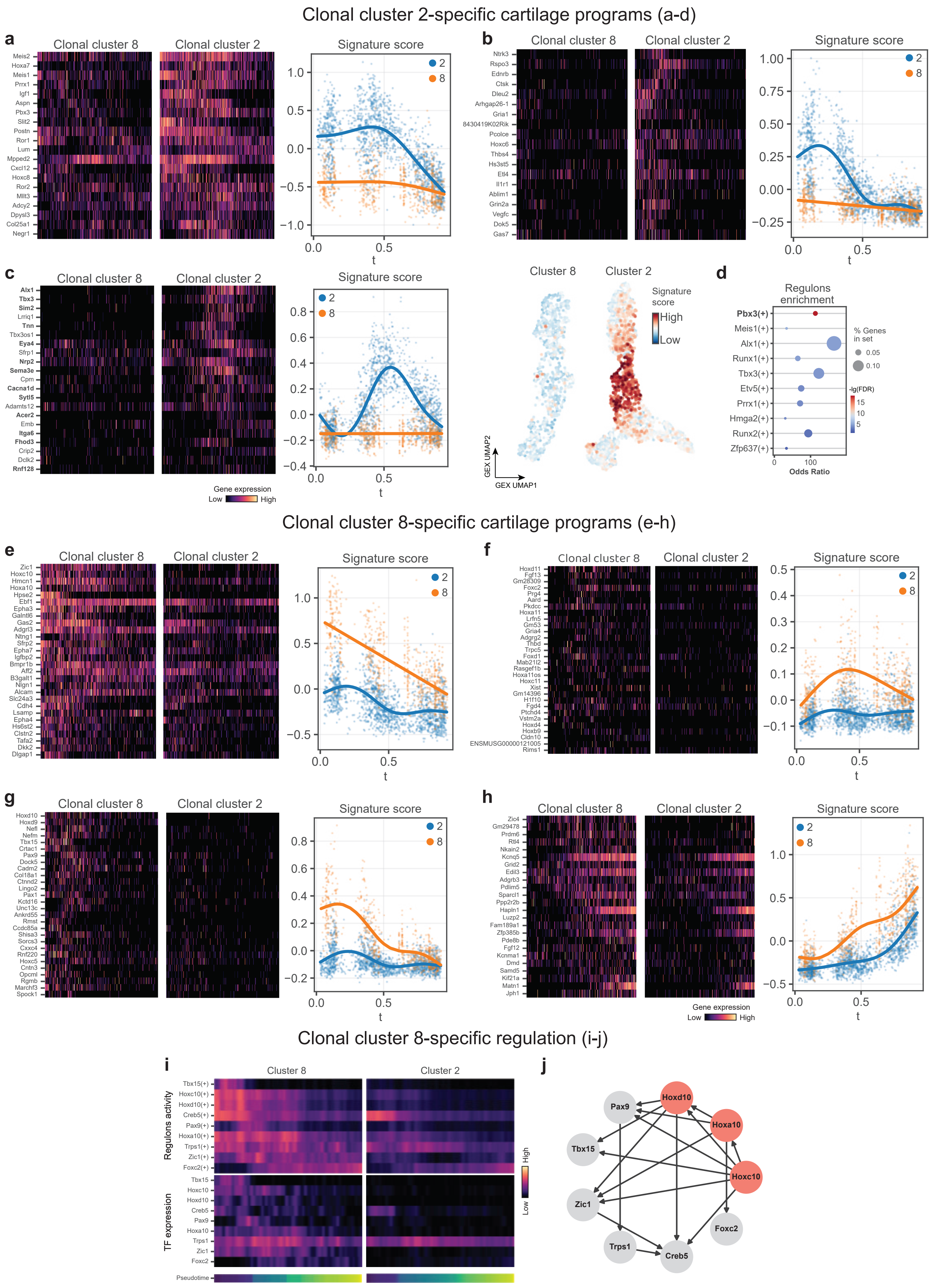

Clonal cluster 2-specific cartilage programs (a-d)
a
Clonal cluster 8
Clonal cluster 2
Signature score
Meis2
Hoxa7
Meis1
Prrx1
Igf1
Aspn
Pbx3
Slit2
Postn
Ror1
Lum
Mpped2
Cxcl12
Hoxc8
Ror2
Mllt3
Adcy2
Dpysl3
Col25a1
Negr1
t
b
Clonal cluster 8
Clonal cluster 2
Signature score
Ntrk3
Rspo3
Ednrb
Ctsk
Dleu2
Arhgap26-1
Gria1
8430419K02Rik
Pcolce
Hoxc6
Thbs4
Hs3st5
Etl4
Il1r1
Ablim1
Grin2a
Vegfc
Dok5
Gas7
t
c
Clonal cluster 8
Clonal cluster 2
Signature score
Alx1
Tbx3
Sim2
Lrriq1
Tnn
Tbx3os1
Eya4
Sfrp1
Nrp2
Sema3e
Cpm
Cacna1d
Sytl5
Adamts12
Acer2
Emb
Itga6
Fhod3
Crip2
Dclk2
Rnf128
Gene expression
Low
High
t
Cluster 8
Cluster 2
Signature score
High
Low
GEX UMAP2
GEX UMAP1
d
Regulons enrichment
Pbx3(+)
Meis1(+)
Alx1(+)
Runx1(+)
Tbx3(+)
Etv5(+)
Prrx1(+)
Hmga2(+)
Runx2(+)
Zfp637(+)
% Genes in set
0.05
0.10
-lg(FDR)
Odds Ratio
Clonal cluster 8-specific cartilage programs (e-h)
e
Clonal cluster 8
Clonal cluster 2
Signature score
Zic1
Hoxc10
Hmcn1
Hoxa10
Hpse2
Ebf1
Epha3
Galntl6
Gas2
Adgrl3
Ntng1
Sfrp2
Epha7
Igfbp2
Bmpr1b
Aff2
B3galt1
Nlgn1
Alcam
Slc24a3
Cdh4
Lsamp
Epha4
Hs6st2
Clstn2
Tafa2
Dkk2
Dlgap1
t
f
Clonal cluster 8
Clonal cluster 2
Signature score
Hoxd11
Fgf13
Gm28309
Foxc2
Prg4
Aard
Pkdcc
Hoxa11
Lrfn5
Gm53
Gria4
Adgrg2
Thbd
Trpc5
Foxd1
Mab21l2
Rasgef1b
Hoxa11os
Hoxc11
Xist
Gm14396
H1f10
Fgd4
Ptchd4
Vstm2a
Hoxd4
Hoxb9
Cldn10
ENSMUSG00000121005
Rims1
t
g
Clonal cluster 8
Clonal cluster 2
Signature score
Hoxd10
Hoxd9
Nefl
Nefm
Tbx15
Crtac1
Pax9
Dock5
Cadm2
Col18a1
Ctnnd2
Lingo2
Pax1
Kctd16
Unc13c
Ankrd55
Rmst
Ccdc85a
Shisa3
Sorcs3
Cxxc4
Rnf220
Hoxc5
Cntn3
Opcml
Rgmb
Marchf3
Spock1
t
h
Clonal cluster 8
Clonal cluster 2
Signature score
Zic4
Gm29478
Prdm6
Rtl4
Nkain2
Kcnq5
Grid2
Edil3
Adgrb3
Pdlim5
Sparcl1
Ppp2r2b
Hapln1
Luzp2
Fam189a1
Zfp385b
Pde8b
Fgf12
Kcnma1
Dmd
Samd5
Kif21a
Matn1
Jph1
Gene expression
Low
High
t
Clonal cluster 8-specific regulation (i-j)
i
Cluster 8
Cluster 2
Regulons activity
Tbx15(+)
Hoxc10(+)
Hoxd10(+)
Creb5(+)
Pax9(+)
Hoxa10(+)
Trps1(+)
Zic1(+)
Foxc2(+)
TF expression
Tbx15
Hoxc10
Hoxd10
Creb5
Pax9
Hoxa10
Trps1
Zic1
Foxc2
Pseudotime
High
Low
j
Hoxd10
Pax9
Hoxa10
Tbx15
Hoxc10
Zic1
Foxc2
Trps1
Creb5

**Extended Data Figure 17. Characterization of clonal clusters resulting from perturb-TREX.**

**(a-b)** Clone2vec embedding of the combined control and perturbed clones from the craniofacial region, color coded by the discovered clone types (a) and perturbations (b). **(c-d)** Bar chart of the clone types in the combined control and perturbation datasets from trunk (a) and craniofacial region (b), indicating the observed enrichment of each perturbation within each clone type. **(e-f)** Dot plot showing the distribution of cell types across each clone type in the combined control and perturbation dataset from trunk (e) and craniofacial (f) regions. Dot size indicates the fraction of clones of a particular clone type that contain cells within a particular fate, and the color of the dot shows the average number of cells within that fate per clone within that clone type.

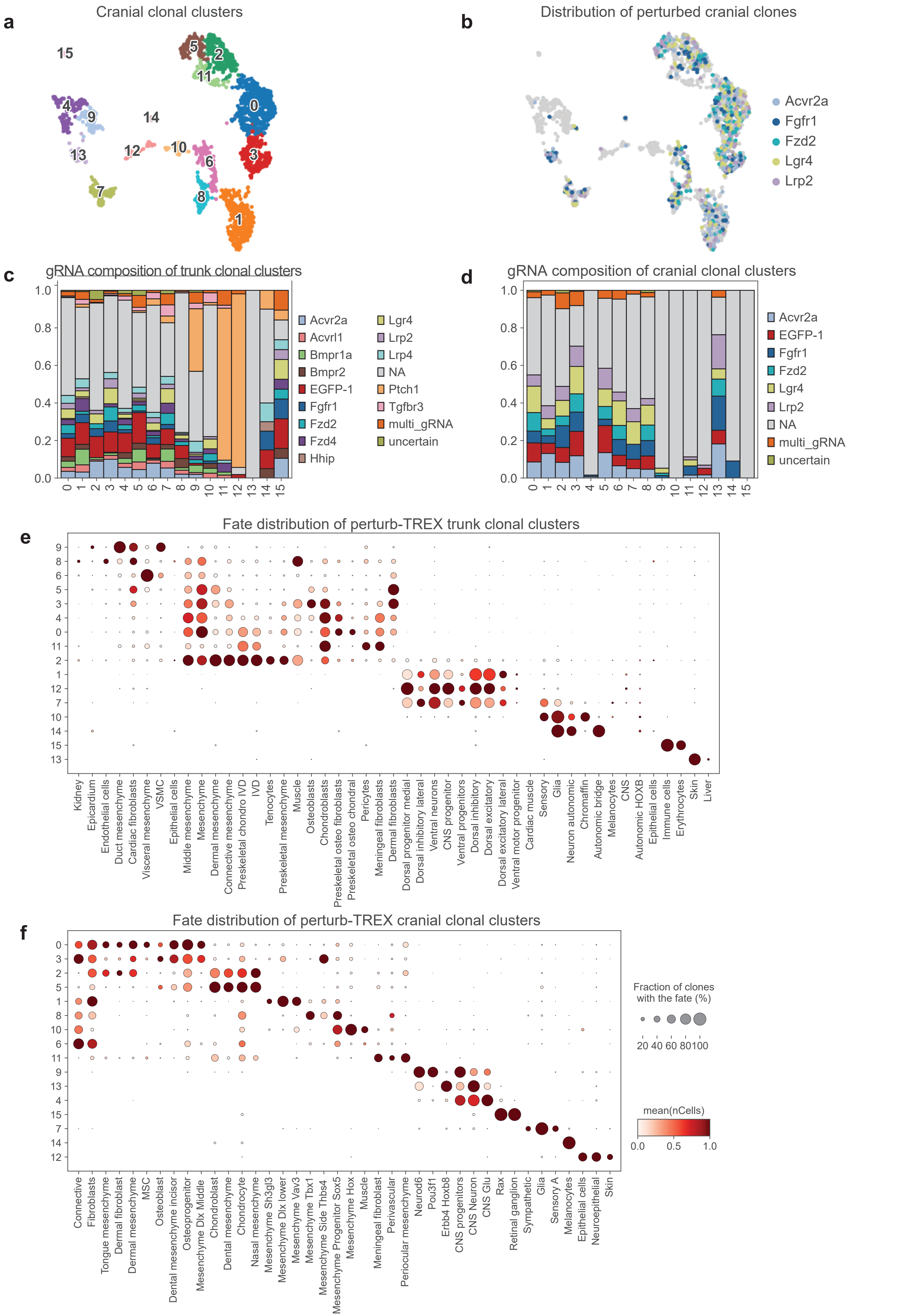
a
Cranial clonal clusters
15
5
2
11
0
4
9
14
12
10
6
3
13
8
7
1
b
Distribution of perturbed cranial clones
Acvr2a
Fgfr1
Fzd2
Lgr4
Lrp2
c
gRNA composition of trunk clonal clusters
1.0
0.8
0.6
0.4
0.2
0.0
0 1 2 3 4 5 6 7 8 9 10 11 12 13 14 15
Acvr2a
Acvrl1
Bmpr1a
Bmpr2
EGFP-1
Fgfr1
Fzd2
Fzd4
Hhip
Lgr4
Lrp2
Lrp4
NA
Ptch1
Tgfbr3
multi_gRNA
uncertain
d
gRNA composition of cranial clonal clusters
1.0
0.8
0.6
0.4
0.2
0.0
0 1 2 3 4 5 6 7 8 9 10 11 12 13 14 15
Acvr2a
EGFP-1
Fgfr1
Fzd2
Lgr4
Lrp2
NA
multi_gRNA
uncertain
e
Fate distribution of perturb-TREX trunk clonal clusters
9
8
6
5
3
4
0
11
2
1
12
7
10
14
15
13
Kidney
Epicardium
Endothelial cells
Duct mesenchyme
Cardiac fibroblasts
Visceral mesenchyme
VSMC
Epithelial cells
Middle mesenchyme
Mesenchyme
Dermal mesenchyme
Connective mesenchyme
Preskeletal chondro IVD
IVD
Tenocytes
Preskeletal mesenchyme
Muscle
Osteoblasts
Chondroblasts
Preskeletal osteo fibroblasts
Preskeletal osteo chondral
Pericytes
Meningeal fibroblasts
Dermal fibroblasts
Dorsal progenitor medial
Dorsal inhibitory lateral
Ventral neurons
CNS progenitor
Ventral progenitors
Dorsal inhibitory
Dorsal excitatory
Dorsal excitatory lateral
Ventral motor progenitor
Cardiac muscle
Sensory
Glia
Neuron autonomic
Chromaffin
Autonomic bridge
Melanocytes
CNS
Autonomic HOXB
Epithelial cells
Immune cells
Erythrocytes
Skin
Liver
f
Fate distribution of perturb-TREX cranial clonal clusters
0
3
2
5
1
8
10
6
11
9
13
4
15
7
14
12
Connective
Fibroblasts
Tongue mesenchyme
Dermal fibroblast
Dermal mesenchyme
MSC
Osteoblast
Dental mesenchyme incisor
Osteoprogenitor
Mesenchyme Dlx Middle
Chondroblast
Dental mesenchyme
Chondrocyte
Nasal mesenchyme
Mesenchyme Sh3gl3
Mesenchyme Dlx lower
Mesenchyme Vav3
Mesenchyme Tbx1
Mesenchyme Side Thbs4
Mesenchyme Progenitor Sox5
Mesenchyme Hox
Muscle
Meningeal fibroblast
Perivascular
Periocular mesenchyme
Neurod6
Pou3f1
Erbb4 Hoxb8
CNS progenitors
CNS Neuron
CNS Glu
Rax
Retinal ganglion
Sympathetic
Glia
Sensory A
Melanocytes
Epithelial cells
Neuroepithelial
Skin
Fraction of clones with the fate (%)
20 40 60 80100
mean(nCells)
0.0 0.5 1.0
